# The preclinical cardiac phenotype of the DE50-MD dog model of Duchenne muscular dystrophy

**DOI:** 10.1101/2025.02.20.638615

**Authors:** Julia Sargent, Virginia Luis Fuentes, Rebecca Terry, Dominique O. Riddell, Rachel CM Harron, Dominic J. Wells, Richard J. Piercy

## Abstract

Cardiomyopathy is the leading cause of death in the X-linked disorder, Duchenne Muscular Dystrophy (DMD) yet optimal management strategies remain undetermined. Advances in the search for novel DMD treatments, particularly at cell and molecular levels, rely heavily on the use of translational animal models. It is crucial that these models faithfully recapitulate the human clinical phenotype to best expedite the development of promising treatments.

We sought comprehensively to describe the cardiac phenotype of DE50-MD dogs, a novel dystrophin-deficient model that harbours a mutation within the principal DMD mutational hotspot. Cardiac magnetic resonance imaging and echocardiographic studies were performed at approximately 12-week intervals in male, 3- to 18-month-old DE50-MD (n=17) and age-matched littermates, wild type (WT, n=14) dogs. Late gadolinium enhancement (LGE) imaging was performed in a subpopulation of DE50-MD (n=10) and WT (n=11) dogs aged 9 to 18 months. The DE50-MD dogs had smaller left ventricular (LV) mass and LV dimensions than WT dogs. While global ventricular systolic function was preserved, DE50-MD dogs showed early differences in strain and strain rate parameters. Only DE50-MD dogs demonstrated LGE (3/8 dogs studied at 18 months); the subepicardial to transmural, mid-to-basal LV LGE distribution resembling that of DMD patients and of other dystrophic dog models. Histopathological assessment confirmed that LGE corresponded to fibrofatty myocardial scarring, as described in DMD patients and other canine models of dystrophin-deficient cardiomyopathy. The DE50-MD early preclinical cardiac phenotype shares key features of DMD cardiomyopathy prior to onset of global LV systolic dysfunction. Their disproportionately low LV volume to mass supports possible combined physiological hypotrophy and tonic contraction in affected animals.

## Introduction

The X-linked recessive childhood disorder, Duchenne muscular Dystrophy (DMD) is a fatal, progressive, muscle-wasting disease affecting approximately one in ∼5000-6000 new-born males.[1, 2]. Affected boys lack dystrophin, an integral ultrastructural protein in skeletal, smooth and cardiac muscles. The resultant progressive skeletal and cardiac muscle degeneration culminates in loss of ambulation, respiratory failure and cardiomyopathy; the latter in virtually all cases.[3] Myocardial fibrofatty replacement, beginning in the basal and mid left ventricular free wall, ultimately causes widespread myocardial dysfunction,[4] characterised by advancing ventricular systolic dysfunction and chamber dilation, i.e. a dilated cardiomyopathy (DCM) phenotype.[5, 6] Cardiac imaging with transthoracic echocardiography (TTE) and cardiac magnetic resonance imaging (CMR) are paramount, both to guide clinical decision-making and to stratify disease for objective end-points in clinical trials.[7–10]

Supportive interventions for the neuromuscular and respiratory complications of DMD, (largely through the use of palliative steroid therapy, spinal corrective surgery and mechanical ventilator assistance)[11]) have extended the predicted life expectancy for affected individuals to a median survival time into their late twenties to early thirties.[12, 13] Cardiovascular complications have become the leading cause (20-40%) of deaths in DMD.[3, 14–17] The progression and extent of cardiomyopathy varies amongst patients but clinical manifestations include onset of congestive heart failure symptoms, arrhythmia and sudden cardiac death.[5, 18] Despite its clinical significance, the pathophysiology of DMD cardiomyopathy remains incompletely understood and there is a lack of consensus regarding optimal treatment strategies. There is a renewed urgency for research focused on the search for early therapeutic targets for DMD cardiomyopathy and consequently, a need to improve our understanding of the pathophysiology behind disease progression.

Translational animal models of DMD provide a platform with which to explore mechanistic insights for the disease and to test novel treatments, (in particular, molecular and cell-based techniques). Given the impact of cardiac disease on morbidity and mortality in DMD patients, it is essential that such models closely recapitulate the human DMD cardiac phenotype. Arguably, the most extensively studied translational model is the mdx mouse, which lacks dystrophin expression due to a nonsense point mutation in exon 23 in the dystrophin gene.[19] Despite being a genetic homologue of DMD, the natural progression of the mdx model is notably different to that of DMD, demonstrating only a mild clinical phenotype with a near normal life expectancy (reduced by only 17-19%).[20] Likewise, mdx cardiomyopathy progression does not fully emulate that of DMD as global myocardial function is typically preserved until late in life, (12 months and over)[21–23], and the histological distribution of fibrotic lesions is dissimilar to those of DMD patients, (demonstrating preferential RV and septal involvement).[21, 24]

Large animal models bridge the translational gap between small animal models and people so are crucial to the drug discovery process. The Golden Retriever with Muscular Dystrophy, (GRMD; or ‘Canine X-linked muscular dystrophy’ (CXMD)) models have been extensively characterised: they exhibit a clinical phenotype that closely resembles the cardiac and skeletal muscle progression of DMD.[25–27]. The GRMD cardiomyopathy has recently been comprehensively described in a colony maintained at Texas A&M University.[28, 29] Global systolic dysfunction reportedly occurs between 30 and 45 months of age, preceded by echocardiographic evidence of myocardial deformation (circumferential strain) derangements and myocardial fibrosis, detectable using contrast (late gadolinium) enhancement (LGE) imaging with cardiac magnetic resonance (CMR).[29] An exciting recent development is the discovery of a spontaneous mutation in the Cavalier King Charles Spaniel (CKCS) breed. Unlike the exon 6 mutation in the GRMD model, this missense point mutation of the 5’donor splice site in intron 50 results in deletion of exon 50 from DMD gene transcripts. It is therefore located in the middle of the major hot spot region for mutations in DMD patients. [30] This is particularly pertinent to the substantial work underway to develop genetic repair or editing therapies and exon skipping approaches, which aim to achieve dystrophin re-expression in affected individuals. Indeed, the DE50-MD mutation is located within the major human DMD mutational hotspot and targeted approaches tested in this model are applicable to the highest proportion of DMD human patients.[31]

Since the first identification and genotyping of the proband CKCS dog at the Royal Veterinary College (RVC) in London in 2010,[30] a colony has been established through outbreeding the offspring of the proband’s dam, (CKCS-Bichon Frisé crossbred dog) onto a Beagle background (RCC strain, Marshall Bioresources) [32]. The relatively small size of these animals and the mutation site make this model highly applicable for translational research. Extensive phenotypic characterisation using magnetic resonance imaging, functional muscle assessments and histopathological studies have demonstrated that the novel model of DMD, named the Delta,(D)E50-MD dog faithfully recapitulates many of the important features of the skeletal muscle phenotype with age, [33–36]but the cardiac phenotype of the dogs is yet to be described.

In this study we report the natural history of the preclinical cardiac phenotype in DE50-MD dogs. Our goal was to provide a comprehensive description of the cardiac structure and function of young DE50-MD male dogs through serial assessment with conventional TTE and CMR techniques and comparison to age-matched WT littermate control male dogs. We developed protocols using contrast-enhanced CMR to explore for evidence of early myocardial scarring in a subpopulation of young DE50-MD dogs, our aim being to compare our findings to the cardiac phenotype of the preclinical DMD cardiomyopathy described in people and in the GRMD model.

## Materials and methods

### Study population

All dogs (*Canis familiaris*) were recruited from the DE50-MD colony housed at the RVC, UK, and cared for in accordance with the guidelines stipulated within two successive project licences, (70/7515 and P9A1D1D6E) assigned under the UK’s Animal (Scientific Procedure’s) Act, (A(SP)A) 1986 and approved by the RVC’s Animal Welfare Ethical Review Body. The dogs were direct relatives of the previously reported CKCS proband dog with the DE50-MD mutation.[30, 32] Puppies were genotyped using a previously validated polymerase chain reaction assay conducted on products amplified from cheek swab-derived DNA within 7 days from birth and were identified by digital microchip. Male DE50-MD dogs and male littermate wild type, (WT) controls were followed from approximately 12 weeks of age until the 18-month endpoint, or until the clinical phenotype reached a humane endpoint stipulated by the Home Office A(SP)A licence. Pre-determined endpoints included dehydration (unresolved by short-term fluid treatment), lethargy/motor dysfunction, weight loss/dysphagia, dyspnoea, listless behaviour/demeanour, or heart failure, as noted during daily observations by trained animal technicians and Named Veterinary Surgeons. The decision to remove animals from the study was ultimately made by the study director, Named Veterinary Surgeons and the Named Animal Care and Welfare Officer.

Serial cardiac assessments using TTE and CMR were scheduled with an intended study interval of 12 weeks. At the end of the 18-month study period, 5/17 DE50-MD and 6/14 WT dogs were recruited into a separate, ongoing longitudinal study, reported separately. All remaining DE50-MD dogs underwent humane euthanasia using an intravenous injection of sodium pentobarbital (250 mg/kg, Dolethal, Covetrus) via a preplaced catheter. Four of the 14 WT dogs included in this study also underwent planned euthanasia at the18-month endpoint of the study period (WT-G2, WT-J1, WT-K4 and WT-M2), having also been included in separate longitudinal studies that conducted in parallel.[33, 34, 36, 37] The remaining 10 WT dogs were rehomed.

### Transthoracic echocardiographic assessment

All TTE examinations were performed by a cardiology diplomate of the American College of Veterinary Internal Medicine using the same ultrasound machine (Vivid E9, General Electric Medical Systems Ultrasound, Hatfield, UK). All studies were analysed offline by that same cardiologist using proprietary software (Echopac, General Electric Medical Systems Ultrasound, Hatfield, UK.). Measurements were recorded as averages from at least 5 cardiac cycles. All standard images and measurements were collected according to the recommendations of the American Society of Echocardiography (Supplementary information Table 1).[38] Echocardiographic examinations were performed without sedation within 7 days of CMR examinations. Where TTE assessments were performed after CMR, examinations took place at least 48 hours after recovery from general anaesthesia. All dogs underwent a habituation process in order to minimise stress.

**Table 1:** Cardiac magnetic resonance imaging: Typical image acquisition parameters. Philips 1.5T Intera MRI protocols. B-FFE: balanced Fast Field echo (balanced steady state free precession imaging). IR TFE: inversion recovery with Turbo Fast gradient Echo imaging. FA flip angle TR: Repetition time. TE Echo time.

| Image Obtained | Sequence | TE (ms) | TR(ms) | No. of slices | FA (°) | Field of view (mm<sup>2</sup>) | Matrix | Voxel | Approx. scan time (min) |
| --- | --- | --- | --- | --- | --- | --- | --- | --- | --- |
| Short axis bright blood cine stack | B-FFE | 2.2 | 3.2 | 16 | 60 | 137x137x80 | 80x77 | 1.7x1.8x5 | 2.31 |
| Inversion recovery gradient echo images for LGE imaging | IR-TFE | 3.5 | 7.1 | 12 | 20 | 140x140x60 | 116x102 | 1.2x1.4x5.0 | 2.15 |

Conscious dogs were gently restrained in lateral recumbency. Standard echocardiographic views were obtained from the right parasternal, subcostal, left apical and left cranial parasternal positions using a 6 or 12 MHz phased-array transducer with continuous ECG recording. For a detailed description of echocardiographic measurements and allometric scaling see Supplementary Table 1. Briefly, LV volumes, (LVEDVI, LVESVI: LV end-diastolic and LV end-systolic volume indexed to body surface area (BSA)), LV length, left atrial (LA) dimensions and aortic (Ao) diameter were obtained by standard 2D measurements. Left ventricular internal dimensions (LV diastolic internal diameters normalised with allometric scaling according to bodyweight: LVIDDN and LVIDSN. LV diastolic and systolic dimensions indexed to Ao: LVIDd:Ao and LVIDs:Ao) and tricuspid annular plane systolic excursion (TAPSE, normalised to aortic diameter obtained from the left cranial view, using allometric scaling according to bodyweight[39]) were obtained using M Mode recordings. Relative wall thickness, (RWT) was calculated as the average of the septal and free wall diastolic diameter divided by LV diastolic diameter, obtained from M Mode recordings. Measurements of LV systolic function were made as follows: Ejection Fraction (EF, using a monoplane method from the right parasternal 4 chamber long axis view), Fractional shortening (FS), Cardiac Index (CI), Stroke volume Index (SVI), and Sphericity Index (SI) (calculated as described in Supplementary information Table 1).

### Colour tissue Doppler imaging

Tissue Doppler imaging (TDI) was performed to evaluate LV myocardial velocities using a previously described method.[40, 41] A detailed description of the TDI protocols is provided in the supplementary information. Endocardial and epicardial radial LV myocardial velocities were recorded from the inferolateral wall from standard right parasternal short axis echocardiographic views at the level of the mid-ventricle (papillary muscle level). Radial myocardial velocity gradients (MVGs) were defined as the difference between endocardial and epicardial velocities at the peak of systole, early and active diastole.[42] Longitudinal LV s’, e’ and a’ velocities and RV systolic velocity (RS’) were measured from the left apical 4 chamber view using pulsed-wave spectral Doppler. [40] Values for RS’ were adjusted for body size using the equation RS/BW^0.233^.[39]

#### Speckle Tracking Echocardiography

Strain parameters were derived using speckle tracking echocardiography (STE) every 3 months between 9 and 18 months of age. Studies were performed by the same observer using the same machine and transducer as for the conventional TTE assessment. [43][44][45] Image analysis was performed offline using the proprietary software listed above to obtain peak systolic strain (S) and strain rate during systole (systolic strain rate: SSR), early diastole (early diastolic strain rate: ESR) and late diastole (active diastolic strain rate: ASR), as defined by the concurrent ECG recordings. Peak systolic and early and late diastolic values were defined as the maximal deflections of the respective curves during the systolic and diastolic phases. A detailed description of the STE protocols is provided in the supplementary information.

Values for peak strain were obtained from endomyocardial, midmyocardial and epimyocardial layers and averaged to obtain the peak systolic strain value for each individual wall segment. Average peak strain and strain rate parameters were calculated by averaging the results from the 6 individual segments from 3 cycles. For longitudinal parameters the strain values for the interventricular septum and left ventricular anterolateral (free wall) were calculated from the apical, mid and basal peak longitudinal strain value for each wall.

### Cardiac magnetic resonance imaging

CMR imaging assessments were performed under general anaesthesia which was induced with intravenous propofol (4-6mg/kg, Propoflo, Zoetis) and maintained with sevoflurane, (SevoFlo, Zoetis) following premedication with intravenous methadone (0.2mg/kg, Synthadon, Animalcare). All dogs received maintenance intravenous fluid therapy during general anaesthesia with isotonic compound sodium lactate (Aquapharm11, Animalcare). Each dog was positioned in left lateral recumbency and intermittent positive pressure ventilation was maintained throughout using a mechanical ventilator. CMR imaging was performed using 1·5 Tesla Magnetic resonance imaging (MRI) machine, (Intera Pulsar System, Philips Medical Systems, Surrey) and a flexible, sensitivity-encoding, phased array surface coil (SENSETM Flex M). A MRI compatible vectorcardiogram gating device was used to derive electromotive signals in orthogonal spatial planes, to provide R peak triggering for retrospective gating. This was to synchronise image acquisition within the cardiac cycle, while minimising the appearance of artifacts generated by the magnetohydrodynamic effects during blood flow. Typical acquisition protocols used for CMR studies are summarised in table 1 and a detailed description of CMR protocols is provided in Supplementary information. Standard CMR imaging planes were prescribed, including 2-, 3- and 4-chamber LV vertical long axis (VLA) cine loops and a series of short axis images of the LV and RV.[46]

CMR images were viewed using open source Osirix software (http:// www.osirix-viewer.com). Ventricular end-systolic and end-diastolic volumes, and LV mass were obtained using the disc summation method (see supplementary information). The LV mass was calculated both in systole and diastole by subtracting the endocardial volume from the epicardial volume and multiplying the result by the myocardial mass density (1.05g/ml).[43, 44]. Ventricular volumes were used to calculate LV and RV stroke volumes and LV and RV ejections fraction (see supplementary information). Ventricular volumes and LV mass were normalised to patient size by indexing calculated values to body surface area (BSA) to yield the following variables: indexed LV end diastolic volume (LVEDVI CMR) and indexed stroke volume (LVESVI CMR). Additional exploratory analysis was performed by indexing the LV mass both to the 5th lumbar vertebra (L5) length and to femur length (as measured from the head of the femur to the medial femoral condyle from 3D multiplanar reconstructions). The LV remodelling index (LVRI) was calculated by dividing LV mass by end-diastolic volume.

### Late gadolinium enhancement imaging

Late gadolinium enhancement (LGE) imaging was performed in a subpopulation of dogs between 6 and 15 minutes after intravenous administration of 0.1mol/kg of gadolinium (Gd-DTPA (Gadovist, Bayer)). Images were obtained using inversion recovery prepared turbo fast echo pulse sequences, with optimal inversion time based on visual inspection of the Look-Locker sequence as the time taken effectively to null the normal myocardium. Images were acquired in the 2, 3 and 4 chamber vertical long axes and in SA at the mid, apical and basal myocardial segments. Additional PSIR images were repeated in the same short axis views to ensure optimal imaging of the ventricular myocardium. Images were subsequently manually reviewed using the Osirix platform for evidence of bright signal intensity associated with LGE. Myocardial segments were classified according to the American Heart Association guidelines for cardiac imaging.[45]

### Histopathology

Hearts were collected from dogs that underwent euthanasia at 18 months. Hearts were immediately removed, flushed to remove blood clots and fixed whole in 10% neutral buffered formalin. After fixation, the ventricles of each heart were sectioned in short axis transverse sections (approximately 10 mm thick) from base to apex. Basal, mid and apical sections were sectioned to approximate the imaging planes obtained during CMR assessments, placed into tissue cassettes. When present, myocardial samples were obtained from the LV wall from regions corresponding to areas of LGE positivity. Paraffin embedded tissue blocks were processed routinely and sectioned and stained with haematoxylin and eosin (H&E) and Masson’s trichrome stains. All slides were reviewed (blind to animal genotype and clinical/imaging data) by a single, diplomate of the American College of Veterinary Pathologists, with experience in laboratory animal (including Beagle) pathology and with the mdx murine model.

### Cardiac Biomarkers

Cardiac biomarkers were measured from blood samples collected from DE50-MD and WT dogs as part of a parallel study evaluating blood-borne musculoskeletal disease biomarkers in the DE50-MD model.[46] This cohort included 3 DE50-MD dogs that were not included in the imaging protocols. Approximately 2.5mls of blood was taken, and this volume was split equally between lithium heparin and plain collection tubes to obtain plasma and serum respectively. Blood samples were spun in a centrifuge at 500 x g for 10 minutes at 4°C. Serum was extracted from the blood samples, aliquoted into 1.5ml Eppendorf tubes and frozen at -80°C until required for analysis. Samples were sent to an external commercial laboratory (IDEXX Laboratories, Ludwigsburg, Germany) for quantification of cardiac troponin I (cTnI) using a high sensitivity assay (healthy range <70pg/ml) and N-Terminal pro brain natriuretic peptide (NT proBNP). Values below the lower limit of detection of the NT-proBNP assay (<250pmol/l) were documented as 250pmol/l.

### Statistical analysis

Statistical analysis was performed using commercially available software (Prism 7.70e, Graphpad and SPSS statistics, V23. IBM). Statistical significance was set at p<0.05. Differences in repeated measures were analysed using a linear mixed model, (LMM) with a compound symmetry correlation structure, Fisher’s LSD post-hoc comparisons, entering age, genotype and their interaction as fixed effects and dog as a random effect. The relationship between left ventricular dimensions and function were evaluated using Pearson’s correlation coefficient. The significance of the influence of RV parameters on LV parameters was evaluated in LMM, entering each dog’s ID as a random effect to account for repeated measures, and the LV parameter as a covariate in the model. For the latter, each genotype was considered in separate analyses. Normality of the computed models’ residuals were tested by visual inspection of residual histograms and quantile-quantile plots and using the Kolmogorov-Smirnov test for normality. Where criteria for normality were not achieved, the data for the dependent variable were logarithmically transformed and re-entered into the model.

## Results

### Study Population: TTE and CMR assessments

The study population is summarised in supplementary tables 2 and 3. Six of the earliest recruited DE50-MD dogs did not reach the intended 18-month end point of the study due to progression of their clinical phenotype to a humane endpoint stipulated in our Home Office licence. In five dogs the reason for euthanasia was onset of dysphagia, which is a common complication in other dystrophin-deficient canine models secondary to reduced jaw opening, macroglossia and pharyngo-oesophageal dysfunction.[47, 48] No other humane endpoints were reached. A sixth dog (DE50-G4) was euthanised due to developmental elbow dysplasia at 11 months of age, believed to be independent of the DMD phenotype. To account for these early withdrawals, additional dogs were recruited but not all were studied from 3 months of age. Of the final population enrolled, eight DE50-MD dogs did not reach the endpoint: 2 DE50-MD dogs that were euthanised after the 3-month data collection point, 3 DE50-MD dogs after the 6 months data collection point and 1 DE50-MD dog each at the 9-month, 12-month and 15-month data collection point respectively. In addition to these dogs, a single WT dog was euthanised after the 12-month data collection point due to development of spinal pain that was attributed to steroid responsive meningitis-arteritis, (a condition known to afflict Beagles).[49] The final number of dogs available for analyses was therefore 14 WT and 17 DE50-MD dogs, but not all dogs were studied at every time point. The dogs that were euthanased early because they prematurely reached a humane endpoint did not demonstrate significant differences in LV or RV function or linear measures of left chamber size at the time of their final assessment when compared with DE50-MD dogs aged 18 months that completed study (EF; 66.3%± 6.9 vs 67.4±3.4, p=0.681, FS; 43.8±7.0 vs 39.1 ±5.6, p=0.142), LVIDDN; 1.21±0.08 vs 1.26 ± 0.10, p=0.325, La:Ao; 1.24±0.11 vs 1.26 ±0.06, p=0.728 or TAPSEn; 4.86±1.13 vs 5.6±0.963, p=0.173.)

### Structural assessments

Complete conscious TTE assessments could not be completed in a small number of WT dogs due to poor tolerance of conscious restraint. Conscious TTE assessments could be completed in all DE50-MD dogs, however these dogs tended to have narrow and deep chest conformations with prominent sternebrae that somewhat restricted the acoustic window for collection of left apical views.

### Left cardiac chambers and dimensions: (Table 2, figures 1 and 2)

DE50-MD dogs had a lower body mass than WT dogs, becoming significant from 6 months of age, (Supplementary table 2, inset; p<0.001). Despite allometric scaling of left cardiac dimensions to account for differences in bodyweight, there was a significant effect of age on LA and LV dimensions. (TTE parameters: LAN, LVEDVI andLVESVI p<0.001; LVIDDN p=0.015; LVIDSN p=0.003. CMR parameters: LVMi, LVEDVI, LVESVI, LV mass:femur and LV mass:L5, p<0.001 Figure 1) but there was a positive interaction with genotype (see below, figure 1 and table 2).The effect of age on echocardiographic linear dimensions persisted when LV diastolic diameter was dimensions were indexed to aortic diameter, (LA:Ao p=0.043; LVIDd:Ao p=0.013) except for LVIDs:Ao (LVIDs:Ao p=0.121).

**Figure 1:**
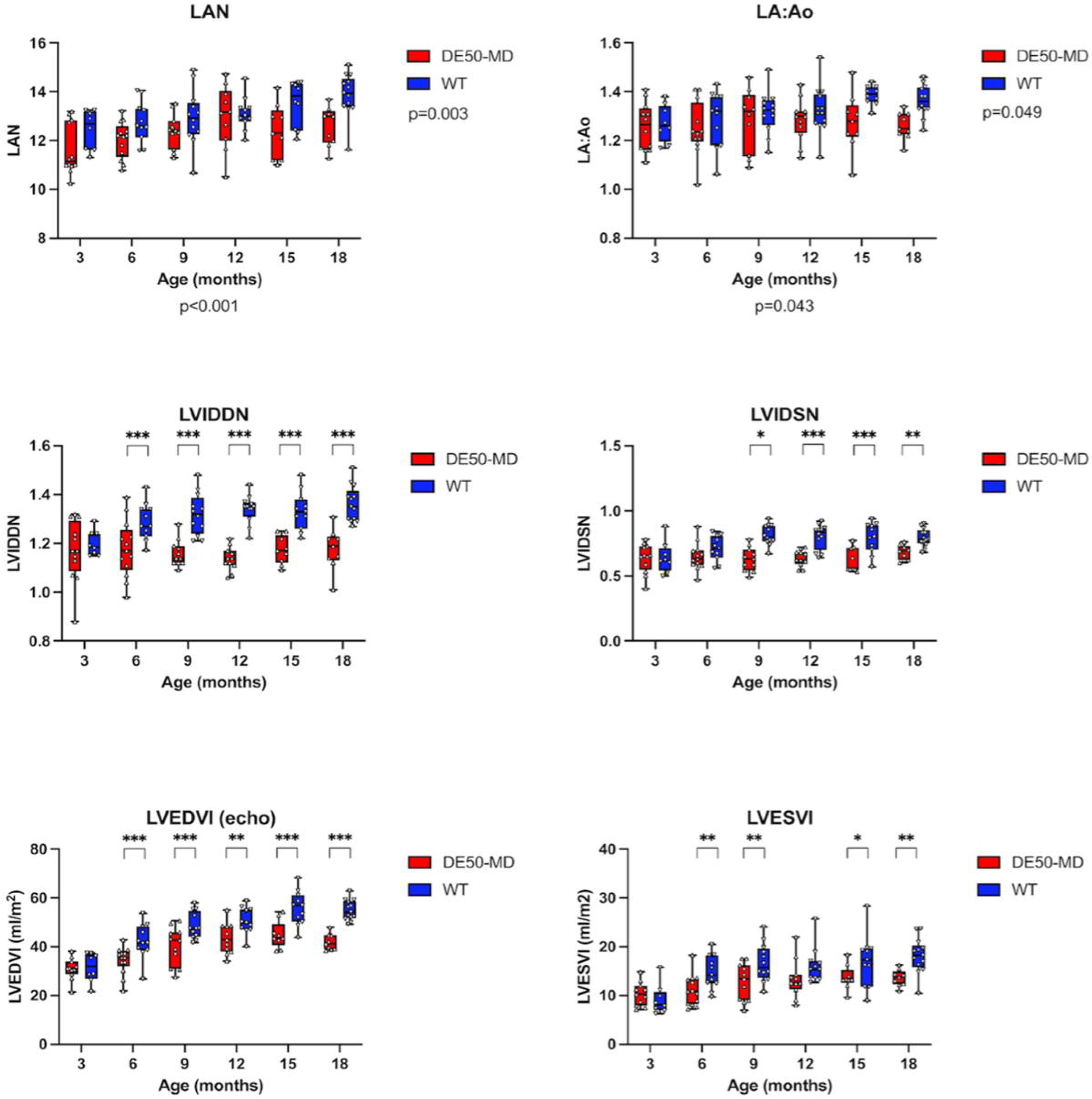
Chamber dimensions measured from conscious echocardiograms of age-matched wild type (WT, n=6-8) and DESO-MD dogs (n=7-11), performed at 3-month intervals between 3 and 18 months of age. fudividual dogs are represented by staggered dots. Whiskers represent the minimum to maximum range. Horizontal lines represent median values. P values represent results of linear mixed model (LMM) analysis of repeated measures to explore effects of genotype (to the right of each graph) and age (below x axes). Where an interaction between genotype and age was established p values are not shown, instead significant differences, where present, for each age point studied are represented by asterisks as follows: *p<0.05, **p<0.01, ***p<0.001. *Abbreviations: WT, Wild Type. LA:Ao, Left atrial indexed to aortic short-axis diameter. LAN, Left atrial diameter indexed to bodyweight^0.324. LVIDDN, Diastolic left ventricular (LV) internal diameter indexed to bodyweight^0.322. LVIDSN, Systolic LV internal diameter indexed to bodyweight^0.346. LVEDVI, LV end diastolic volume indexed to body surface are. LVESVI, LV end systolic volume indexed to body surface area. The graphs demonstrate the smaller left atrial and left ventricular diastolic and systolic dimensions in DE50-MD dogs compared to age-matched WT dogs. The difference was most pronounced for the LV. Systolic LV volume and diastolic LV dimensions were significantly smaller from 6 months of age, whereas weight normalised systolic LV internal diameter was smaller from 9 months of age*.

**Figure 2:**
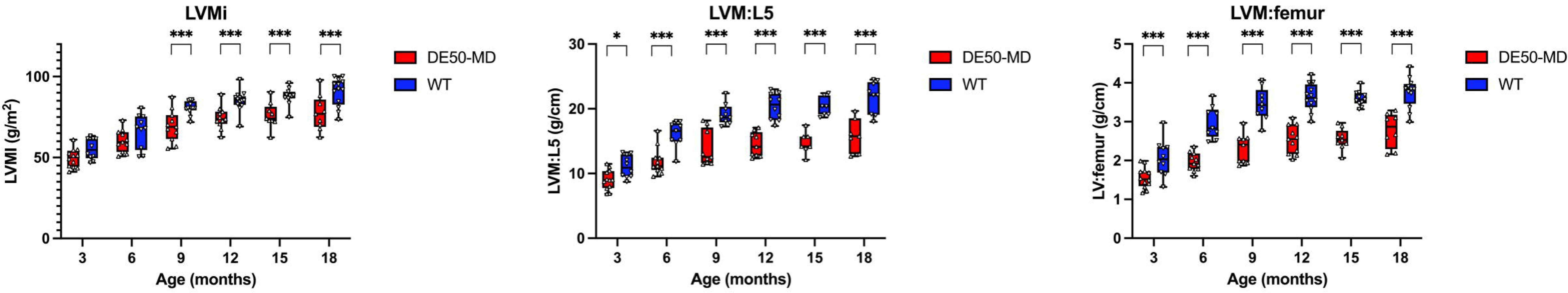
Different methods of normalisation for left ventricular mass (LVM) measured using cardiac magnetic resonance imaging in age-matched wild type (WT, n=6-8) and DESO-MD (n=7-ll) dogs, performed at 3-month intervals between 3 and 18 months of age. Individual dogs are represented by staggered dots. Whiskers represent the minimum to maximum range. Horizontal lines represent median values. For each age point studied significant differences, where present, are represented by asterisks as follows: *p<0.05, **p<0.01, ***p<0.001. *Abbreviations: WT, Wild Type. LVM Left ventricular mass. LVMi: LVM indexed to body surface area. LVM:L5, LVM indexed to the length of the fifth lumbar vertebral body (L5), measured from magnetic resonance images. LVM:femur, LVM indexed to the length of the femur measured from magnetic resonance images. The graphs demonstrate the smaller LVM of DE50-MD dogs compared to age-matched WT dogs irrespective of the method of normalisation. The difference became significant from 9 months of age for LVMi, but was even earlier (3 months of age) when LVM was indexed to femur and L5 length*

**Table 2:**
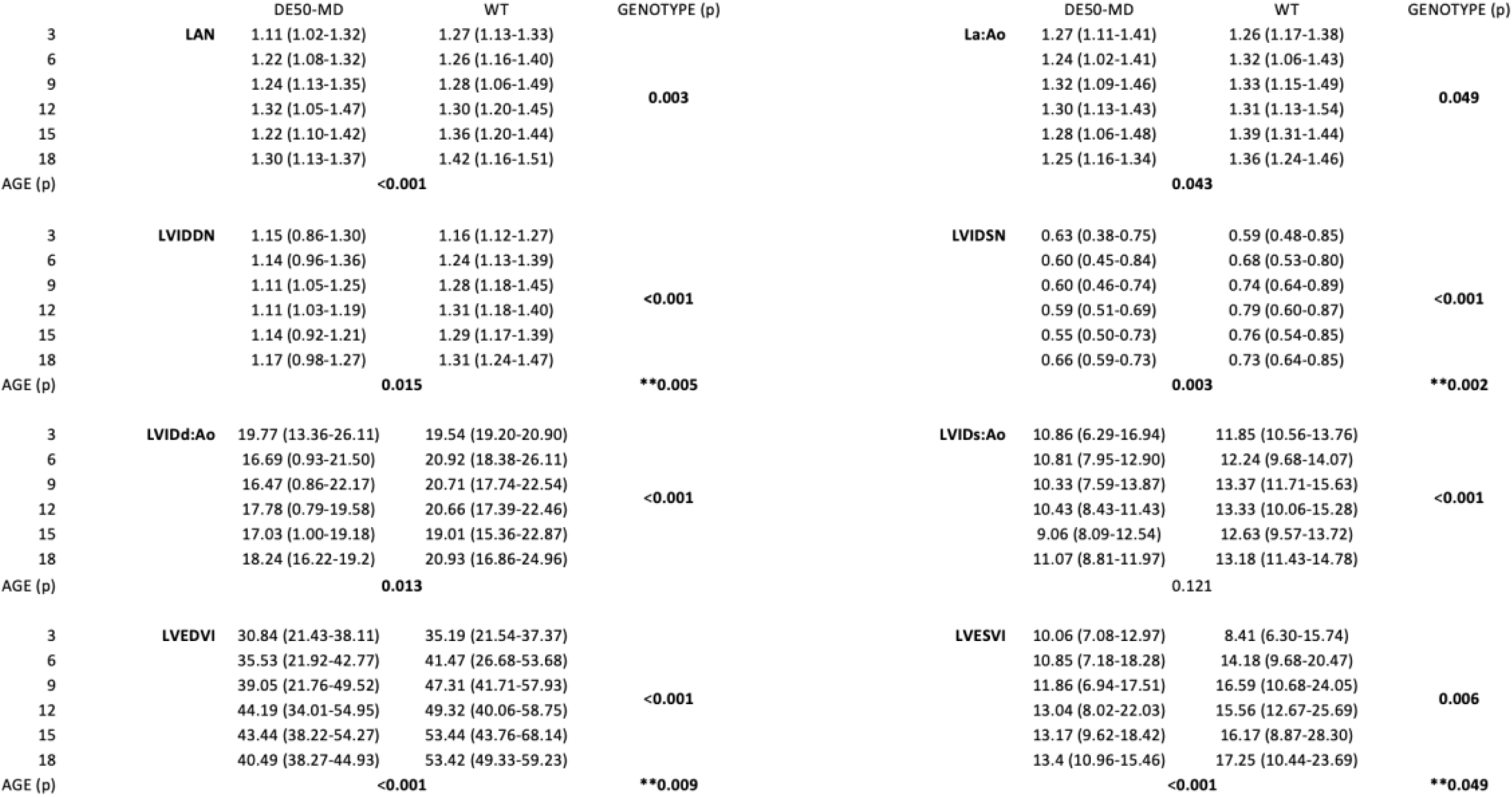
Results of linear mixed model analysis to explore the effect of genotype and age on structural and functional assessments using conscious transthoracic echocardiography. Abbreviations: WT; Wild type control dogs. LA:Ao, Left atrum indexed to aortic short-axis diameter. LAN, Left atrial diameter indexed to bodyweight^0.324. LVIDDN, Diastolic left ventricular (LV) internal diameter indexed to bodyweight^0.322. LVIDSN, Systolic LV internal diameter indexed to bodyweight^0.346. LVIDd:Ao, Diastolic LV internal diameter indexed to long axis aortic diameter. LVIDs:Ao, Systolic LV internal diameter indexed to long axis aortic diameter. LVEDVI, LV end diastolic volume indexed to body surface are. LVESVI, LV end systolic volume indexed to body surface area. ** denotes significant interaction between genotype and age.

The DE50-MD dogs had significantly smaller weight-adjusted, (systolic and diastolic) left cardiac chamber dimensions across all ages (LAN p=0.003, LVIDdN and LVIDsN p<0.001). Significance was retained irrespective of whether normalisation was made according to body weight or aortic diameter (La:Ao p=0.049, LVIDd:Ao and LVIDs:Ao p<0.001). The difference in LV diastolic internal dimensions measured using TTE (LVEDVI and LVIDDN) became significant from 6 months of age (p<0.001), whilst CMR-derived LVEDVI was significantly reduced from 3 months of age, (p=0.006). DE50-MD dogs also had lower LVMi, the difference becoming significant from 9 months of age (p=0.006). In dogs where data were available for femoral and L5 vertebral length, the adjusted LV mass was significantly lower in DE50-MD dogs from 3 months of age (p<0.001) for both indices.

### Left ventricular geometry

The LV SI value was higher in DE50-MD dogs from 6 months of age, (p=0.002) reflecting a less spherical LV than WT dogs. The LVRI index significantly increased with age (p<0.001) and, despite the lower LVMi in DE50-MD dogs, LVRI was statistically increased in DE50-MD dogs compared with WT dogs, (p=0.001) representing a disproportionally low LV diastolic volume relative to LV mass. This was paralleled by an increase in RWT in DE50-MD dogs, (LnRWT, p=0.017) which, like LVRI, increased with age, (p=0.014). Overall, the LA diameter of DE50-MD dogs was wider relative to diastolic LV internal diameter than those of WT dogs (p=0.041), although when individual time points were examined, statistical significance was only reached at 9 (p=0.015) and 12 (p<0.001) months. (Results are summarised in figure 3.)

**Figure 3:**
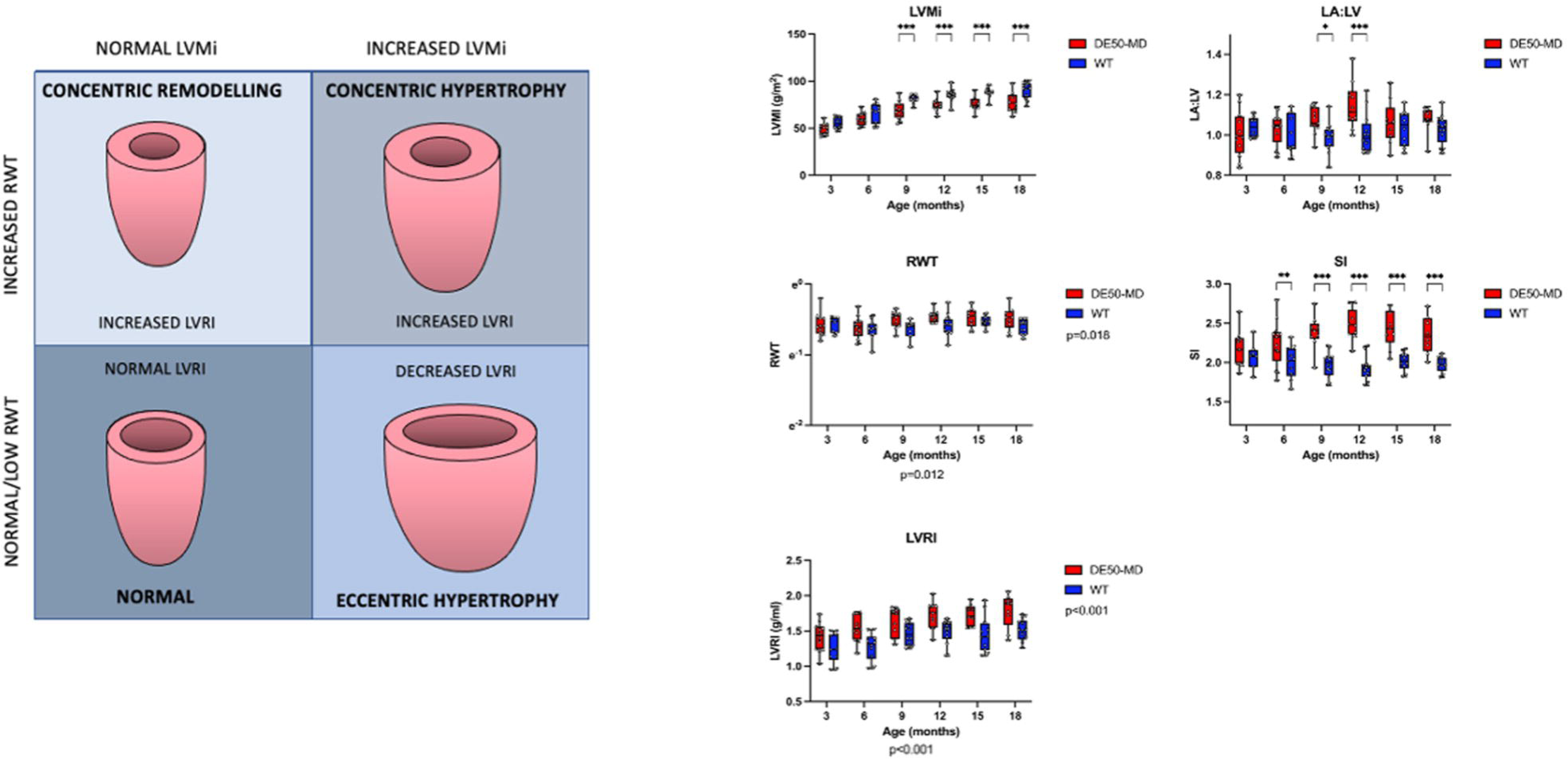
**Left panel: Different forms of left ventricular (LV) geometric remodeling as defined in people.** Patients with increased LV Mass indexed to body surface area (LVMi) are considered to have hypertrophy. Hypertrophy in the presence of normal or reduced relative wall thickness (RWT) is eccentric hypertrophy. Where hypertrophy is present with increased RWT it is considered concentric hypertrophy. IfRWT is increased but LVMi is within the healthy reference range, the geometry is consistent with concentric remodelling, a subtle adaptive change in LV geometry secondary to increased afterload (aortic stenosis or systemic hypertension). Left ventricular remodelling index (LVRI) is the relationship between LV mass and LV end diastolic volume. Increased LVRI signifies concentric hypertrophy or remodelling while reduced LVRI supports eccentric hypertrophy. **Right panel: Geometric assessments in age-matched wild type (WT) and DE50-MD dogs,** performed at 3-month intervals between 3 and 18 months of age. Individual dogs are represented by staggered dots. Whiskers represent the minimum to maximum range. Horizontal lines represent median values. For each age point studied significant differences, where present, are represented by asterisks as follows: *p<0.05, **p<0.01, ***p<0.001. DE50-MD dogs have increased LVRI and RWT, but lower LVMi, and so the geometry is not supportive of hypertrophy. The LA is mildly increased compared to the LV (although significance was only achieved at 9 and 12 months of age). These results, in combination with the reduced LV sphericity (denoted by higher values for SI) in DE50-MD dogs supports a picture of reduced LV mass with a disproportionate reduction in LV volume. Abbreviations: WT, Wild Type. LVMi Left ventricular mass indexed to body surface area. LVRI, Left ventricular remodeling index. RWT, Relative wall thickness. SI, Sphericity index. LA, left atrial diameter. LV, Left ventricular diameter

### Left ventricular function

Global LV systolic function parameters were similar in DE50-MD and WT dogs. Ejection fraction exceeded 55% and FS was above 27% in all dogs, but FS declined with age in both groups (p=0.007). There was a trend for higher FS in DE50-MD dogs, (p=0.07) which became statistically significant at 9 and 15 months of age (p=0.004 and p=0.019 respectively). Ejection fraction was similar between DE50-MD and WT dogs (p=0.660) and there was no effect of age (p=0.376). There was a trend for lower longitudinal systolic velocities of the basal IVS (but not the basal LV anterolateral wall (p=0.450)) in DE50-MD dogs, but this did not achieve statistical significance (p=0.056). Stroke volume index increased with age in all dogs (p<0.001) but SVI was statistically lower in DE50-MD dogs than WT dogs, (p<0.001) with no interaction between genotype and age, (p=0.118). Despite this finding, the cardiac index (CI) was similar between groups (p=0.092) and did not change with age (p=0.734). Results are summarised in figure 4.

**Figure 4:**
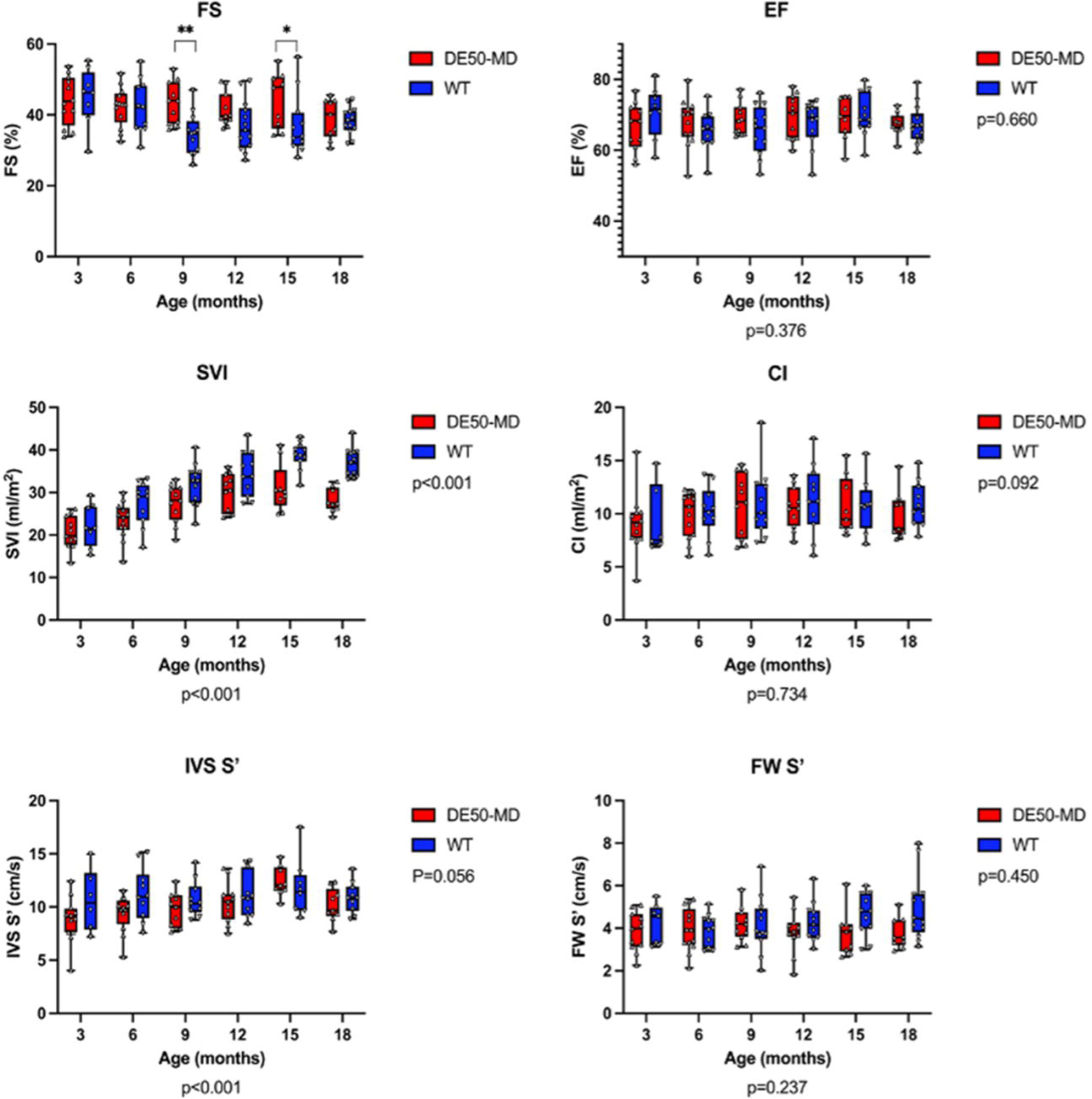
Left ventricular (LV) systolic function parameters derived from conscious echocardiograms of age-matched wild type (WT, n=6-9) and DESO-MD dogs (n=8-11), performed at 3-month intervals between 3 and 18 months of age. Individual dogs are represented by staggered dots. Whiskers represent the minimum to maximum range. Horizontal lines represent median values. P values represent results of linear mixed model (LMM) analysis of repeated measures to explore effects of genotype (to the right of each graph) and age (below x axes). Where an interaction between genotype and age was established, p values are not shown, instead significant differences, where present, for each age point studied are represented as follows: *p<0.05, **p<0.01. *Abbreviations: WT, Wild Type. FS, Fractional shortening. EF, Ejection fraction. SVI, stroke volume index. CL Cardiac index,* S’ *systolic longitudinal myocardial velocity measured using pulse waved tissue Doppler imaging at the basal septa/ (IVS) and inferolateral (FW) L V wall.* The graphs demonstrate the similar systolic function in DESO-MD dogs and age-matched WT dogs. The SVI in DESO-MD dogs was significantly smaller across ages but CI was maintained due to the higher heart rates in DESO-MD dogs. Results of LMM analysis demonstrated hypercontractile FS in DESO-MD dogs at 9 and 15 months.

Spectral Doppler interrogation of mitral inflow was attempted in all dogs but there was persistent summation of early and late diastolic flow profiles in 1 (9-month-old) WT dog and in 4 DE50-MD dogs, (aged 3 (n=2), 6 (n=1) and 9 months, (n=1)) associated with persistently high heart rates. The ratio of early to late diastolic filling was >1 in all dogs at all ages. The velocity of early diastolic flow tended to be lower in the DE50-MD dogs, but this did not reach statistical significance (p=0.064). There was no significant group effect on velocity of late diastolic filling (p=0.173), the ratio of early to late diastolic filling (p=0.628), IVRT (p=0.624) or E:IVRT (p=0.164). Both IVRT and E:IVRT increased with age in all dogs (p<0.001 for both).

Longitudinal pulsed-wave TDI Interrogation of the LVFW and basal IVS revealed reduced early diastolic myocardial velocities of the LVFW only (LVFW E’ p <0.001. LnIVS E’ p=0.188) in DE50-MD dogs compared to WT dogs, with no effect of age (p=0.527 and p=0.119 respectively). There was an overall reduction in LVFW E’A’ in DE50-MD dogs (p=0.029) that was significant from 15 and 18 months of age (p=0.021 and p<0.001 respectively) but the septal E’A’ was similar between age-matched DE50-MD and WT dogs (p=0.674, age effect p=0.504). Results are summarised in table 3.

**Table 3:**
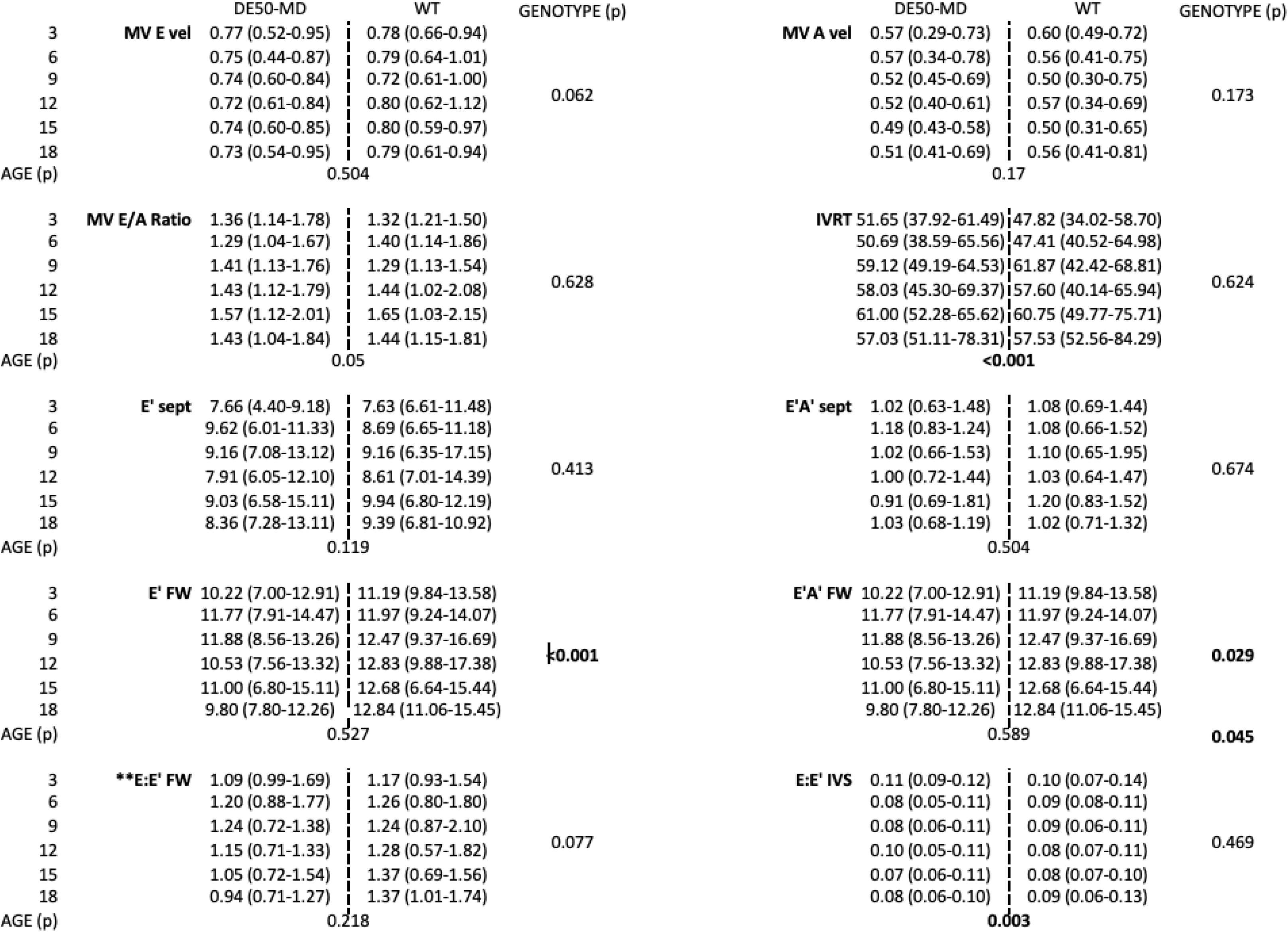
Results of linear mixed model analysis to explore the effect of genotype and age on diastolic functional assessments using conscious transthoracic echocardiography. Abbreviations: WT; Wild type control dogs. LA:Ao, Left atrial indexed to aortic short-axis diameter. LAN, Left atrial diameter indexed to bodyweight^0.324^. LVIDDN, Diastolic left ventricular (LV) internal diameter indexed to bodyweight^0.322^. LVIDSN, Systolic LV internal diameter indexed to bodyweight^0.346^. LVIDd:Ao Diastolic LV internal diameter indexed to long axis aortic diameter. LVIDs:Ao Systolic LV internal diameter indexed to long axis aortic diameter. LVEDVI, LV end diastolic volume indexed to body surface are. LVESVI, LV end systolic volume indexed to body surface area. ** denotes significant interaction between genotype and age.

#### Radial Myocardial Velocity Gradients (Table 4)

Analysis of MVGs revealed abnormalities in radial deformation parameters from the inferolateral wall in DE50-MD dogs. As expected, myocardial velocities were higher in the endocardial layer than the epicardial layer in both groups of dogs but the systolic (p=0.018) and diastolic (early p<0.001, active p=0.027) MVGs were reduced in DE50-MD dogs. The ratio between early and active diastolic MVGs (p=0.012) was also reduced in this group. The reduced MVG was due to lower peak endomyocardial systolic (p=0.05) and early diastolic (p=0.002) myocardial velocities in DE50-MD, while epicardial velocities (systolic and diastolic) were preserved. There was no significant effect of age on radial myocardial velocities or MVGs, except for early epicardial velocity, which increased with age (p=0.03), (see table 4).

**Table 4:** Results of linear mixed model analysis to explore the effect of genotype and age on radial inferolateral wall systolic and diastolic regional velocities and myocardial velocity gradient. Abbreviations (S; systolic. E; early diastolic. A; active diastolic. MVG; myocardial velocity gradient).

|  | AGE (MONTHS) | DE50-MD | WT | Genotype (p) |  | DE50-MD | WT | Genotype (p) |  | DE50-MD | WT | Genotype (p) |
| --- | --- | --- | --- | --- | --- | --- | --- | --- | --- | --- | --- | --- |
| EPICARDIAL S | 3 | 3.64 (2.14,5.38) | 4.37 (3.37,5.38) | p=0.216 | ENDOCARDIAL S | 5.45 (3.74,6.62) | 6.32 (5.54,6.74) | p=0.05 | MVG S | 1.59 (1.23,2.47) | 1.78 (1.24,2.17) | p=0.018 |
| (cm/s) | 6 | 4.50 (2.73,5.27) | 4.24 (2.98,8.15) |  | (cm/s) | 6.31 (3.94,6.79) | 5.73 (4.79,10.98) |  | (cm/s) | 1.54 (1.21,2.36) | 1.86 (1.26,2.83) |  |
|  | 9 | 4.23 (3.09,5.35) | 4.36 (3.28,6.01) |  |  | 6.14 (4.10,7.01) | 6.63 (5.06,8.01) |  |  | 1.84 (0.61,2.34) | 1.84 (1.03,2.98) |  |
|  | 12 | 4.24 (3.26,4.88) | 4.15 (2.72,6.85) |  |  | 5.78 (4.24,7.29) | 6.05 (4.78,8.53) |  |  | 1.90 (0.71,2.46) | 1.74 (1.47,2.79) |  |
|  | 15 | 4.58 (2.40,5.75) | 4.71 (2.81,6.59) |  |  | 6.20 (3.84,7.30) | 6.58 (4.87,9.08) |  |  | 1.50 (0.86,2.35) | 1.78 (1.12,2.49) |  |
|  | 18 | 4.04 (3.04,6.41) | 4.37 (3.53,6.29) |  |  | 5.99 (4.13,8.05) | 6.43 (4.89,9.71) |  |  | 1.64 (0.08,2.29) | 2.14 (1.34,3.57) |  |
| Age (p) |  | p=0.794 |  |  |  | p=0.802 |  |  |  | p=0.853 |  |  |
| EPICARDIAL E | 3 | -1.83 (-3.28,-1.20) | -2.38 (-3.36,-0.49) | p=0.555 | ENDOCARDIAL E | -4.01 (-5.65,-2.70) | -4.30 (-6.53,-2.09) | p=0.002 | MVG E | 2.38 (0.87,3.51) | 2.29 (0.87,3.17) | p<.001 |
| (cm/s) | 6 | -1.50 (-5.04,-0.02) | -3.41 (-4.09,-2.09) |  | (cm/s) | -5.07 (-7.35,-1.51) | -5.32 (-6.92,-1.70) |  | (cm/s) | 2.27 (1.41,3.67) | 2.52 (1.62,3.51) |  |
|  | 9 | -3.06 (-5.91,-0.94) | -2.76 (-5.06,-1.10) |  |  | -4.60 (-8.04,-3.44) | -5.24 (-7.36,-4.24) |  |  | 1.88 (0.40,2.63) | 2.51 (1.76,3.85) |  |
|  | 12 | -3.94 (-4.70,-1.77) | -3.65 (-5.15,-2.03) |  |  | -5.04 (-7.29,-3.57) | -6.40 (-7.67,-4.24) |  |  | 1.60 (0.36,2.73) | 2.75 (1.70,3.90) |  |
|  | 15 | -3.52 (-5.45,-0.73) | -3.23 (-5.28,-1.39) |  |  | -5.11 (-6.88,-3.34) | -6.66 (-8.37,-4.28) |  |  | 1.33 (0.78,2.74) | 3.01 (1.33,4.39) |  |
|  | 18 | -2.09 (-4.47,-0.84) | -3.78 (-6.74,-1.70) |  |  | -4.48 (-6.65,-2.55) | -6.26 (-12.53,-4.87) |  |  | 2.16 (0.08,3.63) | 3.18 (1.06,5.50) |  |
| Age (p) |  | p=0.026 |  |  |  | p=0.22 |  |  |  | p=0.621 |  |  |
| EPICARDIAL A | 3 | -2.40 (-4.02,-1.56) | -3.05 (-4.20,-1.92) | p=0.764 | ENDOCARDIAL A | -3.99 (-6.13,-3.29) | -4.85 (-5.57,-3.29) | p=0.23 | MVG A | 1.75 (1.34,2.11) | 1.37 (1.25,2.34) | p=0.039 |
| (cm/s) | 6 | -2.81 (-3.23,-1.04) | -2.88 (-3.77,-0.86) |  | (cm/s) | -4.71 (-6.85,-2.67) | -4.56 (-6.19,-2.74) |  | (cm/s) | 1.74 (1.41,4.05) | 1.88 (1.69,2.42) |  |
|  | 9 | -2.81 (-4.35,-1.54) | -2.60 (-4.04,-1.50) |  |  | -4.60 (-6.09,-3.32) | -4.25 (-6.57,-3.57) |  |  | 1.79 (1.09,2.51) | 2.05 (1.18,3.51) |  |
|  | 12 | -2.68 (-3.92,-1.45) | -2.58 (-4.26,-0.89) |  |  | -4.60 (-6.19,-2.55) | -4.56 (-6.43,-3.37) |  |  | 1.40 (0.59,2.33) | 2.01 (1.12,3.29) |  |
|  | 15 | -3.05 (-5.37,-1.26) | -2.81 (-4.49,-1.94) |  |  | -4.65 (-6.88,-2.83) | -5.13 (-6.78,-3.45) |  |  | 1.60 (0.60,2.38) | 2.08 (0.52,3.52) |  |
|  | 18 | -2.69 (-4.59,-1.32) | -3.12 (-5.00,-1.37) |  |  | -4.74 (-5.85,-2.95) | -4.80 (-7.59,-3.98) |  |  | 1.79 (0.26,2.44) | 2.16 (0.44,3.31) |  |
| Age (p) |  | p=0.451 |  |  |  | p=0.673 |  |  |  | p=0.586 |  |  |

#### Speckle Tracking Echocardiography

Speckle tracking echocardiographic assessment was made challenging by limited acoustic windows in DE50-MD dogs, and panting/patient agitation in some WT dogs. Consequently only targeted right parasternal views were collected from those dogs. Poor image quality and/or inadequate tracking by the software led to exclusion of 7/49 available studies from WT dogs and 4/40 studies from DE50-MD dogs for circumferential strain analysis. For longitudinal strain analysis, 10/47 available studies were excluded from WT dogs and 3/40 DE50MD dogs. There was no significant difference in the heart rate, or the achieved frame rate recorded for WT and DE50-MD dogs.

#### Average longitudinal and circumferential peak strain and strain rate

Average peak longitudinal systolic strain (peak LS) values (figure 5, supplementary table 4) were similar between WT and DE50-MD dogs, (p=0.162) and this finding did not change significantly with age, (p=0.663). There was no significant effect of genotype or age on values for average longitudinal SSR, (p=0.793 and p=0.667 respectively) ESR, (p=0.280 and p=0.0.363 respectively) or ASR, (p=0.899 and p=0.326 respectively).

**Figure 5:**
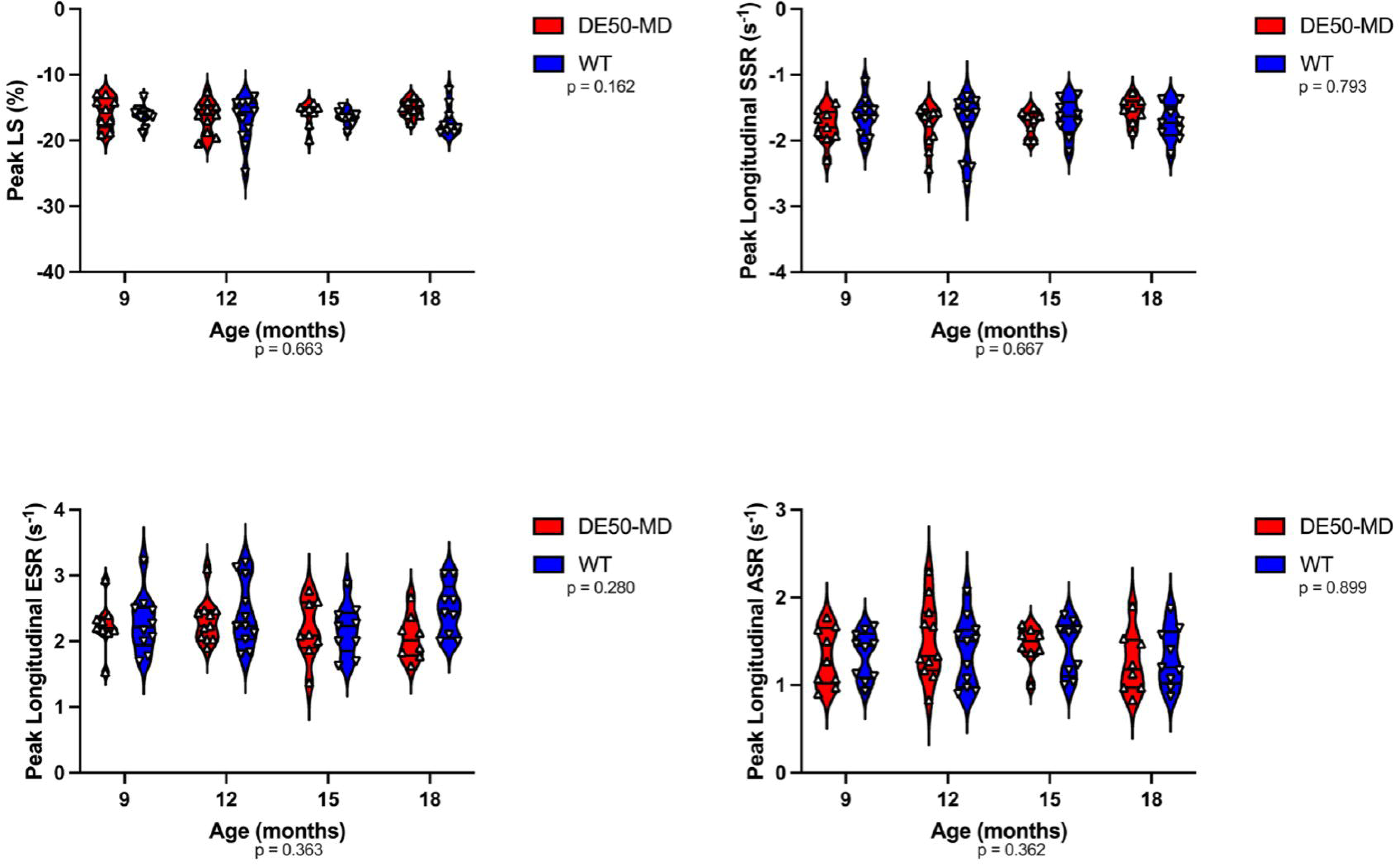
Average longitudinal strain and strain rate in WT (n=9-11) and DESO-MD (n=S-11) dogs with age. A) Average peak longitudinal strain (Peak LS, %). B) Systolic longitudinal strain rate (longitudinal SSR, s-1), C) Early diastolic longitudinal strain rate (longitudinal ESR, s-1) and D) Active diastolic longitudinal strain rate (longitudinal ASR, s-1). Each staggered point (triangle) represents an individual dog. Horizontal lines are present at the median, 25%, and 75% percentiles. Letters denote differences in the mean(p<0.05). Note that Peak LS, SSR, ESR and ASR are similar between genotypes with no significant effect of age.

Average peak circumferential systolic strain, (peak CS) values were more negative in DE50-MD dogs, implying hypercontractility in this plane at the mid-ventricular level (p=0.003). Peak CS declined, (became less negative) between 15 and 18 months of age in the DE50-MD dogs, (figure 6, supplementary table 4) such that values approximated those of WT dogs (-22.92±3.96 % vs -22.0±3.24% respectively; p=0.112, Age*Group interaction p=0.046) at 18 months of age. On further inspection, the decline in the median peak circumferential strain in the DE50-MD dogs was due to a small number of dogs with values >-20%. There was no significant difference in average circumferential SSR between DE50-MD versus WT dogs (p= 0.098). The circumferential ESR and ASR were increased in the DE50-MD dogs compared to the WT dogs (p=0.006 and p=0.016 respectively); active circumferential strain rate values required natural log transformation to achieve normality of the LMM residuals), with no effect of age on any strain rate parameter (SSR p=0.418, ESR p=0.688, ASR p=0.770).

**Figure 6:**
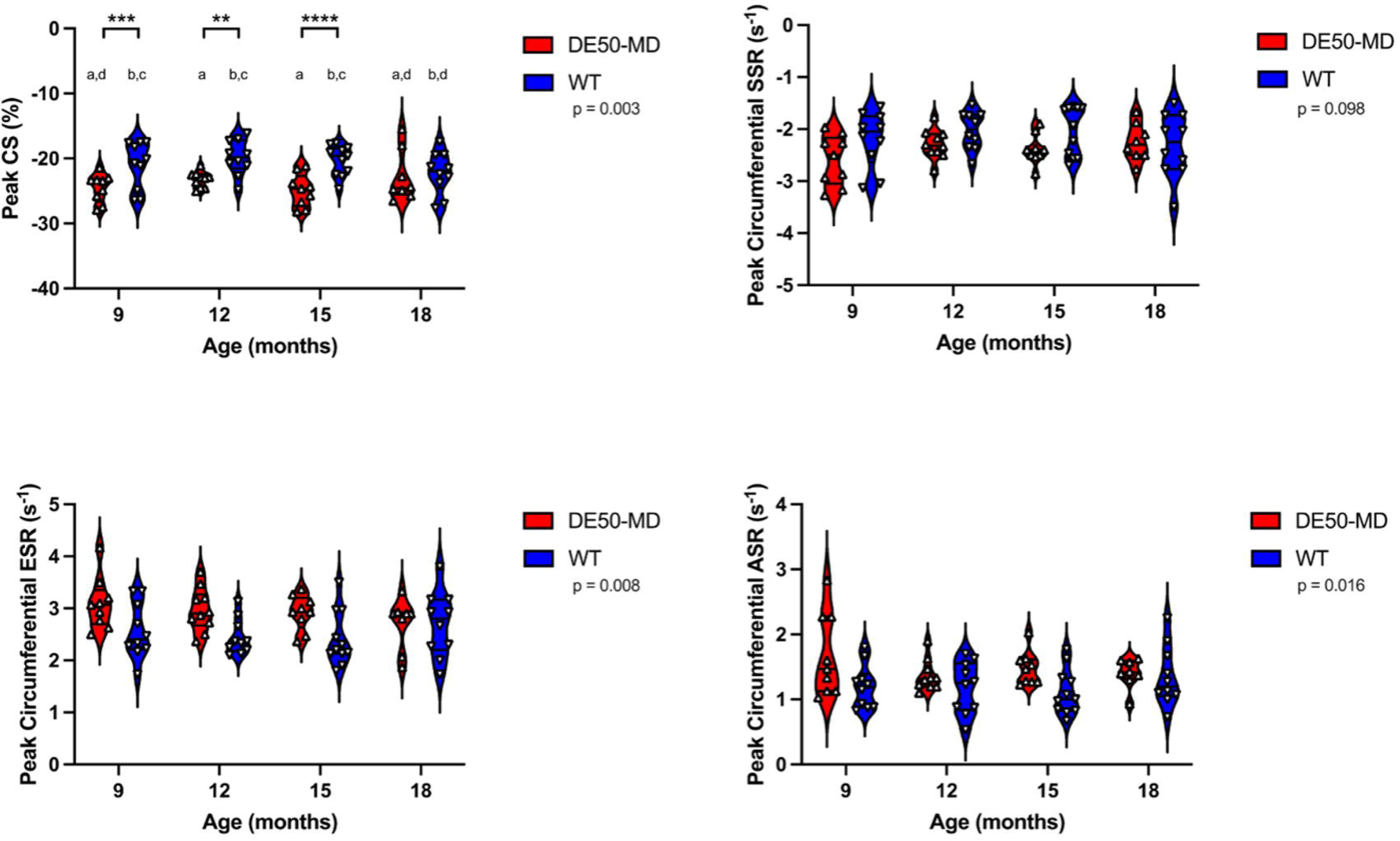
Average midventricular circumferential strain and strain rate in WT (n=l0-11) and DESO-MD (n=S-10) dogs with age. A) Average peak circumferential strain (Peak CS, %). Letters denote differences in the mean. Means sharing a letter are not significantly different (p > 0.05). ** p=0.01, *** p<0.005, ****p<0.001. B) Systolic circumferential strain rate (circumferential SSR, s-1), C) Early diastolic circumferential strain rate (circumferential ESR, s-1) and D) Active diastolic circumferential strain rate (Ln transformed circumferential ASR, s-1). Each staggered point (triangle) represents an individual dog. Horizontal lines are present at the median, 25%, and 75% percentiles Note that the magnitude of Peak CS increased (i.e. more negative) in DE50-MD dogs between 9 and 15 months of age. Likewise, ESR and (ln)ASR are increased in DE50-MD dogs across all ages. Peak SSR is similar between genotypes.

#### Peak Longitudinal wall region and segmental strain

Peak longitudinal strain values were heterogenous amongst wall segments in both the WT and DE50-MD dogs.

#### A: Septum vs anterolateral wall peak strain and strain rate

Septal wall strain values were more negative than those of the anterolateral wall in both groups of dogs, (p<0.001 for both genotypes, Supplementary table 5). There was a significant interaction between wall region and genotype for both longitudinal ESR and ASR: Longitudinal ESR and ASR were lower in the anterolateral wall of DE50MD dogs compared to WT dogs (p=0.001 and p=0.02 respectively, table 5) but there was no significant effect of genotype on septal diastolic longitudinal deformation rates.

**Table 5:** Results of linear mixed model analysis, describing the overall effect of genotype and age on regional diastolic strain rate within each genotype. Tabulated values are expressed as median (minimum, maximum). Abbreviations: ESR; early diastolic longitudinal strain rate. ASR; active longitudinal strain rate. diastolic strain rate. Longitudinal ESR and ASR were lower in the anterolateral wall of DE50MD dogs compared to WT dogs but there was no significant effect of genotype on septal diastolic longitudinal deformation rates. There was no interaction between genotype and age in LMM analysis for ESR or ASR.

| SEPTUM |  |  |  | ANTEROLATERAL |  |  |  |
| --- | --- | --- | --- | --- | --- | --- | --- |
| ESR | DE50-MD | WT | Group | DE50-MD | WT | Group |  |
| (s<sup>-1</sup>) |  |  |  |  |  |  |  |
| 9 | 3.02 (2.21,3.48) | 2.65 (1.91,3.93) | $p=0.295$ | 2.69 (2.22,3.68) | 3.66 (2.58,4.26) | $p<0.001$ | |
| 12 | 2.98 (2.18,3.75) | 2.99 (1.97,4.45) |  | 2.72 (2.14,3.81) | 3.16 (2.23,4.34) |  |  |
| 15 | 2.43 (2.22,2.86) | 2.90 (1.97,3.62) |  | 2.75 (1.77,3.11) | 3.35 (1.99,4.33) |  |  |
| 18 | 2.64 (2.24,3.38) | 3.34 (2.17,4.19) |  | 2.39 (1.48,3.39) | 3.52 (2.96,4.05) |  |  |
| | $p=0.420$ | | | | | | |
| SEPTUM |  |  |  | ANTEROLATERAL |  |  |  |
| ASR | DE50-MD | WT | Group | DE50-MD | WT | Group |  |
| (s<sup>-1</sup>) |  |  |  |  |  |  |  |
| 9 | 2.01 (1.34,2.57) | 1.68 (1.06,2.57) | $p=0.131$ | 1.39 (0.14,1.49) | 1.55 (0.48,2.02) | $p=0.02$ | |
| 12 | 1.85 (1.25,2.59) | 1.60 (1.11,2.06) |  | 1.44 (0.49,2.21) | 1.64 (0.85,2.11) |  |  |
| 15 | 1.70 (1.53,2.38) | 1.78 (1.29,1.93) |  | 1.40 (0.86,1.92) | 1.72 (0.98,2.51) |  |  |
| 18 | 1.49 (1.00,2.19) | 1.65 (1.13,2.04) |  | 1.33 (0.62,2.15) | 1.66 (0.88,2.29) |  |  |
| | $p=0.559$ | | | | | | |

#### B: Apex, Mid and Basal segment systolic strain

A significant interaction between genotype and wall segment on peak longitudinal strain was identified in LMM analysis. WT dogs demonstrated a base to apex longitudinal peak strain gradient in the septal and anterolateral wall, with highest peak longitudinal systolic strain values being documented in apical segments (figure 7). In contrast, DE50-MD dogs demonstrated a lack of the base to apex gradient. This was due to depressed (less negative) longitudinal strain in the septal and anterolateral apical segments, such that Fisher’s post hoc comparisons demonstrated significantly attenuated longitudinal systolic strain in these segments in the DE50-MD dogs compared to corresponding apical segments of WT dogs (p=0.004 and p<0.001 for the septum and anterolateral wall respectively).

**Figure 7:**
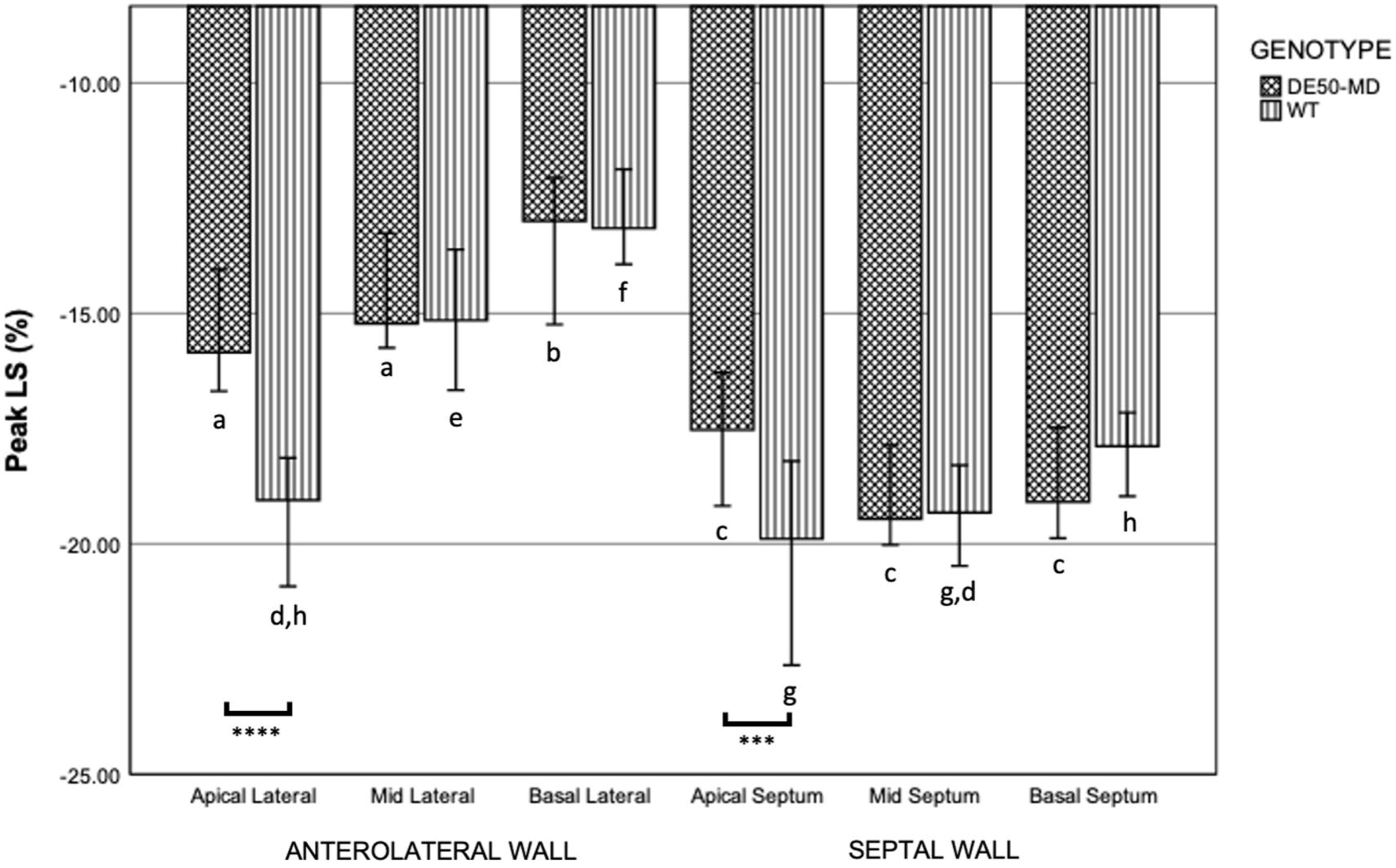
Median segmental peak longitudinal systolic strain values in WT (n=9-11) and DESO-MD (n=9-ll) dogs across all age groups. Error bars represent the 95% confidence intervals. Letters denote differences in longitudinal strain according to segment within each genotype. Segments sharing a letter are not significantly different (p > 0.05).*** p<0.005, ****p<0.001.

#### Peak Circumferential segmental strain

Peak circumferential strain values were heterogenous amongst wall segments in WT and DE50-MD dogs (p<0.001). Segmental systolic strain values increased in a counterclockwise direction from the inferolateral wall segments, such that septal segments had the most negative (greatest) peak circumferential strain values, whilst the inferolateral segments had the most positive (weakest) peak, circumferential strain values (figure 8). Strain values for individual midventricular wall segments were similar between DE50-MD and WT dogs except for the anterolateral and inferoseptal wall segments, being comparatively hypercontractile in DE50-MD dogs (-24.29±4.93% vs -20.26±4.64% and -28.45±3.62% vs -24.11±3.65%, p=0.003 and p=0.002 respectively).

**Figure 8:**
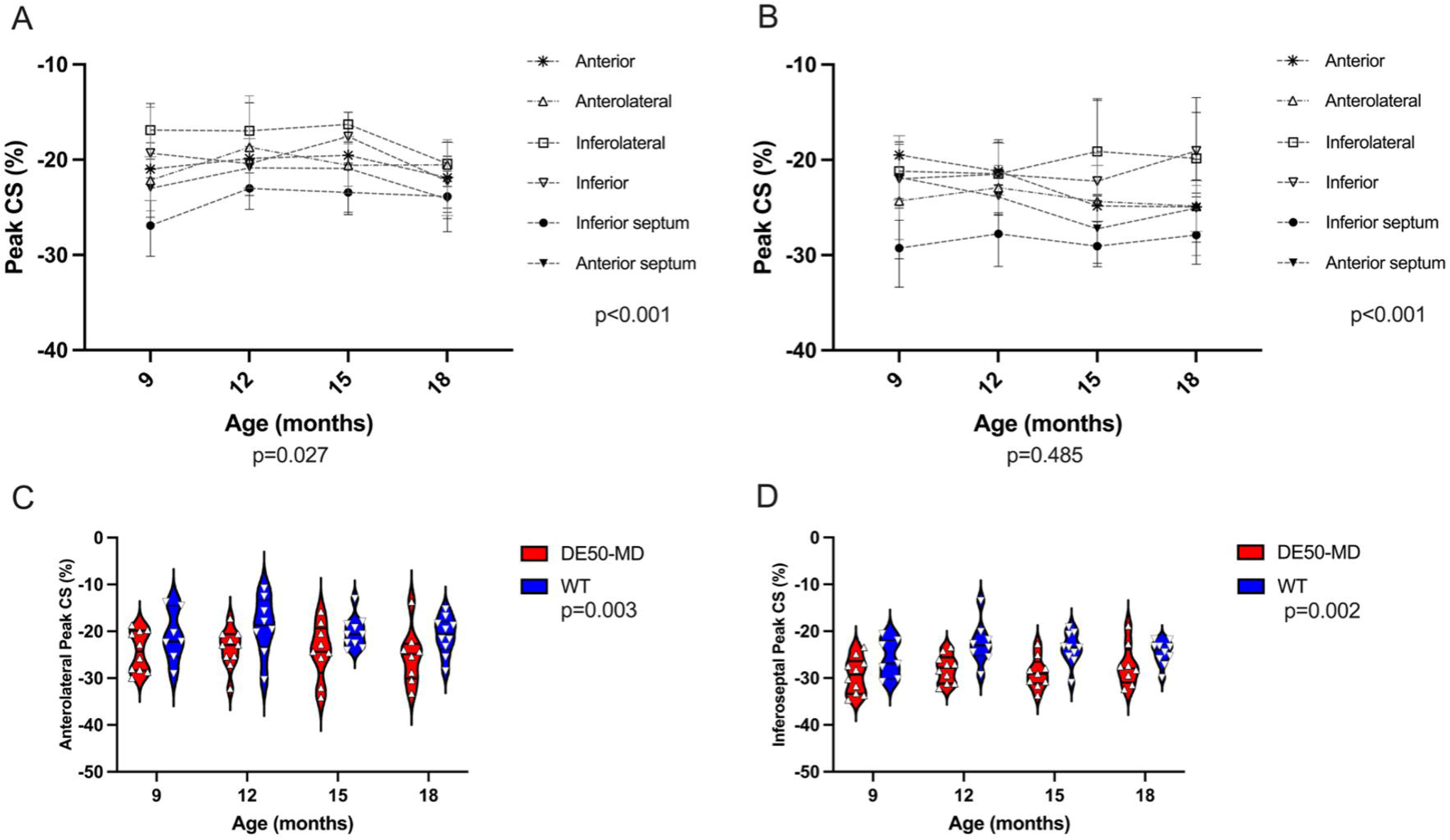
**Upper panel: Change in midventricular segment peak circumferential strain with age in A) WT (n=7-10) and B) DESO-MD (n=S-10) dogs.** Each symbol represents the median value for a specific midventricular wall segment. Error values depict interquartile range. **Lower panel: Peak circumferential strain of C) the anterolateral and D) the inferoseptal midventricular wall segments in WT and DESO-MD dogs.** P values represent the results of (A & B) within genotype and (C &D) between genotype analysis with LMM, evaluating the influence of wall segment and age, and genotype and age respectively. Note the statistically significant heterogeneity between segmental circumferential strain values in both DE50-MD dogs and WT dogs and relative hypercontractility of the anterolateral and inferoseptal wall segments in DE50-MD dogs compared to those of WT dogs

### Right ventricular dimensions and systolic function

Right ventricular diastolic volume was only assessed using CMR. Like LV dimensions, RVEDVI was significantly smaller in DE50-MD dogs compared to age-matched WT dogs, from 3 months of age (p=0.001). Weight-normalised TAPSE (TAPSEn and TAPSE:Ao) and RS’ (RS’n) were considered surrogate markers for longitudinal RV function. Values for RS’n were similar for DE50-MD and WT dogs (p=0.157). Although there was only a trend for TAPSEn to be lower in DE50-MD dogs, (p=0.054) TAPSE:Ao was significantly lower compared to WT controls (p=0.044). Values for TAPSEn and TAPSE:Ao, but not RS’n changed with age (p=0.001, p=0.015 and p=0.619 respectively).

### Relationship between left and right cardiac structure and function

Results of the Pearson’s correlation analyses revealed strong positive associations between LV and RV EDV (WT r=0.91. DE50-MD: r=0.89, p<0.0001 for both) and a moderate positive association between LV and RV EF, that was slightly stronger for WT than DE50MD dogs (WT: r=0.68. DE50-MD: r=0.59 p<0.0001 for both)). Results of LMM analysis revealed a highly significant association between RVEDVI and RV EF on LVEDVI and LV EF respectively in both genotypes, (p<0.001 for all). Results are summarised in figure 9.

**Figure 9.**
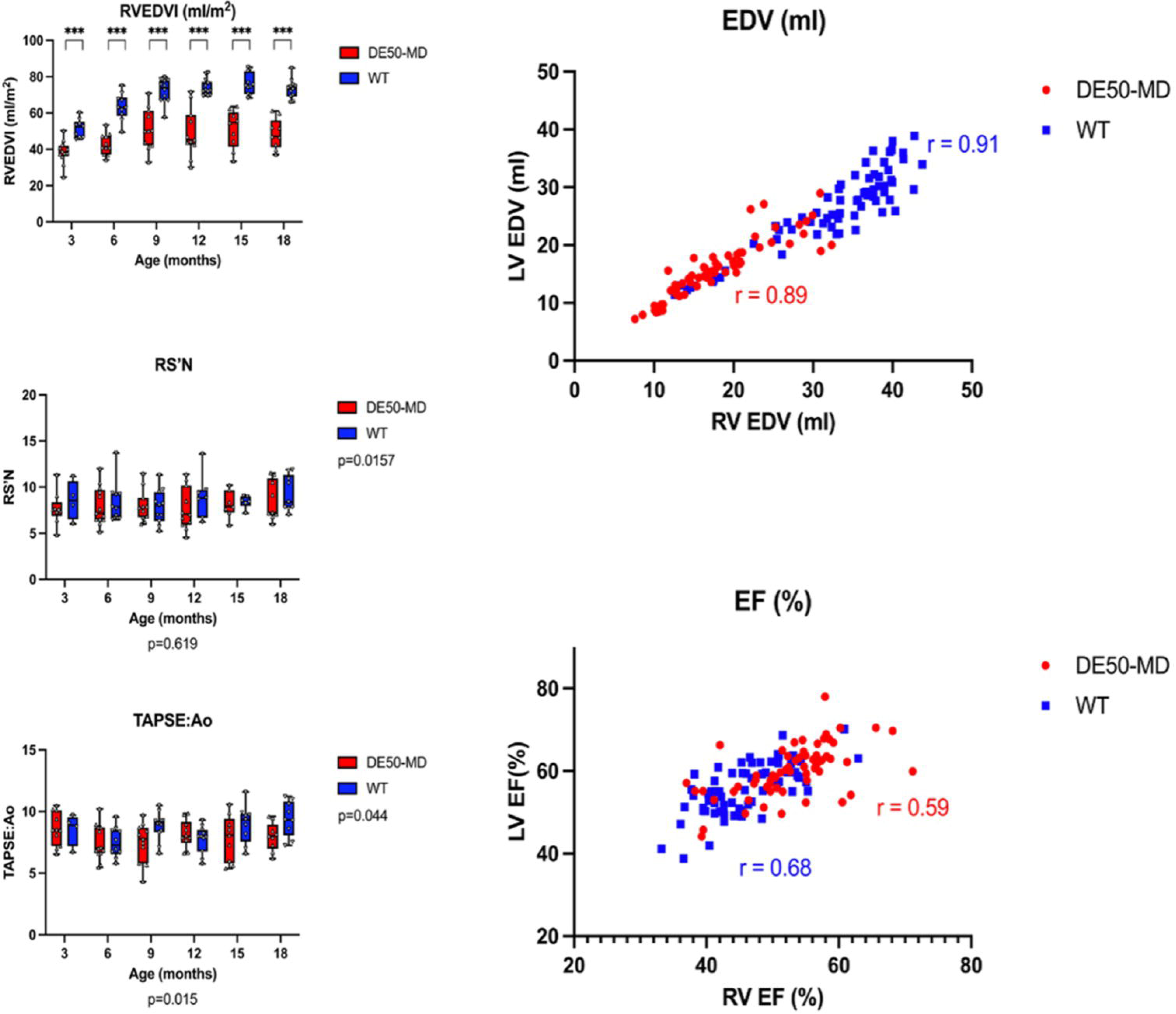
**A: Right ventricular (RV) diastolic volume and systolic function parameters using CMR and echocardiography respectively from age-matched wild type (WT, n=4-14) and DESO-MD (n=9-13) dogs, performed at 3-month intervals between 3 and 18 months of age.** Individual dogs are represented by staggered dots. Whiskers represent the minimum to maximum range. Horizontal lines represent median values. P values represent results of linear mixed model (LMM) analysis of repeated measures to explore effects of genotype (to the right of each graph) and age (below y axes). Where an interaction between genotype and age was established p values are not shown, instead significant differences, where present, for each age point studied are represented as follows: ***p<0.001. *Abbreviations: WT, Wzld Type. RVEDVI: RV end diastolic volume indexed to body surface area. RS’N, systolic longitudinal myocardial velocity measured using pulse waved tissue Doppler imaging at the basal RV free wall, indexed to body weight^0.233. TAPSE:Ao, Tricuspid annular plane systolic excursion indexed to aortic diameter.* DESO-MD dogs had highly significant reduction in RVEDVI at all ages studied. Measures of RV function were similar, apart from TAPSE:Ao, which had a tendency to be lower in DESO-MD, but the difference was only weakly significant in LMM analysis (p=0.044) **B: Association between right and left ventricular function and size.** Results of the Pearson’s correlation analysis revealed strong positive associations between LV and RV volume and moderate positive associations between LV and RV EF, that was slightly stronger for WT than DESO-MD dogs

### Late Gadolinium Enhancement

#### Study Population

A machine and software update to the 1·5 Tesla MRI machine permitted acquisition of LGE studies in dogs after March 2017. The study subpopulation is summarised in Table 6.

**Table 6.**
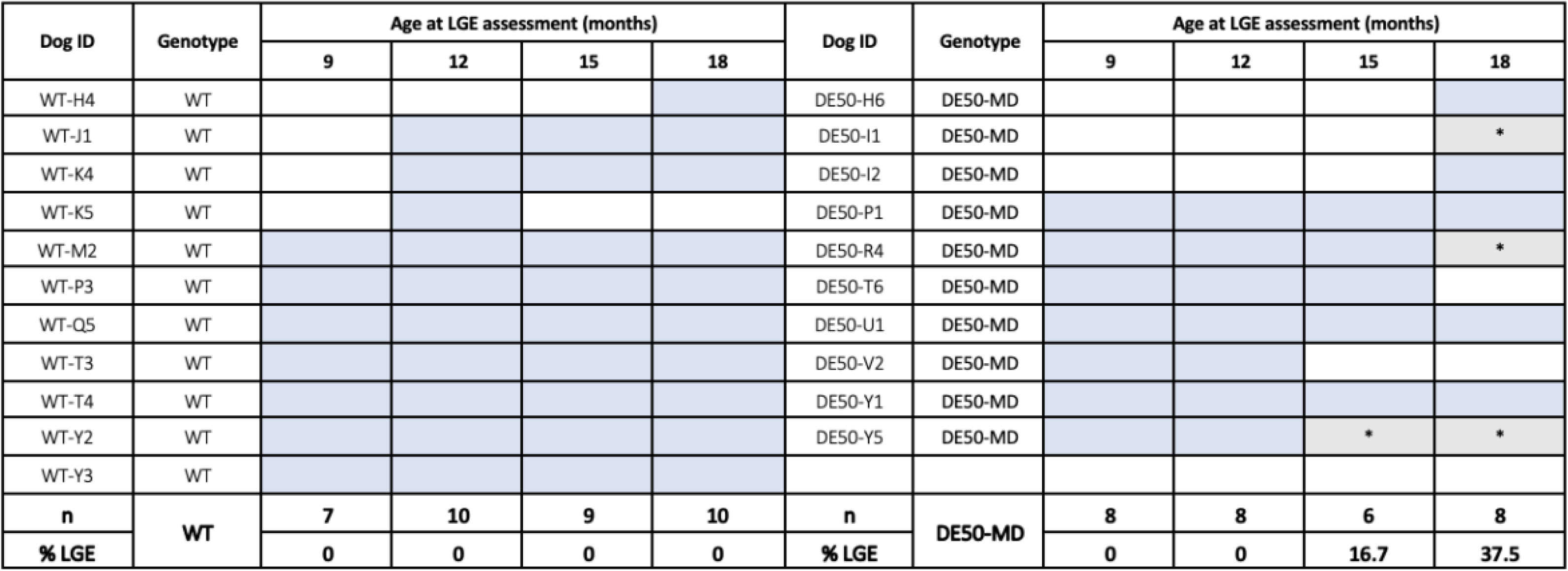
Summary of study population for late gadolinium enhancement (LGE) imaging in wild type (WT) and DE50-MD dogs. Highlighted blue cells represent dogs included in the study. Highlighted grey cells marked with x represent dogs in whom LGE was demonstrated. Approximately a third of DE50-MD dogs studied at 18 months of age showed evidence of LGE accumulation.

#### Results of Late Gadolinium enhancement imaging

Late gadolinium enhancement was identified in 3 dogs: 1/6 DE50-MD dogs studied at 15 months of age and 3/8 DE50-MD dogs studied at 18 months of age, (Table 5). In two dogs, LGE was noted first in a subepicardial distribution in the basal inferior, anterior and lateral wall segments. Of these latter dogs, LGE was present in the LV free wall at both 15 and 18 months, with additional involvement of the inferoseptal segment at 18 months. The third dog underwent LGE imaging for the first time at 18 months. In this dog, LGE imaging was only performed in LV mid-ventricular short axis and 4 chamber VLA views. Enhancement was present in the subepicardial to mid-myocardial wall in the mid, basal and apical inferior, anterior and lateral wall segments and with a patchy midmyocardial distribution in the septal wall. No WT dog was positive for LGE at any age. (Figure 10). The presence of LGE was not associated with overt systolic dysfunction (LV EF <55%) in any of the three LGE positive DE50-MD dogs.

**Figure 10:**
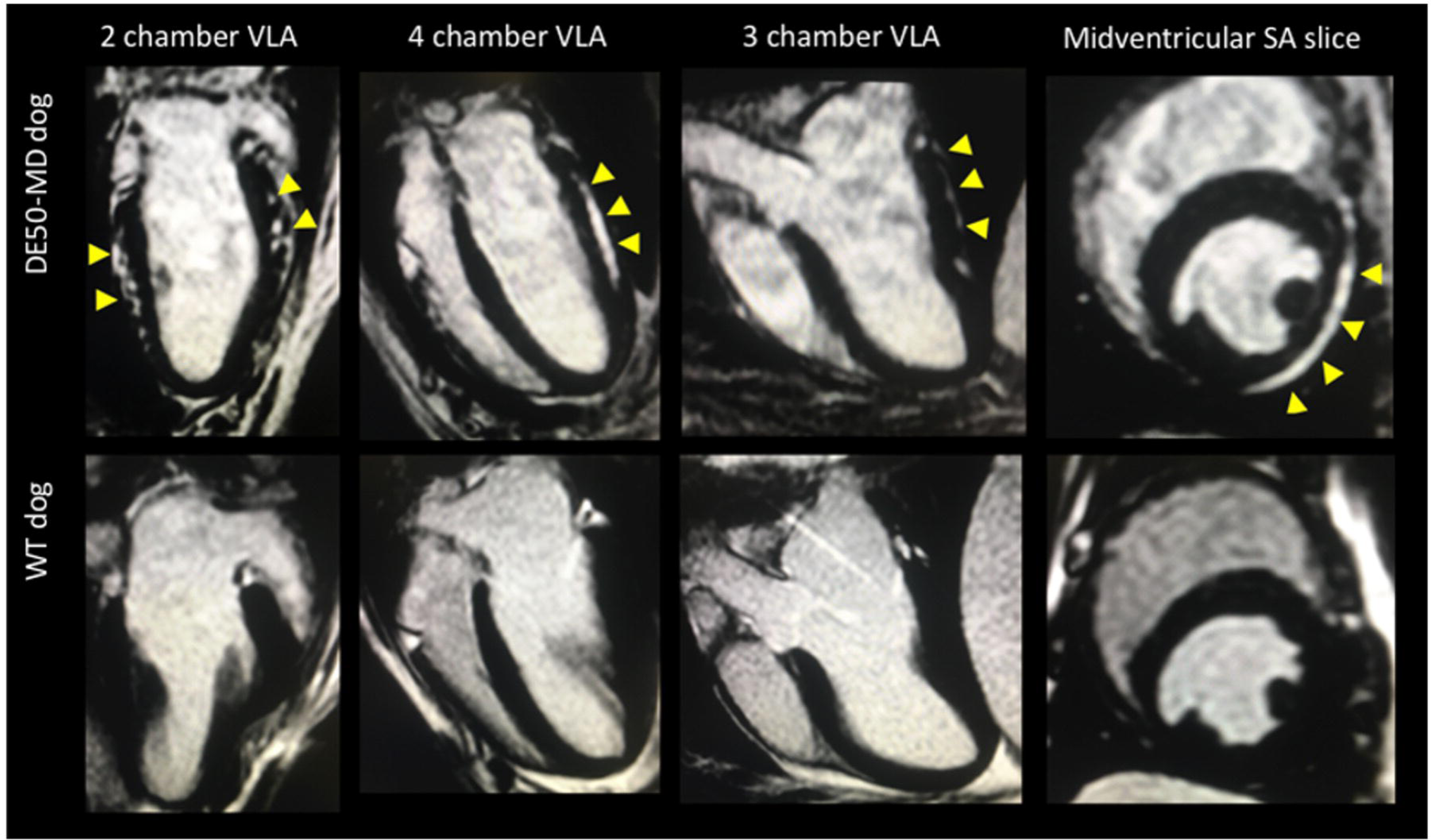
Inversion Recovery Turbo Field Echo images to demonstrate late gadolinium enhancement (LGE) imaging in an 18-month-old DESO-MD dog (upper panel) and age matched WT dog (lower panel). The corresponding 2, 4 and 3 chamber vertical long axis (VLA) views and midventricular short axis view (at the level of the papillary muscles) is show for each dog. Yellow arrow heads signify regions of LGE accumulation in the subepicardium of the basal to mid left ventricular free wall segments in the heart of the DESO-MD dog, compared to the completely nulled myocardium of the WT dog.

### Histopathology

Cardiac specimens were collected from 3 DE50-MD dogs and 3 WT dogs that underwent humane euthanasia within one week of their final CMR assessment at 18 months of age. One DE50-MD dog had demonstrated LGE, as detailed above. No LGE was identified in the other 2 dogs. The dog with LGE had extensive regions of myocardial degeneration and necrosis that were most severe in the basal and midventricular sections (but also present in apical regions). Where present, lesions were typically within the subepicardial and mid myocardium, becoming focally transmural in some sections. The basal sections contained predominately accumulations of mature adipocytes, termed “fatty infiltration”, (figure 11A, B) accompanied by mild fibrosis. In the midventricular sections, fatty infiltration and fibrosis were not as severe but instead there were large areas of myocyte necrosis with loss of normal myocardial architecture (figure 11 C,D). Mild to moderate fibrosis with minimal/no fatty infiltration was noted in septal lesions in basal, midventricular and apical sections. Similar lesions were not present in the WT or LGE negative DE50-MD dogs.

**Figure 11.**
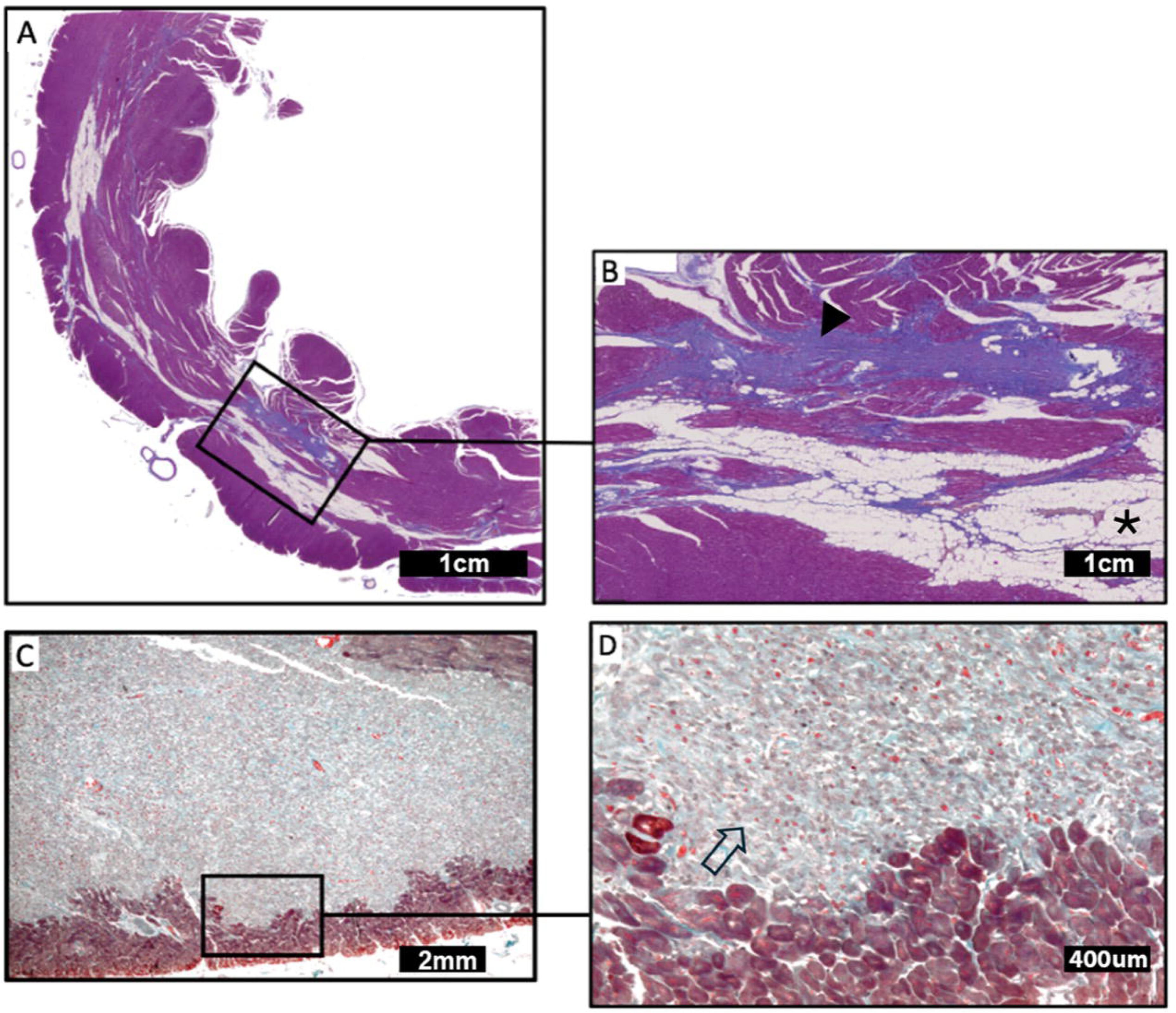
**A, B: Fibrofatty replacement in the basal left ventricular free wall myocardium from an LGE positive DESO-MD dog (A and B).** The sections demonstrate the presence of subepicardial to <u>rnidmyocardial</u> adipocyte infiltration (asterisk) and fibrosis (arrowhead). **C, D: Regions of myocyte loss in the left ventricular free wall myocardium from an LGE positive DESO­ MD dog.** The sections demonstrate the presence ofmyocyte necrosis and loss of normal myocardial architecture (arrow). Masson’s Trichrome stain; C and D are stained using a Gomori protocol (Images C and D courtesy of Dr Michael Ashworth and Dr Ciaran Hutchinson, Great Ormond Street Hospital for Children).

Purkinje fibre vacuolation was identified in both WT and DE50-MD dogs (figure 12). Where present in DE50-MD dogs, changes were not considered to exceed that of WT dogs or beyond that reported as an incidental finding in healthy laboratory beagles[50, 51]. Likewise, vascular wall thickening and narrowing of the vascular lumen were frequently seen in both WT and DE50-MD dogs, consistent with the normal variation in the appearance of mural arteries in the canine heart. At times the coronary arterial wall thickness appeared greater in individual vessels of DE50-MD dogs, but there was no evidence of medial necrosis, disruption of the internal elastic lamina or significant intimal hypertrophy[52] to support significant intramural coronary artery vascular remodelling (figure 12).

**Figure 12:**
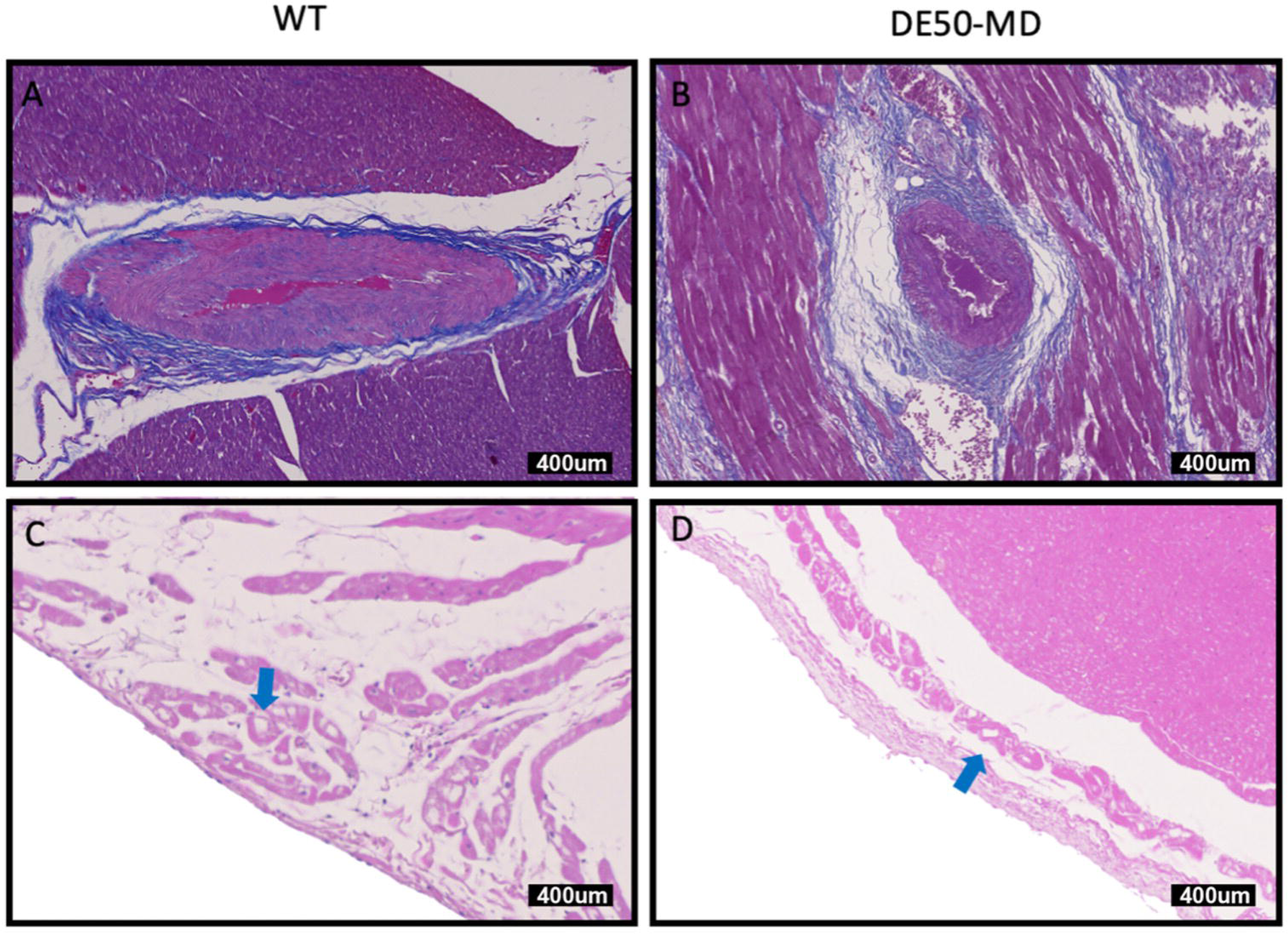
Left ventricular myocardium of 18-month-old WT (A and C) and DESO-MD (B and D) dogs. Figures A and B demonstrate intramural coronary vasculature, stained with Masson’s Trichrome. Note the thickness of the tunica media that is not considered consistent with vascular remodelling in either dog. Figures C and D demonstrate Purkinje fibres stained with haematoxylin and eosin (exhibiting a similar degree of Purkinje fibre vacuolation).

### Cardiac biomarkers

#### Study population

Cardiac troponin I was measured in samples from 5 WT and 8 DE50-MD dogs. NT-proBNP was measured in 14 WT dogs and 20 DE50-MD dogs at three-month intervals between 3 and 18 months of age. As for the imaging study, not all animals were included at each time point (See supplementary table 6a and 6b). Cardiac troponin I was increased in DE50-MD dogs compared to WT dogs with large spikes in values present in 4 DE50-MD dogs; DE50-R4, -U1, -Z6 at 18 months of age and DE50-Y5 at 12 months (figure 13A). Removal of these outlier data and natural logarithmic transformation of CtnI (lncTnI) was required to achieve normalisation of model residuals Values for lncTnI significantly increased with age (p=0.012) in both groups but was significantly increased in DE50-MD dogs at all time points (p<0.001).

**Figure 13:**
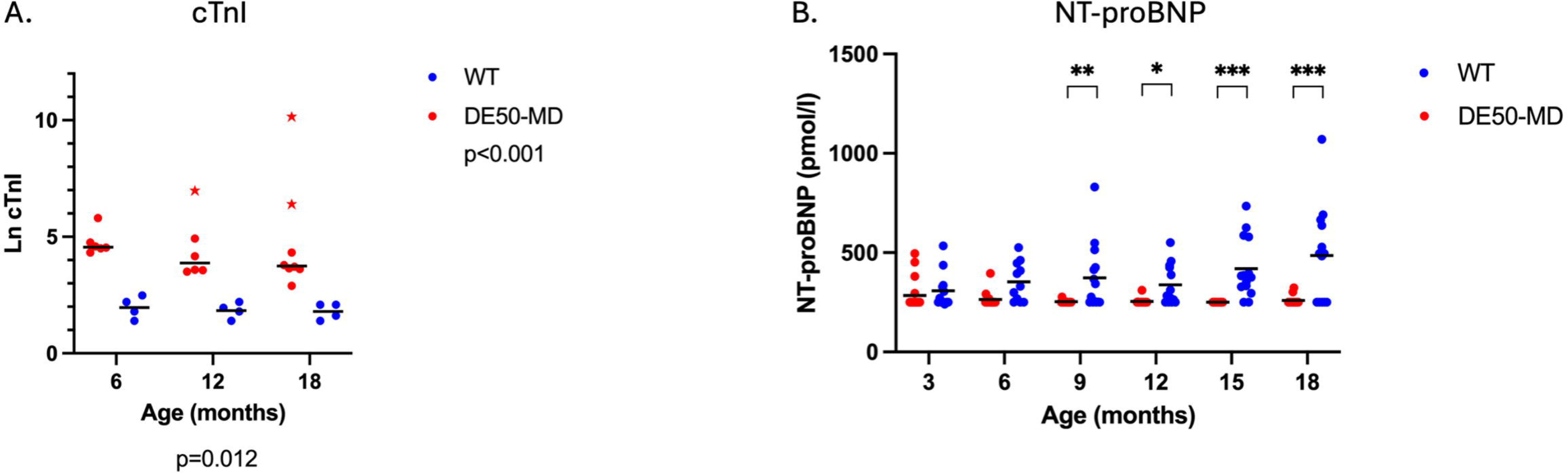
**A) Serum cardiac troponin I (cTnl) concentration in WT dogs (blue, n=5) and DESO-MD dogs, (n=lO) aged 6 to 18 months.** Each dot represents an individual dog. A horizontal line is placed at the mean value. Individual dogs represented by a star had severe transient elevation in serum cTnI, considered to be outliers in the clinical data. Cardiac troponin I concentration decreased with age but was consistently increased in DE50-MD dogs (p<0.001). **B) Plasma NT-proBNP concentration** in **WT (blue, n=14) and DESO-MD (red n=20) dogs.** The lower limit of detection of the assay is 250pmol/l. Each dot represents an individual dog. A horizontal line is placed at the mean value. Plasma NT-proBNP concentration was significantly lower from DE50-MD dogs than WT dogs from 9 months of age. (*p<0.5, **p<0.01, ***p<0.001, overall interaction between age and genotype, p<0.001).

When evaluating the biomarker NT-proBNP, statistical conformation of normality of model residuals could not be achieved, even after attempts to transform the data, likely due to the large number of data points that were below the threshold of the assay and so assigned the same value (250pg/ml). Direct visualization of residual histogram data confirmed reasonable symmetry over residual data, and no obvious outliers or skewness and so the statistical model was retained. A highly significant positive interaction between genotype and age was identified for NT-proBNP data (p<0.001; figure 13B), with values increasing with age in WT dogs (but not DE50-MD dogs) and being significantly lower in DE50-MD dogs from 9 months of age.

## Discussion

These data represent the first descriptions of the early structural and functional cardiac phenotype of the DE50-MD canine model of DMD. They confirm that the model shares preclinical features of DMD-related cardiomyopathy recognised in people and of growing GRMD dogs, the hitherto most widely used large animal model of the disease.[25, 26, 53]

### Left cardiac dimensions and geometry

The most striking feature of the DE50-MD cardiac phenotype is that both weight-adjusted LV dimensions and mass are smaller in DE50-MD dogs than age-matched littermate WT dogs. This disparity becomes greater as the dogs mature. In a recent longitudinal CMR study, Batra *et al*[8] demonstrated similar findings of reduced LVEDVI in children with DMD, highlighting that values were below the expected range for healthy children. The study demonstrated an initial decline in LVEDVI prior to subsequent dilation. The authors proposed that this was due to a combination of cardiac hypotrophy/atrophy and diastolic dysfunction.[8]

Our data are in accordance with the findings of several studies of young children,[54–58] and GRMD[59, 60] and mdx mouse[22] models, but are contradictory to earlier DMD echocardiographic studies, in which no significant difference in LV dimensions could be found between DMD boys and weight- or age-matched comparisons with people with normal systolic function.[61, 62] The apparently conflicting evidence likely reflects differences in study design and patient baseline demographics or small sample sizes in earlier studies. Different imaging modalities have been used to estimate LV volumes, including 2D- and 3D echocardiography and CMR. Two-dimensional echocardiographic studies have used a variety of methods to quantify volumes, but all make assumptions regarding the LV geometry, which itself might be altered in DMD.[57] To further add to the confusion, there is a lack of consensus amongst studies regarding how best to account for the impact of patient age, body size and activity levels on cardiac dimensions in studies of growing healthy children (or those with DMD). Control subjects in studies of DMD boys have included healthy subjects matched according to sex, age and/or body surface area,[61, 63] age-matched non-ambulatory patients[54] age-and weight matched patients with other non-ischaemic cardiomyopathies[64] and, increasingly, calculated “z-scores”, (in which an expected-normal value for volume or mass is calculated according to that patient’s height, body-surface area or lean body mass).[65, 66]

Interpretation of cardiac dimensions in growing dogs in similarly challenging. Growth rates vary according to breed and sex and there is a significant influence of age on echocardiographic parameters.[67] In an attempt to mitigate the influence of body weight of growing age-matched WT and DE50-MD dogs, we adjusted values for LV volumes to BSA, and LV diameters according to both weight-based allometric scaling methods[39, 68–70] [78][22] and alternative scaling methods. The latter included indexing LV and LA diameters to aortic size and LV mass to the length of the femur, and the length of the fifth lumbar vertebrae. The smaller left chamber size of DE50-MD dogs remained highly significant regardless of the method used, providing convincing evidence that the smaller hearts of DE50-MD dogs represent a genuine finding. The low cardiac mass and ventricular volumes are mirrored by low NT-proBNP results in DE50-MD dogs. Similarly, in a study comparing DMD patients with left ventricular systolic dysfunction and non-dystrophic dilated cardiomyopathy patients, showed that the DMD patients had smaller left ventricles and lower serum NT-proBNP concentration than the dilated cardiomyopathy patients. [71]While serum NT-proBNP concentration has been found to be predictive of mortality in DMD patients,[72] measurement of natriuretic peptides poorly identifies early cardiac dysfunction in DMD patients.[73, 74].

A proposed explanation for the smaller hearts of DMD patients is that these changes represent an adaptive response (physiological “hypotrophy”) to the reduced activity levels of dystrophin-deficient patients.[8, 56, 64, 66, 75–77] The heart undergoes functional and morphological adaptation to meet increased metabolic demands of repetitive exercise. This exercise-induced cardiac remodelling (EICR) leads to development of an “athlete’s heart” phenotype and occurs in adults[78] and children[79, 80] undertaking regular strenuous exercise. Similarly, the heart undergoes rapid reverse remodelling just weeks after detraining/deconditioning.[81] DE50-MD dogs show an early skeletal muscle phenotype [33, 82] and work performed by our group[83] has confirmed reduced activity levels in affected dogs when compared to littermate WT controls, (*Karimjee, K, R Piercy et al, unpublished data)*. In dogs, the athlete’s heart phenotype can manifest as different patterns of remodelling, reflecting breed variation and differences in type and duration of exercise/activity levels. In so-called “athletic breeds”, (such as racing greyhounds[84] and English Setters[85]) increased activity/exercise training is also associated with increased LV diastolic wall thickness and LV dimensions, compared to breed-matched, inactive controls. This suggests that these features are not solely related to genetic or epigenetic factors. As in people, exercise training in dogs leads to plasma volume expansion; in dogs by ∼15-30%.[86, 87] and early experimental studies in beagle and cross-bred dogs have demonstrated EICR to occur within as little as 5 to 12 weeks of treadmill exercise training,.[88, 89] On consideration of these findings, it is feasible that an opposite response to reduced activity could explain the reduced LV mass and ventricular volumes in DE50-MD dogs.

An alternative cause for cardiac hypotrophy however is reduced cardiac growth due to dysregulation of the cardiac signalling pathways regulating physiological hypertrophy or cardiac atrophy. Physiological growth during neonatal development is dependent on tight regulation of adaptive responses to mechanical stressors. Dystrophin links the cytoskeleton to the extracellular matrix via the dystrophin-glycoprotein complex (DGC). This adhesion complex interacts with integrin-based focal adhesion molecules within the costamere and with stress active filaments and plays a crucial role in mechanotransduction, i.e. the conversion of biomechanical forces into biochemical and/or epigenetic modulations that serve to modulate gene expression and cell behaviour.[90] It is therefore plausible that disruption of cardiac specific interactions between dystrophin and cytoskeletal proteins could impact physiological growth pathways.

To better understand the observed differences in the cardiac dimensions, we sought to characterise the pattern of cardiac remodelling in growing DE50-MD dogs. Consideration of LVMi, LVRI and RWT usefully distinguishes between different patterns of remodelling in people,[78, 91–93] (figure 6). Exercise-induced remodelling should not significantly impact LVRI or RWT due to concomitant increases in LV mass and volume[93, 94] although the morphometric adaptions encountered in EICR vary according to activity.[78, 95, 96] Endurance training is associated with physiological cardiac hypertrophy alongside augmentation of plasma volume that typically results in biventricular ventricular dilation (eccentric remodelling/hypertrophy). Strength training typically results in concentric hypertrophy, with disproportionate increase in LV mass to overcome increases in systemic blood pressure but with a minimal requirement to increase in cardiac output.[95, 97] CMR-derived LVRI values reported in healthy adults are higher than for those reported in healthy paediatric populations (1.03 ± 0.12 vs 0.68 - 0.91),[58, 98, 99] (which is in concordance with the influence of age on LVRI values for dogs in this study).

The increased LVRI found in DE50-MD dogs is in agreement with a recent CMR study of DMD boys.[58] Increased LVRI, in isolation, might be supportive of concentric hypertrophy or remodelling, which is interesting as early studies of DMD patients reported an early “hypertrophic cardiomyopathy” phenotype due to the presence of reduced LV chamber dimensions.[100]The DMD boys in the study by Mazur *et al* had LVMi and LVRI values that were within/below reported reference ranges for children and so the authors concluded that the geometric pattern was not consistent with LV hypertrophy. Variation of LVRI in health and disease is hitherto not described in dogs, in which characterisation of LV geometry typically relies on two dimensional assessments: i.e. LV sphericity and RWT. The canine athletic cardiac phenotype is typically characterised by an increase in RWT in dogs.[85, 101] Reversal/lack of EICR would be expected to result in a balanced or reduced LVRI and RWT alongside reduced LV Mass. Therefore, the geometric changes in DE50-MD dogs, i.e. increased LVRI, increased RWT but reduced LVMi, do not support isolated physiological hypotrophy. In the DE50-MD dogs, these findings were considered supportive of a disproportionately low diastolic LV volume in DE50-MD dogs that was further reflected by reduced sphericity of the DE50-MD LV.

Increased RWT and LVRI could signify pseudohypertrophy, (due to reduced preload/reduced venous return) or a state of maintained partial contraction during diastole. The latter would seem plausible, given known abnormalities in cardiomyocyte calcium-handling and diastolic dysfunction implicit to dystrophic cardiomyopathy.[57, 61] Diastolic dysfunction is detectable prior to systolic dysfunction in people[102] and in GRMD dogs.[29, 41, 42] There are challenges associated with assessment of diastolic function *in vivo*, however, as changes in loading conditions, (e.g. heart rate, preload and afterload), influence diastolic assessment. While there was only subtle evidence of diastolic dysfunction in the DE50-MD dogs, (attenuation of early diastolic wall motion of the left ventricular free wall), the state of tonic contraction is more credible than a state of reduced preload given the relative increase in LA size in comparison to the LV diastolic diameter encountered in affected dogs. Recent work on ex-vivo GRMD myocardial preparations has demonstrated transmural differences in myofilament structure and interactions associated with altered calcium sensitivity in the dystrophic myocardium.[103] Additional investigations in the DE50-MD dog using isolated myocardial preparations alongside detailed histomorphometric evaluations of the myocardium could provide additional information and would be an interesting focus of additional study in this model. The reduced cardiac dimensions in this model and in people are of particular importance in the setting of clinical trials as subsequent cardiac chamber enlargement in treated DMD patients could be considered evidence of cardiomyopathy progression (lack of treatment benefit). However, “normalisation” of cardiac dimensions to that of an untreated, healthy control population, could actually be a secondary indicator of improved skeletal muscle function or reflect genuine improvement of the cardiac phenotype.

### Left ventricular function

The preserved global systolic function in DE50-MD dogs (both on conscious TTE assessments and CMR evaluation under general anaesthesia) is unsurprising given the age range of the dogs studied. Overt systolic dysfunction is not typically apparent in untreated DMD boys until their second decade, (median onset of 13-16 years of age[104–107] being a relatively late manifestation of DMD-related cardiomyopathy). GRMD dogs develop overt global LV systolic dysfunction that mimics the human phenotype,[29, 41, 59, 108, 109] but this also occurs late:[29, 60] though the exact age of onset in the GRMD model is inconsistent amongst studies from separate colonies; between 30 and 40 months of age in one study,[29] but as young as 24 months according to the study from the colony from Centre d’Elevage du Domaine des Souches.[109] The disparity between the reports is likely related to the different genetic backgrounds of the two colonies and/or environmental factors. Similarly, longitudinal evaluation of the Japanese beagle crossed with GRMD (CXMDj) model in one study did not show evidence of overt systolic dysfunction (based on FS measurement) in any dog up to the age of 24 months.[110] It is therefore highly likely that the DE50-MD dogs followed in this study were too young to show detectable evidence of overt systolic dysfunction.

Tissue Doppler imaging was incorporated into this study to evaluate longitudinal myocardial motion at the basal mitral annulus. Although studies reporting the application of TDI imaging to evaluate DMD-related cardiomyopathy are limited to cross sectional studies in small numbers of patients, abnormalities of myocardial motion have consistently been identified in preclinical disease, prior to onset of global systolic dysfunction.[111–115]. In children with DMD, radial and longitudinal early diastolic and systolic velocities first appear reduced in the inferolateral and basal,[111, 113, 115, 116] mid and apical[111] anterolateral walls segments. These techniques could therefore be useful to expose early abnormalities in wall motion prior to onset of overt global LV dysfunction, providing an opportunity for early identification of systolic and diastolic functional derangements. Although longitudinal systolic velocity was preserved in DE50-MD dogs, early diastolic velocities were reduced in the basal inferolateral wall of DE50-MD dogs supporting early regional diastolic dysfunction.

This study used advanced echocardiographic techniques (STE and TDI imaging) to explore differences in deformation parameters between DE50-MD and healthy WT, littermate, control dogs. Consideration of isolated, (radial, longitudinal and circumferential) deformation planes in affected children show promise in exposing early markers of myocardial dysfunction, prior to onset of overt global dysfunction. This is desirable since early identification of DMD patients with a high risk of imminent progression of cardiomyopathy could help to optimise therapeutic interventions; this could have important prognostic significance later in life. Likewise, early indicators of functional decline in animal models could provide early biomarkers of disease progression in clinical trials. The findings from the colour TDI analysis are identical to those described in the GRMD dog model[41, 42]. Both systolic and diastolic radial MVGs were reduced in the LV free wall of DE50-MD dogs, principally due to lower endocardial radial velocities. Lower systolic [112, 113] and early diastolic ([111–113]) radial MVGs have also been reported in children with DMD prior to onset of global systolic dysfunction. Measurement of early diastolic MVGs is reported to be more sensitive than mitral inflow velocities to detect early aberrations in diastolic function in DMD cardiomyopathy and, in one small study, was to be predictive of worse outcome in young, asymptomatic boys with DMD without conventional echocardiographic signs of a DCM phenotype[112] The abnormality in early diastolic MVGs was more apparent than the systolic MVGs in the DE50-MD dogs in this study, recapitulating findings in children.[113] Measurement of inferolateral radial MVGs is similarly useful to identify early functional derangements in young DE50-MD dogs.

Average peak myocardial systolic strain was preserved in circumferential and longitudinal planes in DE50-MD dogs. There was tendency for lower average longitudinal peak systolic strain, but the difference was not statistically significant in the LMM. DE50-MD dogs showed hypercontractile circumferential peak systolic strain at early ages that normalised by 18 months of age. Li *et al* [117] reported similar findings in young mdx mice prior to subsequent decline in circumferential strain and, in the same study, increased myocardial twist/torsion at a young age. This study mirrored the results of the study by Miyazaki *et al*, [118] that identified a similar increase in LV rotation in DMD boys using STE. The data presented here failed to demonstrate depressed circumferential peak systolic strain in DE50-MD dogs at the young ages studied, which is comparable to longitudinal studies of GRMD dogs: Guo *et al*[60] reported that circumferential strain was unremarkable in DMD affected puppies up to 12 months of age compared to WT littermates, and longitudinal comparison between GRMD young adult dogs of 10 and 22 months of age did not identify significant decline in circumferential systolic strain between these ages.[29] Likewise, abnormalities of circumferential strain were not identified in the beagle crossed GRMD model of muscular dystrophy in Japan (CXMDJ) dogs aged 24.5months (± 15.3 months)[119] The findings from a separate colony of GRMD dogs in France (supplied by Centre d’Elevage du Domaine des Souches, France) are contradictory, however, since composite circumferential strain values were noted to decline over the first two years of life in these GRMD dogs compared to age-matched controls, and longitudinal peak strain depression and reduced twist was apparent from only 2 months of age[109] further highlighting differences in the rate of progression of cardiomyopathy between different GRMD colonies.

Information in the literature regarding average (from the midventricular level) or global (averaged segmental results from apical, mid and basal levels of the LV) circumferential strain in DMD boys using STE or CMR-tagging techniques appears consistent in its ability to stratify changes very early in life. Depressed circumferential strain values are apparent in DMD boys before 10 years of age, [120–122] and in patients prior to global systolic dysfunction.[121–126] The inconsistent results from animal studies could reflect genuine variation in the phenotypic progression of dystrophic cardiomyopathy between species but caution should be taken prior to drawing this conclusion due to relatively small sample numbers recruited in animal studies, which could be underpowered for demonstration of small but clinically relevant variation in systolic strain parameters, (absolute values of strain often differ by less than 5% between healthy and DMD patients).

Longitudinal peak systolic strain was attenuated in the apical segments of the DE50-MD dogs, alongside loss of the expected basal to apical gradient. This change was present in the affected dogs from 9 months of age, (the earliest age at which strain was analysed in this study) but was not sufficient to impact average peak LS measurements. The apical segments contribute substantially to ejection[127, 128] and have also been identified as a site of early decline of longitudinal strain values (measured using STE) in young DMD boys[124, 126] as well as in people with non-ischaemic DCM,[129–131] dogs with tachycardia-induced models of cardiomyopathy[132] and naturally occurring DCM.[133] A suggested mechanism for the preferential preservation of basal and mid-ventricular segments is that they are protected from increases in wall stress, and therefore subsequent remodelling, by the mitral annulus and papillary muscles.[132] Further, the apex contains the greatest proportion of longitudinally orientated fibres and therefore might be more vulnerable to deterioration in longitudinal strain analysis. In concordance with studies of DMD boys, these data support the conclusion that assessment of apical longitudinal strain could be useful as an early indicator of myocardial dysfunction in this model.

There was similar heterogeneity in peak systolic circumferential strain amongst individual midventricular segments. This heterogenicity was exaggerated in DE50-MD dogs compared to WT dogs, with peak systolic circumferential strain measurements identifying regional circumferential hypercontractility in some wall segments of the affected dogs. Assuming that there is minimal cardiac compression, circumferential, longitudinal and radial deformations are complementary for maintenance of cardiac ejection.[134] A plausible explanation for segmental circumferential hypercontractility is that this augmented segmental deformation compensates for depressed longitudinal peak systolic strain and reduced radial deformation, (supported by reduced radial myocardial velocity gradients in this study) in other segments. Studies of DMD boys that have attempted to identify segments showing the earliest decline in circumferential strain have yielded disparate results: using STE, the segments most commonly identified as having reduced circumferential strain are the inferior[135] and the anteroseptal[120, 124, 125, 136] wall segments, whilst CMR tagging studies have suggested that the lateral wall segments show the greatest decline in circumferential strain compared with healthy, age-matched boys.[8, 135, 136] This apparent lack of agreement likely reflects the different ages and severities of cardiomyopathy in the studied populations, confounding factors such as ambulatory status or the use of cardioprotective drugs, and the different methods used for strain analysis (i.e. CMR versus STE). The lack of consensus limits the suitability of specific segmental peak strain values as an early biomarker of the disease but does highlight the widespread myocardial damage that occurs within the left ventricle prior to detection of global systolic dysfunction.[126]

### Right ventricular geometry and function

The DE50-MD dogs had reduced RV dimensions and showed a tendency for reduced RV longitudinal function, (based on TAPSE) compared with age-matched WT controls. These data represent the first studies of RV geometry and function in a canine model of DMD.

Fewer studies report the progression of the RV phenotype in DMD than that of the LV. This probably, in part, relates to the challenges faced when assessing RV dimensions and function due to the complex geometry of this chamber, challenges in standardising echocardiographic windows, the interactive impact of LV volume on RV geometry, the high level of trabeculation in the RV and thin ventricular walls. With emergence of advanced imaging techniques, our understanding of RV function and its importance in cardiovascular disease has increased. Imaging with CMR is the reference standard for imaging the RV in people[137–139] and dogs,[140–142] and has been pivotal for improved characterisation of RV pathology in DMD boys over the past 5 to 10 years.[143–146] The lower RVEDVI of DE50-MD dogs is also a consistent finding in studies of DMD boys.[143–146] Right ventricular volume correlates well with LVEDVI in both species and, in DMD patients, is associated with worsening pulmonary function.[143] The mechanism behind reduced RVEDVI is yet to be confirmed, but it is suggested, as with left cardiac dimensions, to be related to reduced exercise/activity levels in DMD patients. [144] Other potential aetiologies likely parallel those discussed above regarding reduced left cardiac dimensions in dystrophin-deficient hearts.

The tendency for reduced TAPSE in DE50-MD dogs is challenging to interpret. Values for TAPSE are also lower in DMD and have been proposed to be a more sensitive measure of RV function than RV EF in this disease.[126, 145] As a linear measure of RV annular displacement, TAPSE is influenced by patient and/or cardiac dimension, therefore interpretation of results requires adjustment according to patient/cardiac size.[39, 147] This is complicated in paediatric populations due to the non-linear association between TAPSE and BSA identified in the period between birth and adolescence. This has led to the creation of age-dependent Z scores for TAPSE in children.[148] We attempted to adjust TAPSE measurements according to two established methods in dogs,[39, 147] but neither method has been validated in growing dogs and there was a significant effect of age on TAPSE despite their implementation. This complicates the interpretation or our values, and those reported from young DMD boys, in whom neither weight- nor age-adjusted values have been reported. Alternative methods with which to assess RV function reveal evidence of early RV dysynchrony that progresses alongside deterioration in LV function and CMR markers of fibrosis.[145, 146] Despite this, RV EF and deformation parameters appear to be comparatively well preserved in DMD patients compared to the LV.[125, 143, 145, 146, 149] Nevertheless, a recent abstract highlighted that RV dysfunction appears to be important prognostically, independent of LV dysfunction and requirement for mechanical ventilation, suggesting that RV involvement might be more important in dystrophic cardiomyopathy than previously appreciated.[150]

A proposed explanation for initial sparing of RV function in DMD is the reduced workload faced by the RV compared to the LV when pumping against the low pulmonary versus high systemic vascular resistance. As in DE50-MD dogs, LV and RV EF appear to be strongly related in DMD, but RVEF can be augmented in early disease as LV dysfunction progresses.[143] There is a strong association between decline in RV function and volumes and spirometry-derived measures of pulmonary function in DMD patients.[143] Right ventricular functional decline is likely more rapid for DMD patients with respiratory complications, due to increased RV afterload and progression of precapillary pulmonary hypertension secondary to restrictive respiratory disease and alveolar hypoinflation.[151] Consequently, sparing of RV function could have become more evident in recent years due to improvement in ventilatory care and therapeutics influencing respiratory function.

It is well established that mdx mice demonstrate a predominately RV early cardiac phenotype[152, 153] that is manifested by progressive fibrosis and biventricular blunted inotropic and lusiotropic response to dobutamine in invasive pressure-volume loop analyses.[152] This early RV involvement is thought to be related to the preferential diaphragm degeneration that results in respiratory dysfunction in mdx mice from 3 months of age,[154] with pulmonary hypertension proposed as being the major driving force behind the RV pathology.[153, 155] Work by our group has similarly identified early severe respiratory (diaphragmatic) muscle involvement in DE50-MD dogs.[33] The diaphragm develops progressive fibrosis and hypertrophy in the dogs at a young age[33] and increased expiratory diaphragmatic thickness, demonstratable from at least 3 months of age using thoracic ultrasound[156] and at post mortem.[33] Early severe degenerative changes are recognised in the diaphragm of GRMD[157, 158] and CXMDj[159] dogs but RV function has not been comprehensively described in these models, despite the severe fibrofatty degeneration in the RV of GRMD dogs (seen grossly from 10-12 months of age).[28] Restoration of diaphragmatic function in the mdx, (via isolated peptide-conjugated phosphorodiamidate morpholino oligomer treatment only of skeletal muscle) prevents the cardiac phenotype in mdx mice.[155] Further investigation is necessary to investigate the link between RV function and geometry and respiratory dysfunction in DE50-MD dogs, which could have important therapeutic implications.

### Myocardial injury and scarring in DE50-MD dogs

Cardiac troponin I values were increased in DE50-MD dogs, even at the youngest age assessed (3 months of age). Serum concentrations of cTnI decreased with age in DE50-MD and WT dogs, which mirrors that reported in growing children.[160] This likely reflects cardiomyocyte proliferation and cell turnover during physiological growth in post-natal hypertrophy and highlights the need for age-matched controls for consideration of cTnI in growing children and animals. Nevertheless, the increased values support a process of early myocardial injury that is evident in very young DE50-MD dogs, even prior to detection of scarring using cardiac MRI. The earliest age at which LGE was detected was 15 months of age in the DE50-MD dogs, however one LGE-positive dog was imaged for the first time at 18 months of age and so onset of LGE could not be determined in this animal. In the largest study of DMD patients undergoing LGE imaging, Hor *et al* (2013) demonstrated that LGE prevalence increased from 17% of patients <10 years to 34% of those aged 10-15 years and 59% of those >15 years-old,[161] The extent of LGE thus increases with patient age but, importantly, LGE can be identified in individuals from an early age, (before 10 years of age[161, 162]) prior to development of global systolic dysfunction. Similarly, LGE was identified in a small number of DE50-MD dogs as they approached maturity, despite preserved systolic function.

Late gadolinium enhancement was identified in the LV in approximately a third of the DE50-MD dogs. In the dogs exhibiting LGE, the segmental LGE distribution mirrored that of DMD-related cardiomyopathy. The LGE positive scar segments were in the inferior, inferolateral, or anterolateral basal and mid-cavity segments in two of the dogs; the third dog had a more widespread and transmural distribution. In people, transmurality is associated with more advanced fibrosis[161, 163–167] and it is likely that this dog had more advanced cardiomyopathy than the other two dogs. This can only be speculated however, since LGE imaging was not performed at earlier time points in this animal. Septal LGE was documented in 2/3 dogs. In DMD, septal involvement is typically only present in older patients with extensive transmural free wall involvement,[161, 167, 168] and has been associated with increased odds of exhibiting an EF of <55%.[164, 169] Our results recapitulate previous observations in the GRMD dog model, in which early septal involvement can also be demonstrated.[29] This disparity between species is unexplained but further investigation is warranted to evaluate the progression of LGE in DE50-MD dogs, with regards to its association with age, its distribution and impact on cardiac structure, function and electrophysiology.

We compared the results of LGE imaging to histological cardiac sections from a small number of dogs that had undergone CMR imaging immediately prior to euthanasia. Pathological lesions were demonstrated only in the LV myocardium of the dog that had exhibited LGE during post contrast CMR imaging. As described in GRMD dogs, lesions consisted of regions of myocyte injury, characterised by multifocal subepicardial to focally transmural myocyte degeneration and necrosis, with evidence of macrophage and fibroblast infiltration and islands of fibrosis and fatty infiltration.[28, 170] The lesions in this dog were extensive and provided a substantial contrast to the complete absence of myocardial lesions in the other two DE50-MD (and WT) dogs studied. These findings mirrored the phenotypic variation encountered during LGE imaging, highlighting the utility of the latter technique. In addition to the ventricular myocardial lesions identified, we also documented other changes reported to contribute to the dystrophic phenotype of the GRMD and CXMDj canine models.[28, 110, 170–174] We noted the presence of Purkinje fibre vacuolation in DE50-MD dogs, which was investigated as an early pathological finding in the CXMDj model. [171, 172] In the CXMDj model, vacuolation has been noted in association with accumulation and mislocalisation of the calcium-dependent protease-μ calpain at the Purkinje fibre subsarcolemma.[171] In our study, the subjective assessment by an experienced toxicological pathologist was that the extent of vacuolation encountered in the DE50-MD dogs did not exceed that expected as a frequent incidental finding of healthy laboratory beagles.[50, 175] Indeed, we were able to identify similar changes in age-matched WT comparisons. Comparison with results of the CXMDj model are challenging due to lack of a control WT population in these studies. Purkinje fibre vacuoles in CXMDj dogs were reportedly more irregular and extensive than the authors expected and electron microscopy identified severe disruption of myofibrils, suggesting that the vacuoles were the ruins of myofibrillar structures rather than being due to regions of myofiber sparsity or mislocalisation (as expected in healthy Purkinje fibres)[171]. Our findings question the significance of Purkinje vacuolation as a specific feature in canine DMD models. Similarly, intramural coronary artery medial thickening, reported in GRMD dogs, [28] was a prominent feature of both WT and DE50-MD dogs, with no evidence to support significant vascular remodelling in affected dogs.

There were several limitations to the present study. Due to the evident skeletal muscle phenotype in the DE50-MD dogs, the cardiologist acquiring echocardiographic and CMR studies could not be blinded to the phenotype of the dog which might have led to observation bias. Analysis of repeated measures was complicated by missing data points for various data. Reasons for loss of data included: lack of tolerance for restraint in (almost exclusively) juvenile WT dogs, (which is not an uncommon problem in young animals), poor acoustic windows in some DE50-MD dogs that impeded use of some echocardiographic techniques (principally TDI) and early necessary euthanasia of some dogs prior to the 18-month end point. The cause for early euthanasia was not considered related to deterioration in the cardiac function in any dog but exploration was limited by the variation in age at which dogs were euthanased, precluding age-matched comparison. Further characterisation of the strain distribution pattern in young DE50-MD dogs could be achieved through a more complete evaluation of circumferential and longitudinal peak strains in all myocardial segments. Exploration for LGE could only be performed in a small subgroup of dogs, which limits our ability to make firm conclusions regarding prevalence and age at which LGE becomes a feature in this model. Myocardial histology was only available for one dog exhibiting LGE at this young age. We did not attempt direct comparison between location and extent of LGE and lesions detectable during histological analysis for this dog since this dog was one of the earliest individuals in whom LGE imaging was attempted. We therefore lacked high quality images in all imaging planes to permit meaningful comparison. Based on these findings, we subsequently adopted a systematic, cross-sectional approach in a larger group of older dogs (reported separately). Differentiation between incidental “background” lesions and genuine pathology was made more difficult by the small number of dogs included in the study. This highlights the importance for age-matched healthy comparisons and observation by an experienced veterinary pathologist to interpret histological findings in experimental settings. Histological studies in a larger number of dogs over a greater age range could provide additional insight into whether any genuine pathology is present in the coronary vasculature and Purkinje fibres of DE50-MD dogs.

## Conclusions

These data demonstrate that DE50-MD dogs under 18 months of age have a cardiac phenotype that shares many characteristics of the early cardiomyopathy found in boys with DMD. Global LV and RV systolic function is preserved in DE50-MD dogs up to 18 months of age, which is similar to findings in CXMDj dogs[110, 176] and in contemporary reports of GRMD,[29, 60, 109] although the progression of the cardiac phenotype appears to be more benign than that previously been reported in some GRMD colonies.[108] Radial systolic and diastolic myocardial velocity gradients, (approximating radial strain rate) were depressed in DE50-MD dogs, in agreement with studies of preclinical GRMD dogs and in boys with DMD. Composite midventricular circumferential peak systolic strain values from DE50-MD dogs were hypercontractile between 9 and 15 months of age but normalised at 18 months of age. While composite longitudinal strain values were similar between DE50-MD and WT dogs, depressed segmental apical longitudinal systolic strain was present from at least 9 months of age, (as reported in young DMD boys) and was associated with loss of the expected apical to basal longitudinal strain gradient encountered in healthy people and WT control dogs. The LV dimensions in DE50-MD dogs are strikingly reduced compared to WT littermate comparisons, which is a variably reported, poorly characterised and possibly overlooked finding both in animal models and in people with DMD. These findings should be the subject of future investigation as they could be important in the early pathophysiology of DMD-related cardiomyopathy.[57] Approximately a third of DE50-MD dogs showed early evidence of LGE in the LV by 18 months of age that corresponded to regions of myocyte necrosis and fibrofatty scarring on histopathology. The distribution of LGE resembled that reported in the GRMD dog model, affecting the basal and mid ventricular LV free wall and variably, mild septal involvement. Onset of LGE was not associated with overt systolic dysfunction at the young ages studied. In summary, the data suggests that these animals represent a highly relevant model to study the earliest phenotypic features of cardiac dystrophin deficiency.

## Acknowledgements

The authors thank colleagues at the Royal Veterinary College for the many contributions that permitted these studies, and staff within the Biological Services Unit for excellent and compassionate care of the animals. Also, Dr Michael Ashworth and Dr Ciaran Hutchinson, Great Ormond Street Hospital for Children, for their kind assistance with the preparation and imaging of selected slides for histopathological assessment. Special thanks to Dr. Yu-Mei Chang for advice on statistical analysis.

## REFERENCES

1. Mendell JR, Shilling C, Leslie ND, et al (2012) Evidence-based path to newborn screening for duchenne muscular dystrophy. Ann Neurol 71:304–313

2. Moat SJ, Bradley DM, Salmon R, Clarke A, Hartley L (2013) Newborn bloodspot screening for Duchenne muscular dystrophy: 21 years experience in Wales (UK). Eur J Hum Genet 21:1049–1053

3. Birnkrant DJ, Ararat E, Mhanna MJ (2015) Cardiac phenotype determines survival in Duchenne muscular dystrophy. Pediatr Pulmonol 51:70–76

4. Frankel KA, Rosser RJ (1976) The pathology of the heart in progressive muscular dystrophy: Epimyocardial fibrosis. Hum Pathol 7:375–386

5. Kamdar F, Garry DJ (2016) Dystrophin-Deficient Cardiomyopathy. J Am Coll Cardiol 67:2533–2546

6. Nigro G, Comi LI, Politano L, Bain RJ (1990) The incidence and evolution of cardiomyopathy in Duchenne muscular dystrophy. Int J Cardiol 26:271–277

7. Chrzanowski SM, Darras BT, Rutkove SB (2019) The Value of Imaging and Composition-Based Biomarkers in Duchenne Muscular Dystrophy Clinical Trials. Neurotherapeutics 1–11

8. Batra A, Barnard AM, Lott DJ, et al (2022) Longitudinal changes in cardiac function in Duchenne muscular dystrophy population as measured by magnetic resonance imaging. Bmc Cardiovasc Disor 22:260

9. Lee S, Lee M, Hor KN (2021) The role of imaging in characterizing the cardiac natural history of Duchenne muscular dystrophy. Pediatr Pulm 56:766–781

10. Magrath P, Maforo N, Renella P, Nelson SF, Halnon N, Ennis DB (2018) Cardiac MRI biomarkers for Duchenne muscular dystrophy. Biomarkers in Medicine 12:1271–1289

11. Eagle M, Baudouin SV, Chandler C, Giddings DR, Bullock R, Bushby K (2002) Survival in Duchenne muscular dystrophy: improvements in life expectancy since 1967 and the impact of home nocturnal ventilation. Neuromuscul Disord 12:926–929

12. Kieny P, Chollet S, Delalande P, Fort ML, Magot A, Pereon Y, Verbe BP (2013) Evolution of life expectancy of patients with Duchenne muscular dystrophy at AFM Yolaine de Kepper centre between 1981 and 2011. Annals of Physical and Rehabilitation Medicine 56:443–454

13. Taglia A, Palladino A, Viggiano E, et al (2012) Improvement of survival in Duchenne Muscular Dystrophy: retrospective analysis of 835 patients. Acta Myol 31:1–5

14. Wang M, Birnkrant DJ, Super DM, Jacobs IB, Bahler RC (2018) Progressive left ventricular dysfunction and long-term outcomes in patients with Duchenne muscular dystrophy receiving cardiopulmonary therapies. Open Heart 5:e000783–9

15. Cheeran D, Khan S, Khera R, et al (2017) Predictors of Death in Adults With Duchenne Muscular Dystrophy-Associated Cardiomyopathy. J Am Heart Assoc 6:1–12

16. Punnoose AR, Kaltman JR, Pastor W, McCarter R, He J, Spurney CF (2016) Cardiac Disease Burden and Risk of Mortality in Hospitalized Muscular Dystrophy Patients. Pediatric Cardiology 37:1290–1296

17. Finsterer J, Stöllberger C (2007) Cardiac involvement determines the prognosis of Duchenne muscular dystrophy. Indian J Pediatr 74:209-author reply 210-2

18. D’Amario D, Amodeo A, Adorisio R, et al (2017) A current approach to heart failure in Duchenne muscular dystrophy. Heart 103:1770–1779

19. Sicinski P, Geng Y, Ryder-Cook AS, Barnard EA, Darlison MG, Barnard PJ (1989) The molecular basis of muscular dystrophy in the mdx mouse: a point mutation. Science 244:1578–1580

20. Chamberlain JS, Metzger J, Reyes M, Townsend D, Faulkner JA (2007) Dystrophin-deficient mdx mice display a reduced life span and are susceptible to spontaneous rhabdomyosarcoma. FASEB J 21:2195–2204

21. Quinlan JG, Hahn HS, Wong BL, Lorenz JN, Wenisch AS, Levin LS (2004) Evolution of the mdx mouse cardiomyopathy: physiological and morphological findings. Neuromuscular Disord 14:491–496

22. Erp CV, Loch D, Laws N, Trebbin A, Hoey AJ (2010) Timeline of cardiac dystrophy in 3-18-month-old MDX mice. Muscle Nerve 42:504–513

23. Stuckey DJ, Carr CA, Camelliti P, Tyler DJ, Davies KE, Clarke K (2012) In vivo MRI characterization of progressive cardiac dysfunction in the mdx mouse model of muscular dystrophy. PLoS ONE 7:e28569

24. Mokhtarian A, Lefaucheur JP, Even PC, Sebille A (1995) Effects of treadmill exercise and high-fat feeding on muscle degeneration in mdx mice at the time of weaning. Clin Sci 89:447–452

25. Kornegay JN, Bogan JR, Bogan DJ, et al (2012) Canine models of Duchenne muscular dystrophy and their use in therapeutic strategies. Mamm Genome 23:85–108

26. Wells DJ (2018) Tracking progress: an update on animal models for Duchenne muscular dystrophy. Dis Model Mech 11:

27. Kornegay JN (2017) The golden retriever model of Duchenne muscular dystrophy. Skelet Muscle 7:9

28. Schneider SM, Sansom GT, Guo L-J, Furuya S, Weeks BR, Kornegay JN (2022) Natural History of Histopathologic Changes in Cardiomyopathy of Golden Retriever Muscular Dystrophy. Frontiers Vet Sci 8:759585

29. Guo LJ, Soslow JH, Bettis AK, Nghiem PP, Cummings KJ, Lenox MW, Miller MW, Kornegay JN, Spurney CF (2019) Natural History of Cardiomyopathy in Adult Dogs With Golden Retriever Muscular Dystrophy. J Am Heart Assoc 8:121–29

30. Walmsley GL, Arechavala-Gomeza V, Fernandez-Fuente M, et al (2010) A duchenne muscular dystrophy gene hot spot mutation in dystrophin-deficient cavalier king charles spaniels is amenable to exon 51 skipping. PLoS ONE 5:e8647

31. Aartsma-Rus A, Fokkema I, Verschuuren J, Ginjaar I, Deutekom J van, Ommen G van, Dunnen JT den (2009) Theoretic applicability of antisense-mediated exon skipping for Duchenne muscular dystrophy mutations. Hum Mutat 30:293–299

32. Amoasii L, Hildyard JCW, Li H, et al (2018) Gene editing restores dystrophin expression in a canine model of Duchenne muscular dystrophy. Science 8:eaau1549-91

33. Hildyard JCW, Riddell DO, Harron RCM, Rawson F, Foster EMA, Massey C, Taylor-Brown F, Wells DJ, Piercy RJ (2022) The skeletal muscle phenotype of the DE50-MD dog model of Duchenne muscular dystrophy. Wellcome Open Res 7:238

34. Hildyard JCW, Foster EMA, Wells DJ, Piercy RJ (2022) Rapid histological quantification of muscle fibrosis and lysosomal activity using the HSB colour space. Biorxiv 2022.08.02.502489

35. Hornby N, Drees R, Wells DJ, Piercy RJ (2018) MRI evaluation of the pelvic limb and lumbar muscles of the deltaE50-MD dog model of Duchenne muscular dystrophy. Neuromuscular Disorders 28:S15

36. Riddell DO, Hildyard JCW, Harron RCM, Taylor-Brown F, Kornegay JN, Wells DJ, Piercy RJ (2023) Longitudinal assessment of skeletal muscle functional mechanics in the DE50-MD dog model of Duchenne Muscular Dystrophy. Dis Model Mech 16:dmm050395

37. Crawford AH, Hildyard JCW, Rushing SAM, Wells DJ, Diez-Leon M, Piercy RJ (2022) Validation of DE50-MD dogs as a model for the brain phenotype of Duchenne muscular dystrophy. Dis Model Mech 15:dmm049291

38. Lang RM, Badano LP, Mor-Avi V, et al (2015) Recommendations for cardiac chamber quantification by echocardiography in adults: an update from the American Society of Echocardiography and the European Association of Cardiovascular Imaging. European Heart Journal -Cardiovascular Imaging 16:233–270

39. Visser LC, Scansen BA, Brown NV, Schober KE, Bonagura JD (2015) Echocardiographic assessment of right ventricular systolic function in conscious healthy dogs following a single dose of pimobendan versus atenolol. J Vet Cardiol 17:161–172

40. Chetboul V, Carlos C, Blot S, Thibaud JL, Escriou C, Tissier R, Retortillo JL, Pouchelon J-L (2004) Tissue Doppler assessment of diastolic and systolic alterations of radial and longitudinal left ventricular motions in Golden Retrievers during the preclinical phase of cardiomyopathy associated with muscular dystrophy. Am J Vet Res 65:1335–1341

41. Chetboul V, Escriou C, Tessier D, Richard V, Pouchelon J-L, Thibault H, Lallemand F, Thuillez C, Blot S, Derumeaux G (2004) Tissue Doppler imaging detects early asymptomatic myocardial abnormalities in a dog model of Duchenne’s cardiomyopathy. Eur Heart J 25:1934–1939

42. Chetboul V, Carlos C, Blot S, Thibaud J-L, Escriou C, Tissier R, Retortillo JL, Pouchelon J-L (2004) Tissue Doppler assessment of diastolic and systolic alterations of radial and longitudinal left ventricular motions in Golden Retrievers during the preclinical phase of cardiomyopathy associated with muscular dystrophy. American Journal of Veterinary Research 65:1335–1341

43. Schulz-Menger J, Bluemke DA, Bremerich J, et al (2013) Standardized image interpretation and post processing in cardiovascular magnetic resonance: Society for Cardiovascular Magnetic Resonance (SCMR) Board of Trustees Task Force on Standardized Post Processing. J Cardiov Magn Reson 15:35

44. Vinnakota KC, Bassingthwaighte JB (2004) Myocardial density and composition: a basis for calculating intracellular metabolite concentrations. Am J Physiol-heart C 286:H1742– H1749

45. Cerqueira MD, Weissman NJ, Dilsizian V, Jacobs AK, Kaul S, Laskey WK, Pennell DJ, Rumberger JA, Ryan T, Verani MS (2002) Standardized Myocardial Segmentation and Nomenclature for Tomographic Imaging of the Heart. Circulation 105:539–542

46. Riddell DO, Hildyard JCW, Harron RCM, Wells DJ, Piercy RJ (2022) Longitudinal assessment of blood-borne musculoskeletal disease biomarkers in the DE50-MD dog model of Duchenne muscular dystrophy. Wellcome Open Res 6:354

47. Barthélémy I, Calmels N, Weiss RB, et al (2020) X-linked muscular dystrophy in a Labrador Retriever strain: phenotypic and molecular characterisation. Skelet Muscle 10:23

48. Thibaud J-L, Monnet A, Bertoldi D, Barthélémy I, Blot S, Carlier PG (2007) Characterization of dystrophic muscle in golden retriever muscular dystrophy dogs by nuclear magnetic resonance imaging. Neuromuscular Disord 17:575–584

49. Cizinauskas S, Jaggy A, Tipold A (2000) Long-term treatment of dogs with steroid-responsive meningitis-arteritis: clinical, laboratory and therapeutic results. J Small Anim Pract 41:295–301

50. Woicke J, Al-Haddawi MM, Bienvenu J-G, et al (2021) International Harmonization of Nomenclature and Diagnostic Criteria (INHAND): Nonproliferative and Proliferative Lesions of the Dog. Toxicol Pathol 49:5–109

51. Bodié K, Decker JH (2014) Incidental histopathological findings in hearts of control beagle dogs in toxicity studies. Toxicol Pathol 42:997–1003

52. Kohnken R, Weber A (2020) Characterization of Spontaneous Vascular Findings in the Papillary Muscles of Beagle Dogs. Toxicol Pathol 48:899–904

53. Ameen V, Robson LG (2010) Experimental models of duchenne muscular dystrophy: relationship with cardiovascular disease. Open Cardiovasc Med J 4:265–277

54. Goldberg SJ, Stern LZ, Feldman L, Sahn DJ, Allen HD, Valdes-Cruz LM (1983) Serial left ventricular wall measurements in Duchenne’s muscular dystrophy. J Am Coll Cardiol 2:136–142

55. Farah MG, Evans EB, Vignos PJ (1980) Echocardiographic evaluation of left ventricular function in Duchenne’s muscular dystrophy. Am J Med 69:248–254

56. Shapiro F, Sethna N, Colan S, Wohl ME, Specht L (1992) Spinal fusion in Duchenne muscular dystrophy: a multidisciplinary approach. Muscle Nerve 15:604–614

57. Su JA, Ramos-Platt L, Menteer J (2015) Left Ventricular Tonic Contraction as a Novel Biomarker of Cardiomyopathy in Duchenne Muscular Dystrophy. Pediatric Cardiology 37:678–685

58. Mazur W, Hor KN, Germann JT, et al (2011) Patterns of left ventricular remodeling in patients with Duchenne Muscular Dystrophy: a cardiac MRI study of ventricular geometry, global function, and strain. Int J Cardiovasc Imaging 28:99–107

59. Fine DM, Shin J-H, Yue Y, Volkmann D, Leach SB, Smith BF, McIntosh M, Duan D (2011) Age-matched comparison reveals early electrocardiography and echocardiography changes in dystrophin-deficient dogs. Neuromuscul Disord 21:453–461

60. Guo L-J, Nghiem PP, Bettis AK, Soslow JH, Spurney CF, Kornegay JN (2024) Early Natural History of Cardiomyopathy and Cardiac Stress Response in Young Dogs with Golden Retriever Muscular Dystrophy. bioRxiv 2024.08.19.608721

61. Ahmad M, Sanderson JE, Dubowitz V, Hallidie-Smith KA (1978) Echocardiographic assessment of left ventricular function in Duchenne’s muscular dystrophy. Br Heart J 40:734–740

62. Heymsfield SB, McNish T, Perkins JV, Felner JM (1978) Sequence of cardiac changes in Duchenne muscular dystrophy. Am Heart J 95:283–294

63. Heymsfield SB, McNish T, Perkins JV, Felner JM (1978) Sequence of cardiac changes in Duchenne muscular dystrophy. American Heart Journal 95:283–294

64. Khan S, Cheeran D, Garg S, Grodin J, Morlend R, Araj F, Amin A, Thibodeau J, Drazner M, Mammen P (2017) Cardiac atrophy: A novel mechanism for Duchenne muscular dystrophy (DMD)-associated cardiomyopathy. J Am Coll Cardiol 69:946

65. Su JB, Cazorla O, Blot S, et al (2012) Bradykinin restores left ventricular function, sarcomeric protein phosphorylation, and e/nNOS levels in dogs with Duchenne muscular dystrophy cardiomyopathy. Cardiovasc Res 95:86–96

66. Lee TH, Eun LY, Choi JY, Kwon HE, Lee Y-M, Kim HD, Kang S-W (2014) Myocardial atrophy in children with mitochondrial disease and Duchenne muscular dystrophy. Korean J Pediatr 57:232–8

67. Sisson D, Schaeffer D (1991) Changes in linear dimensions of the heart, relative to body weight, as measured by M-mode echocardiography in growing dogs. Am J Vet Res 52:1591– 1596

68. Cornell CC, Kittleson MD, Torre PD, Häggström J, Lombard CW, Pedersen HD, Vollmar A, Wey A (2004) Allometric scaling of M-mode cardiac measurements in normal adult dogs. J Vet Intern Med 18:311–321

69. Visser LC, Ciccozzi MM, Sintov DJ, Sharpe AN (2019) Echocardiographic quantitation of left heart size and function in 122 healthy dogs: A prospective study proposing reference intervals and assessing repeatability. J Vet Intern Med 33:1909–1920

70. Esser LC, Borkovec M, Bauer A, Häggström J, Wess G (2020) Left ventricular M-mode prediction intervals in 7651 dogs: Population-wide and selected breed-specific values. J Vet Intern Med 34:2242–2252

71. Demachi J, Kagaya Y, Watanabe J, et al (2004) Characteristics of the increase in plasma brain natriuretic peptide level in left ventricular systolic dysfunction, associated with muscular dystrophy in comparison with idiopathic dilated cardiomyopathy. Neuromuscular Disord 14:732–739

72. Soslow JH, Xu M, Slaughter JC, et al (2023) Cardiovascular Measures of All-Cause Mortality in Duchenne Muscular Dystrophy. Circ: Hear Fail 16:e010040

73. Mori K, Manabe T, Nii M, Hayabuchi Y, Kuroda Y, Tatara K (2002) Plasma Levels of Natriuretic Peptide and Echocardiographic Parameters in Patients with Duchenne’s Progressive Muscular Dystrophy. Pediatr Cardiol 23:160–166

74. Westrum SS van, Dekker L, Haan R de, Endert E, Ginjaar I, Visser M de, Kooi A van der (2013) Brain natriuretic peptide is not predictive of dilated cardiomyopathy in Becker and Duchenne muscular dystrophy patients and carriers. BMC Neurol 13:88

75. Matsuoka S, Ii K, Akita H, Tomimatsu H, Kurahashi Y, Nakatsu T, Miyao M (1987) Clinical Features and Cardiopulmonary Function of Patients with Atrophic Heart in Duchenne Muscular Dystrophy. Japanese Heart Journal 28:687–694

76. Sheth R, Galvan D, Lehrenbaum H, Cheeran D, Araj F, Amin A, Drazner MH, Zaha VG, Peshock RM, Mammen PP (2021) Abstract 11725: Low Left Ventricular Mass and Cardiomyopathy in Muscular Dystrophies: A Different Perspective on Potential Cardiac Mechanisms. Circulation 144:

77. Kessler KM, Pina I, Green B, Burnett B, Laighold M, Bilsker M, Palomo AR, Myerburg RJ (1986) Cardiovascular findings in quadriplegic and paraplegic patients and in normal subjects. American Journal of Cardiology 58:525–530

78. Pelliccia A, Caselli S, Sharma S, et al (2017) European Association of Preventive Cardiology (EAPC) and European Association of Cardiovascular Imaging (EACVI) joint position statement: recommendations for the indication and interpretation of cardiovascular imaging in the evaluation of the athlete’s heart. Eur Heart J 39:1949–1969

79. Bjerring AW, Landgraff HE, Leirstein S, et al (2018) Morphological changes and myocardial function assessed by traditional and novel echocardiographic methods in preadolescent athlete’s heart. Eur J Prev Cardiol 25:1000–1007

80. Rodriguez-López AM, Javier G, Carmen P, Esteban P, Luisa G-C, Tomas F, Josefa HM, Luis F (2022) Athlete Heart in Children and Young Athletes. Echocardiographic Findings in 331 Cases. Pediatr Cardiol 43:407–412

81. Petek BJ, Groezinger EY, Pedlar CR, Baggish AL (2022) Cardiac effects of detraining in athletes: A narrative review. Ann Phys Rehabilitation Medicine 65:101581

82. Hornby NL, Drees R, Harron R, Chang R, Wells DJ, Piercy RJ (2021) Musculoskeletal magnetic resonance imaging in the DE50-MD dog model of Duchenne muscular dystrophy. Neuromuscular Disord 31:736–751

83. Karimjee K, Olsen E, Piercy R, Daley M (2019) P.325Frequency characterisation of activity and behaviours in the deltaE50-MD dog model of Duchenne muscular dystrophy. Neuromuscular Disord 29:S162

84. Lonsdale RA, Labuc RH, Robertson ID (1998) Echocardiographic parameters in training compared with non-training greyhounds. Vet Radiol Ultrasoun 39:325–330

85. Vatne L, Dickson D, Tidholm A, Caivano D, Rishniw M (2021) The effects of activity, body weight, sex and age on echocardiographic values in English setter dogs. J Vet Cardiol 37:26–41

86. McKeever KH, Schurg WA, Convertino VA (1985) Exercise training-induced hypervolemia in greyhounds: role of water intake and renal mechanisms. Am J Physiology-regulatory Integr Comp Physiology 248:R422–R425

87. Mackintosh IC, Dormehl IC, Gelder AL van, Plessis M du (1983) Blood volume, heart rate, and left ventricular ejection fraction changes in dogs before and after exercise during endurance training. Am J Vet Res 44:1960–2

88. Ritzer TF, Bove AA, Carey RA (1980) Left ventricular performance characteristics in trained and sedentary dogs. J Appl Physiol 48:130–138

89. Wyatt HL, Mitchell JH (1974) Influences of Physical Training on the Heart of Dogs. Circ Res 35:883–889

90. Wilson DGS, Tinker A, Iskratsch T (2022) The role of the dystrophin glycoprotein complex in muscle cell mechanotransduction. Commun Biology 5:1022

91. Dweck MR, Joshi S, Murigu T, et al (2012) Left ventricular remodeling and hypertrophy in patients with aortic stenosis: insights from cardiovascular magnetic resonance. J Cardiov Magn Reson 14:50

92. Ganau A, Devereux RB, Roman MJ, Simone G de, Pickering TG, Saba PS, Vargiu P, Simongini I, Laragh JH (1992) Patterns of left ventricular hypertrophy and geometric remodeling in essential hypertension. J Am Coll Cardiol 19:1550–1558

93. Castro SD, Caselli S, Maron M, et al (2007) Left ventricular remodelling index (LVRI) in various pathophysiological conditions: a real-time three-dimensional echocardiographic study. Heart 93:205

94. Caselli S, Paolo FMD, Pisicchio C, Pietro RD, Quattrini FM, Giacinto BD, Culasso F, Pelliccia A (2011) Three-Dimensional Echocardiographic Characterization of Left Ventricular Remodeling in Olympic Athletes. Am J Cardiol 108:141–147

95. Baggish AL, Wang F, Weiner RB, Elinoff JM, Tournoux F, Boland A, Picard MH, Hutter AM, Wood MJ (2008) Training-specific changes in cardiac structure and function: a prospective and longitudinal assessment of competitive athletes. J Appl Physiol 104:1121– 1128

96. Binnetoğlu FK, Babaoğlu K, Altun G, Kayabey Ö (2014) Effects That Different Types of Sports Have on the Hearts of Children and Adolescents and the Value of Two-Dimensional Strain-Strain-Rate Echocardiography. Pediatr Cardiol 35:126–139

97. Morganroth J, Maron BJ, Henry WL, Epstein SE (1975) Comparative Left Ventricular Dimensions in Trained Athletes. Ann Intern Med 82:521

98. Robbers-Visser D, Boersma E, Helbing WA (2009) Normal biventricular function, volumes, and mass in children aged 8 to 17 years. J Magn Reson Imaging 29:552–559

99. Buechel EV, Kaiser T, Jackson C, Schmitz A, Kellenberger CJ (2009) Normal right- and left ventricular volumes and myocardial mass in children measured by steady state free precession cardiovascular magnetic resonance. Journal of Cardiovascular Magnetic Resonance 11:261–9

100. Nigro G, Comi LI, Politano L, Bain RJ (1990) The incidence and evolution of cardiomyopathy in Duchenne muscular dystrophy. Int J Cardiol 26:271–277

101. Stepien RL, Hinchcliff KW, Constable PD, Olson J (1998) Effect of endurance training on cardiac morphology in Alaskan sled dogs. J Appl Physiol 85:1368–1375

102. Markham LW, Michelfelder EC, Border WL, Khoury PR, Spicer RL, Wong BL, Benson DW, Cripe LH (2006) Abnormalities of Diastolic Function Precede Dilated Cardiomyopathy Associated with Duchenne Muscular Dystrophy. Journal of the American Society of Echocardiography 19:865–871

103. Mou YA, Lacampagne A, Irving T, Scheuermann V, Blot S, Ghaleh B, Tombe PP de, Cazorla O (2018) Altered myofilament structure and function in dogs with Duchenne muscular dystrophy cardiomyopathy. Journal of Molecular and Cellular Cardiology 114:345– 353

104. Barber BJ, Andrews JG, Lu Z, et al (2013) Oral corticosteroids and onset of cardiomyopathy in Duchenne muscular dystrophy. The Journal of Pediatrics 163:1080–4.e1

105. Jefferies JL, Eidem BW, Belmont JW, Craigen WJ, Ware SM, Fernbach SD, Neish SR, Smith EO, Towbin JA (2005) Genetic predictors and remodeling of dilated cardiomyopathy in muscular dystrophy. Circulation 112:2799–2804

106. James KA, Gralla J, Ridall LA, et al (2020) Left ventricular dysfunction in Duchenne muscular dystrophy. Cardiol Young 30:171–176

107. Spurney C, Shimizu R, Morgenroth LP, Kolski H, Gordish-Dressman H, Clemens PR, Investigators C (2014) Cooperative International Neuromuscular Research Group Duchenne Natural History Study demonstrates insufficient diagnosis and treatment of cardiomyopathy in Duchenne muscular dystrophy. Muscle Nerve 50:250–256

108. Moïse NS, Valentine BA, Brown CA, Erb HN, Beck KA, Cooper BJ, Gilmour RF (1991) Duchenne’s cardiomyopathy in a canine model: electrocardiographic and echocardiographic studies. J Am Coll Cardiol 17:812–820

109. Ghaleh B, Barthélemy I, Sambin L, Bizé A, Hittinger L, Blot S, Su JB (2020) Alteration in Left Ventricular Contractile Function Develops in Puppies With Duchenne Muscular Dystrophy. J Am Soc Echocardiog 33:120–129.e1

110. Yugeta N, Urasawa N, Fujii Y, et al (2006) Cardiac involvement in Beagle-based canine X-linked muscular dystrophy in Japan (CXMDJ): electrocardiographic, echocardiographic, and morphologic studies. BMC Cardiovasc Disord 6:47

111. Mertens L, Ganame J, Claus P, et al (2008) Early Regional Myocardial Dysfunction in Young Patients With Duchenne Muscular Dystrophy. Journal of the American Society of Echocardiography 21:1049–1054

112. Giatrakos N, Kinali M, Stephens D, Dawson D, Muntoni F, Nihoyannopoulos P (2006) Cardiac tissue velocities and strain rate in the early detection of myocardial dysfunction of asymptomatic boys with Duchenne’s muscular dystrophy: relationship to clinical outcome. Heart 92:840–842

113. Mori K, Edagawa T, Inoue M, Nii M, Nakagawa R, Takehara Y, Kuroda Y, Tatara K (2004) Peak negative myocardial velocity gradient and wall-thickening velocity during early diastole are noninvasive parameters of left ventricular diastolic function in patients with Duchenne’s progressive muscular dystrophy. Journal of the American Society of Echocardiography 17:322–329

114. Mori K, Hayabuchi Y, Inoue M, Suzuki M, Sakata M, Nakagawa R, Kagami S, Tatara K, Hirayama Y, Abe Y (2007) Myocardial Strain Imaging for Early Detection of Cardiac Involvement in Patients with Duchenne’s Progressive Muscular Dystrophy. Echocardiography 24:598–608

115. Shabanian R, Aboozari M, Kiani A, Seifirad S, Zamani G, Nahalimoghaddam A, Kocharian A (2011) Myocardial Performance Index and Atrial Ejection Force in Patients with Duchenne’s Muscular Dystrophy. Echocardiography 28:1088–1094

116. Bahler RC, Mohyuddin T, Finkelhor RS, Jacobs IB (2005) Contribution of Doppler Tissue Imaging and Myocardial Performance Index to Assessment of Left Ventricular Function in Patients with Duchenne’s Muscular Dystrophy. Journal of the American Society of Echocardiography 18:666–673

117. Li W, Liu W, Zhong J, Yu X (2009) Early manifestation of alteration in cardiac function in dystrophin deficient mdx mouse using 3D CMR tagging. Journal of Cardiovascular Magnetic Resonance 11:40–11

118. Miyazaki T, Tatara K, Mori K, Inoue M, Hayabuchi Y, Kagami S (2010) Increased mid-left ventricular rotation in patients with Duchenne muscular dystrophy using two-dimensional speckle tracking echocardiography. Journal of Echocardiography 8:14–24

119. Takano H, Fujii Y, Yugeta N, Takeda S, Wakao Y (2011) Assessment of left ventricular regional function in affected and carrier dogs with Duchenne muscular dystrophy using speckle tracking echocardiography. BMC Cardiovasc Disord 11:23

120. Ryan TD, Taylor MD, Mazur W, et al (2013) Abnormal circumferential strain is present in young Duchenne muscular dystrophy patients. Pediatric Cardiology 34:1159–1165

121. Hor KN, Wansapura J, Markham LW, Mazur W, Cripe LH, Fleck R, Benson DW, Gottliebson WM (2009) Circumferential Strain Analysis Identifies Strata of Cardiomyopathy in Duchenne Muscular Dystrophy. J Am Coll Cardiol 53:1204–1210

122. Hagenbuch SC, Gottliebson WM, Wansapura J, Mazur W, Fleck R, Benson DW, Hor KN (2010) Detection of Progressive Cardiac Dysfunction by Serial Evaluation of Circumferential Strain in Patients With Duchenne Muscular Dystrophy. Am J Cardiol 105:1451–1455

123. Spurney CF, McCaffrey FM, Cnaan A, et al (2015) Feasibility and Reproducibility of Echocardiographic Measures in Children with Muscular Dystrophies. J Am Soc Echocardiogr 28:999–1008

124. Taqatqa A, Bokowski J, Al-Kubaisi M, Khalil A, Miranda C, Alaksham H, Fughhi I, Kenny D, Diab KA (2016) The Use of Speckle Tracking Echocardiography for Early Detection of Myocardial Dysfunction in Patients with Duchenne Muscular Dystrophy. Pediatric Cardiology 37:1422–1428

125. Amedro P, Vincenti M, Villeon GDL, et al (2019) Speckle-Tracking Echocardiography in Children With Duchenne Muscular Dystrophy: A Prospective Multicenter Controlled Cross-Sectional Study. J Am Soc Echocardiogr 32:412–422

126. Oreto L, Vita GL, Mandraffino G, et al (2020) Impaired myocardial strain in early stage of Duchenne muscular dystrophy: its relation with age and motor performance. Acta Myologica 39:191–199

127. Opdahl A, Helle-Valle T, Remme EW, Vartdal T, Pettersen E, Lunde K, Edvardsen T, Smiseth OA (2008) Apical Rotation by Speckle Tracking Echocardiography: A Simplified Bedside Index of Left Ventricular Twist. J Am Soc Echocardiog 21:1121–1128

128. Bogaert J, Rademakers FE (2001) Regional nonuniformity of normal adult human left ventricle. Am J Physiol-heart C 280:H610–H620

129. Joseph S, Moazami N, Cupps BP, Howells A, Craddock H, Ewald G, Rogers J, Pasque MK (2009) Magnetic Resonance Imaging–based Multiparametric Systolic Strain Analysis and Regional Contractile Heterogeneity in Patients With Dilated Cardiomyopathy. J Hear Lung Transplant 28:388–394

130. Kar J, Knutsen AK, Cupps BP, Zhong X, Pasque MK (2015) Three-dimensional regional strain computation method with displacement encoding with stimulated echoes (DENSE) in non-ischemic, non-valvular dilated cardiomyopathy patients and healthy subjects validated by tagged MRI. J Magn Reson Imaging 41:386–396

131. Duan F, Xie M, Wang X, Li Y, He L, Jiang L, Fu Q (2012) Preliminary clinical study of left ventricular myocardial strain in patients with non-ischemic dilated cardiomyopathy by three-dimensional speckle tracking imaging. Cardiovasc Ultrasoun 10:8

132. Kusunose K, Zhang Y, Mazgalev TN, Thomas JD, Popović ZB (2013) Left ventricular strain distribution in healthy dogs and in dogs with tachycardia-induced dilated cardiomyopathy. Cardiovasc Ultrasoun 11:43

133. Pedro B, Stephenson H, Linney C, Cripps P, Dukes-McEwan J (2017) Assessment of left ventricular function in healthy Great Danes and in Great Danes with dilated cardiomyopathy using speckle tracking echocardiography. J Vet Cardiol 19:363–375

134. Støylen A, Mølmen HE, Dalen H (2019) Left ventricular global strains by linear measurements in three dimensions: interrelations and relations to age, gender and body size in the HUNT Study. Open Hear 6:e001050

135. Hor KN, Kissoon N, Mazur W, et al (2015) Regional circumferential strain is a biomarker for disease severity in duchenne muscular dystrophy heart disease: a cross-sectional study. Pediatric Cardiology 36:111–119

136. Siegel B, Olivieri L, Gordish-Dressman H, Spurney CF (2018) Myocardial Strain Using Cardiac MR Feature Tracking and Speckle Tracking Echocardiography in Duchenne Muscular Dystrophy Patients. Pediatr Cardiol 39:478–483

137. Rothstein ES, Palac RT, O’Rourke DJ, Venkataraman P, Gemignani AS, Friedman SE (2021) Evaluation of echocardiographic derived parameters for right ventricular size and function using cardiac magnetic resonance imaging. Echocardiogr 38:1336–1344

138. Agasthi P, Chao C, Siegel RJ, et al (2020) Comparison of echocardiographic parameters with cardiac magnetic resonance imaging in the assessment of right ventricular function. Echocardiogr 37:1792–1802

139. Mah K, Mertens L (2022) Echocardiographic Assessment of Right Ventricular Function in Paediatric Heart Disease: A Practical Clinical Approach. Cjc Pediatric Congenit Hear Dis 1:136–157

140. Sieslack AK, Dziallas P, Nolte I, Wefstaedt P, Hungerbühler SO (2014) Quantification of right ventricular volume in dogs: a comparative study between three-dimensional echocardiography and computed tomography with the reference method magnetic resonance imaging. BMC Vet Res 10:242

141. Fries RC, Gordon SG, Saunders AB, Miller MW, Hariu CD, Schaeffer DJ (2019) Quantitative assessment of two- and three-dimensional transthoracic and two-dimensional transesophageal echocardiography, computed tomography, and magnetic resonance imaging in normal canine hearts. J Vet Cardiol 21:79–92

142. Toaldo MB, Glaus T, Campagna I, Matos JN, Dennler M (2021) Echocardiographic assessment of right ventricular systolic function in healthy Beagle dogs compared to high field cardiac magnetic resonance imaging. Vet J 271:105653

143. Mehmood M, Hor KN, Al-Khalidi HR, et al (2015) Comparison of right and left ventricular function and size in Duchenne muscular dystrophy. Eur J Radiol 84:1938–1942

144. Mehmood M, Ambach SA, Taylor MD, et al (2016) Relationship of Right Ventricular Size and Function with Respiratory Status in Duchenne Muscular Dystrophy. Pediatric Cardiology 37:878–883

145. Dual SA, Maforo NG, McElhinney DB, Prosper A, Wu HH, Maskatia S, Renella P, Halnon N, Ennis DB (2021) Right Ventricular Function and T1-Mapping in Boys With Duchenne Muscular Dystrophy. J Magn Reson Imaging 54:1503–1513

146. Brown NK, Berhane H, Gambetta K, Markl M, Rigsby CK, Robinson JD, Husain N (2022) Right Ventricular Remodeling Assessed by MRI in Duchenne Muscular Dystrophy. J. Magn. Reson. Imaging

147. Caivano D, Dickson D, Pariaut R, Stillman M, Rishniw M (2018) Tricuspid annular plane systolic excursion-to-aortic ratio provides a bodyweight-independent measure of right ventricular systolic function in dogs. J Vet Cardiol 20:79–91

148. Koestenberger M, Ravekes W, Everett AD, Stueger HP, Heinzl B, Gamillscheg A, Cvirn G, Boysen A, Fandl A, Nagel B (2009) Right Ventricular Function in Infants, Children and Adolescents: Reference Values of the Tricuspid Annular Plane Systolic Excursion (TAPSE) in 640 Healthy Patients and Calculation of z Score Values. J Am Soc Echocardiog 22:715–719

149. Bosser G, Lucron H, Lethor J-P, Burger G, Beltramo F, Marie P-Y, Marçon F (2004) Evidence of early impairments in both right and left ventricular inotropic reserves in children with Duchenne’s muscular dystrophy. Am J Cardiol 93:724–727

150. Orlikowski D, Mansencal N, Nguyen SL, et al (2023) Prognosis of right ventricular systolic dysfunction in Duchenne muscular dystrophy patients. Archives Cardiovasc Dis Suppl 15:50

151. Yotsukura M, Miyagawa M, Tsua T, Ishihara T, Ishikawa K (2008) PULMONARY HYPERTENSION IN PROGRESSIVE MUSCULAR DYSTROPHY OF THE DUCHENNE TYPE. Jpn Circ J 52:321

152. Meyers TA, Townsend D (2015) Early right ventricular fibrosis and reduction in biventricular cardiac reserve in the dystrophin-deficient mdx heart. Am J Physiol-heart C 308:H303–H315

153. Barbin ICC, Pereira JA, Rovere MB, Moreira D de O, Marques MJ, Neto HS (2016) Diaphragm degeneration and cardiac structure in mdx mouse: potential clinical implications for Duchenne muscular dystrophy. J Anat 228:784–791

154. Stedman HH, Sweeney HL, Shrager JB, Maguire HC, Panettieri RA, Petrof B, Narusawa M, Leferovich JM, Sladky JT, Kelly AM (1991) The mdx mouse diaphragm reproduces the degenerative changes of Duchenne muscular dystrophy. Nature 352:536–539

155. Crisp A, Yin H, Goyenvalle A, et al (2011) Diaphragm rescue alone prevents heart dysfunction in dystrophic mice. Hum Mol Genet 20:413–421

156. Hornby (2021) Musculoskeletal Magnetic Resonance Imaging and complementary imaging techniques in the DE50-MD dog model of Duchenne Muscular Dystrophy. Doctoral thesis (Ph.D).

157. Thibaud J-L, Azzabou N, Barthelemy I, Fleury S, Cabrol L, Blot S, Carlier PG (2012) Comprehensive longitudinal characterization of canine muscular dystrophy by serial NMR imaging of GRMD dogs. Neuromuscul Disord 22 Suppl 2:S85–99

158. Lessa TB, Abreu DK de, Rodrigues MN, Brólio MP, Miglino MA, Ambrósio CE (2014) Morphological and ultrastructural evaluation of the golden retriever muscular dystrophy trachea, lungs, and diaphragm muscle. Microsc Res Tech 77:857–861

159. Yuasa K, Nakamura A, Hijikata T, Takeda S (2008) Dystrophin deficiency in canine X-linked muscular dystrophy in Japan (CXMDJ) alters myosin heavy chain expression profiles in the diaphragm more markedly than in the tibialis cranialis muscle. BMC Musculoskelet Disord 9:1–12

160. Caselli C, Cangemi G, Masotti S, Ragusa R, Gennai I, Ry SD, Prontera C, Clerico A (2016) Plasma cardiac troponin I concentrations in healthy neonates, children and adolescents measured with a high sensitive immunoassay method High sensitive troponin I in pediatric age. Clin Chim Acta 458:68–71

161. Hor KN, Taylor MD, Al-Khalidi HR, Cripe LH, Raman SV, Jefferies JL, O’Donnell R, Benson DW, Mazur W (2013) Prevalence and distribution of late gadolinium enhancement in a large population of patients with Duchenne muscular dystrophy: effect of age and left ventricular systolic function. Journal of Cardiovascular Magnetic Resonance 15:107

162. Tandon A, Villa CR, Hor KN, et al (2015) Myocardial Fibrosis Burden Predicts Left Ventricular Ejection Fraction and Is Associated With Age and Steroid Treatment Duration in Duchenne Muscular Dystrophy. J Am Hear Assoc Cardiovasc Cerebrovasc Dis 4:e001338

163. Aikawa T (2018) Progressive left ventricular dysfunction and myocardial fibrosis in Duchenne and Becker muscular dystrophy: a longitudinal cardiovascular magnetic resonance study. Pediatric Cardiology 0:0–0

164. Puchalski MD, Williams RV, Askovich B, Sower CT, Hor KH, Su JT, Pack N, Dibella E, Gottliebson WM (2009) Late gadolinium enhancement: precursor to cardiomyopathy in Duchenne muscular dystrophy? Int J Cardiovasc Imaging 25:57–63

165. Florian A, Ludwig A, Engelen M, Waltenberger J, Rösch S, Sechtem U, Yilmaz A (2014) Left ventricular systolic function and the pattern of late-gadolinium-enhancement independently and additively predict adverse cardiac events in muscular dystrophy patients. Journal of Cardiovascular Magnetic Resonance 16:81

166. Florian AR, Shomanova Z, Bietenbeck M, Chatzantonis G, Meier C, Yilmaz A (2020) Occurrence of Cardiovascular Events and Progression of Cardiomyopathy in Muscular Dystrophy Patients A CMR-Based Study. Jacc Cardiovasc Imaging 13:2258–2259

167. Menon SC, Etheridge SP, Liesemer KN, Williams RV, Bardsley T, Heywood MC, Puchalski MD (2014) Predictive Value of Myocardial Delayed Enhancement in Duchenne Muscular Dystrophy. Pediatric Cardiology 35:1279–1285

168. Bilchick KC, Salerno M, Plitt D, Dori Y, Crawford TO, Drachman D, Thompson WR (2011) Prevalence and distribution of regional scar in dysfunctional myocardial segments in Duchenne muscular dystrophy. Journal of Cardiovascular Magnetic Resonance 13:20

169. Hor KN, Wansapura JP, Al-Khalidi HR, et al (2011) Presence of mechanical dyssynchrony in Duchenne muscular dystrophy. Journal of Cardiovascular Magnetic Resonance 13:12

170. Malvestio LM, Martins IM, Moares FR, Moraes JRE (2015) Histopathologic Evolution of Cardiomyopathy in a Canine Model of Duchenne Muscular Dystrophy. Journal of Advanced Veterinary Research 5:1–6

171. Urasawa N, Wada MR, Machida N, et al (2008) Selective vacuolar degeneration in dystrophin-deficient canine Purkinje fibers despite preservation of dystrophin-associated proteins with overexpression of Dp71. Circulation 117:2437–2448

172. Echigoya Y, Nakamura A, Nagata T, et al (2017) Effects of systemic multiexon skipping with peptide-conjugated morpholinos in the heart of a dog model of Duchenne muscular dystrophy. Proc Natl Acad Sci USA 114:4213–4218

173. Valentine BA, Cooper BJ, Cummings JF, deLahunta A (1986) Progressive muscular dystrophy in a golden retriever dog: light microscope and ultrastructural features at 4 and 8 months. Acta Neuropathol 71:301–310

174. Valentine BA, Cummings JF, Cooper BJ (1989) Development of Duchenne-type cardiomyopathy. Morphologic studies in a canine model. Am J Pathol 135:671–678

175. Sato J, Doi T, Wako Y, Hamamura M, Kanno T, Tsuchitani M, Narama I (2012) Histopathology of Incidental Findings in Beagles Used in Toxicity Studies. J Toxicol Pathol 25:103–134

176. Shimatsu Y, Katagiri K, Furuta T, Nakura M, Tanioka Y, Yuasa K, Tomohiro M, Kornegay JN, Nonaka I, Takeda S (2003) Canine X-Linked Muscular Dystrophy in Japan (CXMD. Experimental Animals 52:1–5

