## Supplementary information for "The preclinical cardiac phenotype of the DE50-MD dog model of Duchenne muscular dystrophy"

| Dimension/ Volume | View | Description of technique | Timing in cardiac cycle | Method of normalisation |
|---|---|---|---|---|
| **LV volume** | RPx4ch | Estimated using Simpson's method of discs by tracing the endocardial border of the LV at end diastole and systole. | LV end diastolic volume (LVEDV): first frame after mitral valve closure<br>LV end systolic volume (LVESV): last frame prior to mitral valve opening | To calculated body surface area: LVEDVI and LVESVI |
| **LA diameter Short axis:** | RPxSA. Level of the heart base. | Measured from the junction of the left and non-coronary cusp of the aorta to the LA wall, using an inner edge to inner edge technique | At the end of the T wave, to coincide with the first frame after aortic valve closure | Indexed to aortic diameter measured in the same view: LA:Ao. |
| **LA diameter Long axis:** | RPx 4ch | Measured using a line drawn roughly parallel to the mitral annulus and an inner edge to inner edge technique. | The frame immediately prior to mitral valve opening (end LV systole) | According to body weight(BW)$^{0.324}$ |
| **Aortic diameter Short axis** | RPxSA at the level of the heart base | Measured measured along the commissure between the noncoronary and left coronary aortic valve cusps. | At the end of the T wave, to coincide with the first frame after aortic valve closure | |
| **Aortic diameter Long axis** | Left cranial view | Distance measured between the open aortic valve leaflets | At time of aortic valve leaflets opening, when clearly visualised | Early to mid -systole |
| **LV internal diameter And wall thicknesses** | | RPxSA views at the level of the chordae tendinae. M Mode recording, with the cursor bisecting the LV. Measured using a leading edge to leading edge technique. | LV end diastolic internal diameter (LVIDd), septal(IVS) and free wall (FW) diameters: at the onset of the QRS complex<br>LV end systolic internal diameter (LVIDs): at the nadir of septal excursion | 1. According to body weight using Esser et al allometric scaling method:<br>LVIDDN: LVIDD/BW $^{0.322}$<br>LVIDSN:LVIDS/BW$^{0.346}$<br>2. According to aortic diameter (AV diam)<br>LVIDd:Ao and LVIDs:Ao<br><br>Relative wall thickness: ((IVS+FW)/2)/LVID |
| **Tricuspid Annular Systolic plane excursion (TAPSE)** | | Left apical view, optimised for the RV. M-mode recordings with the cursor as parallel as possible to the majority of the RV free wall. Measurement of the maximal longitudinal displacement of the lateral tricuspid valve annulus toward the RV apex | Peak systole | According to bodyweight: TAPSE/ BW$^{0.297}$<br>2. According to aortic diameter (AV diam)<br>LVIDd:Ao and LVIDs:Ao |

**Supplementary Table 1: Table A: Detailed descriptions of two-dimensional diameter and volume measurements for transthoracic echocardiographic assessments.**

**Supplementary Table 1.B Detailed descriptions of pulsed wave spectral Doppler assessments of the WT and DE50-MD dogs**

| Flow recorded | View | Sample volume placement | Parameter recorded |
|---|---|---|---|
| **Mitral inflow** | Left apical 4ch | A 2-3mm sample volume placed at the open mitral valve leaflet tips, in alignment with blood flow. | Peak velocity of early diastolic filling: E wave velocity<br>Peak velocity of late (active) diastolic filling: A wave velocity |
| **Simultaneous recording: Aortic ejection and mitral inflow** | Left apical 5ch | A 5-6 mm sample volume placed between the LV outflow tract and mitral inflow | IVRT: Time from end of aortic ejection and beginning of LV early diastolic filling. |

| Dog ID | Genotype | Age at echocardiography assessment (months) | | | | | |
|---|---|---|---|---|---|---|---|
| | | 3 | 6 | 9 | 12 | 15 | 18 |
| WT-E3 | WT | | | | | | |
| WT-E5 | WT | | | | | | |
| WT-G2 | WT | | | | | | |
| WT-H4 | WT | | | | | | |
| WT-J1 | WT | | | | | | |
| WT-K4 | WT | | | | | | |
| WT-K5 | WT | | | | | | |
| WT-M2 | WT | | | | | | |
| WT-P3 | WT | | | | | | |
| WT-Q5 | WT | | | | | | |
| WT-T3 | WT | | | | | | |
| WT-T4 | WT | | | | | | |
| WT-Y2 | WT | | | | | | |
| WT-Y3 | WT | | | | | | |
| **n** | **WT** | **8** | **11** | **12** | **13** | **11** | **13** |

| Dog ID | Genotype | Age at echocardiography assessment (months) | | | | | |
|---|---|---|---|---|---|---|---|
| | | 3 | 6 | 9 | 12 | 15 | 18 |
| DE50-E4 | DE50-MD | | | | | | |
| DE50-G4 | DE50-MD | | | | | | |
| DE50-H6 | DE50-MD | | | | | | |
| DE50-I1 | DE50-MD | | | | | | |
| DE50-I2 | DE50-MD | | | | | | |
| DE50-K2 | DE50-MD | | | | | | |
| DE50-K3 | DE50-MD | | | | | | |
| DE50-L1 | DE50-MD | | | | | | |
| DE50-M1 | DE50-MD | | | | | | |
| DE50-P1 | DE50-MD | | | | | | |
| DE50-R4 | DE50-MD | | | | | | |
| DE50-T4 | DE50-MD | | | | | | |
| DE50-T6 | DE50-MD | | | | | | |
| DE50-U1 | DE50-MD | | | | | | |
| DE50-V2 | DE50-MD | | | | | | |
| DE50-Y1 | DE50-MD | | | | | | |
| DE50-Y5 | DE50-MD | | | | | | |
| **n** | **DE50-MD** | **12** | **13** | **11** | **11** | **9** | **9** |

| | | DE50-MD | Wild Type | |
|---|---|---|---|---|
| 3 | BW | 4.05 (3.10-5.80) | 5.08 (3.90-6.40) | |
| 6 | | 6.60 (5.30-9.00) | 9.23 (7.60-10.70) | |
| 9 | | 8.20 (6.20-11.20) | 10.70 (9.10-10.95) | <0.001 |
| 12 | | 8.50 (6.75-11.25) | 11.25 (9.45-12.40) | |
| 15 | | 8.70 (6.90-12.25) | 11.50 (9.10-12.15) | |
| 18 | | 8.65 (7.15-11.60) | 11.50 (9.60-12.5) | |
| AGE (p) | | <0.001 | | <0.001 |

**Supplementary Table 2: Summary of study population for echocardiography studies in wild type (WT) and DE50-MD dogs.** Highlighted cells represent dogs included in the study. Dogs that were euthanased prior to the planned 18 month end point are indicated by X. Inset shows results of linear mixed model analysis to explore the effect of genotype and age on body weight parameters in DE50-MD and WT dogs.

| Dog ID | Genotype | Age at MRI assessment (months) | | | | | |
|---|---|---|---|---|---|---|---|
| | | 3 | 6 | 9 | 12 | 15 | 18 |
| WT-E3 | WT | | | | | | |
| WT-E5 | WT | | | | | | |
| WT-G2 | WT | | | | | | |
| WT-H4 | WT | | | | | | |
| WT-J1 | WT | | | | | | |
| WT-K4 | WT | | | | | | |
| WT-K5 | WT | | | | | | |
| WT-M2 | WT | | | | | | |
| WT-P3 | WT | | | | | | |
| WT-Q5 | WT | | | | | | |
| WT-T3 | WT | | | | | | |
| WT-T4 | WT | | | | | | |
| WT-Y2 | WT | | | | | | |
| WT-Y3 | WT | | | | | | |
| n | **WT** | **10** | **10** | **12** | **14** | **11** | **13** |

| Dog ID | Genotype | Age at MRI assessment (months) | | | | | |
|---|---|---|---|---|---|---|---|
| | | 3 | 6 | 9 | 12 | 15 | 18 |
| DE50-E4 | DE50-MD | | | | | | |
| DE50-G4 | DE50-MD | | | | | | |
| DE50-H6 | DE50-MD | | | | | | |
| DE50-I1 | DE50-MD | | | | | | |
| DE50-I2 | DE50-MD | | | | | | |
| DE50-K2 | DE50-MD | | | | | | |
| DE50-K3 | DE50-MD | | | | | | |
| DE50-L1 | DE50-MD | | | | | | |
| DE50-M1 | DE50-MD | | | | | | |
| DE50-P1 | DE50-MD | | | | | | |
| DE50-R4 | DE50-MD | | | | | | |
| DE50-T4 | DE50-MD | | | | | | |
| DE50-T6 | DE50-MD | | | | | | |
| DE50-U1 | DE50-MD | | | | | | |
| DE50-V2 | DE50-MD | | | | | | |
| DE50-Y1 | DE50-MD | | | | | | |
| DE50-Y5 | DE50-MD | | | | | | |
| n | **DE50-MD** | **12** | **11** | **11** | **11** | **9** | **9** |

**Supplementary Table 3: Summary of study population for cardiac magnetic resonance imaging studies in wild type (WT) and DE50-MD dogs.** Highlighted cells represent dogs included in the study.

**LONGITUDINAL STRAIN**

AVERAGE PEAK STRAIN (%)

| | Peak Average LS (%) | | AGE | SSR ($s^{-1}$) | | AGE |
|---|---|---|---|---|---|---|
| Age (m) | DE50-MD | WT | (p) | DE50-MD | WT | (p) |
| 9 | -15.26 (-19.12,-12.89) | -16.11 (-18.9,-13.31) | | -1.80 (-2.30,-1.43) | -1.65 (-2.10,-1.10) | |
| 12 | -16.07 (-20.41,-12.62) | -15.7 (-24.83,-13.31) | 0.683 | -1.63 (-2.43,-1.47) | -1.57 (-2.67,-1.33) | 0.639 |
| 15 | -15.42 (-19.92,-14.31) | -16.56 (-18.76,-14.92) | | -1.65 (-2.00,-1.50) | -1.63 (-2.17,-1.30) | |
| 18 | -15.22 (-17.37,-13.23) | -18.07 (-18.83,-12.27) | | -1.52 (-1.87,-1.27) | -1.73 (-2.20,-1.37) | |
| Genotype (p) | 0.148 | | | 0.71 | | |

STRAIN RATE ($s^{-1}$)

| | ESR ($s^{-1}$) | | AGE | ASR ($s^{-1}$) | | AGE |
|---|---|---|---|---|---|---|
| Age (m) | DE50-MD | WT | (p) | DE50-MD | WT | (p) |
| 9 | 2.20 (1.53,2.93) | 2.22 (1.70,3.23) | | 1.27 (0.90,1.77) | 1.45 (0.93,1.67) | |
| 12 | 2.23 (1.90,3.10) | 2.23 (1.80,3.20) | 0.386 | 1.33 (0.83,2.30) | 1.50 (0.90,2.07) | |
| 15 | 2.09 (1.37,2.77) | 2.23 (1.63,2.87) | | 1.50 (1.00,1.69) | 1.60 (1.00,1.80) | |
| 18 | 2.02 (1.63,2.67) | 2.43 (2.00,3.03) | | 1.18 (0.83,1.90) | 1.20 (0.87,1.87) | 0.336 |
| GENOTYPE (p) | 0.25 | | | 0.942 | | |

**CIRCUMFERENTIAL STRAIN**

AVERAGE PEAK STRAIN (%)

| | Average Peak CS (%) | | AGE | SSR ($s^{-1}$) | | AGE |
|---|---|---|---|---|---|---|
| Age (m) | DE50-MD | WT | (p) | DE50-MD | WT | (p) |
| 9 | -23.46 (-27.89,-21.42) | -20.44 (-26.41,-17.49) | | -2.50 (-3.27,-1.97) | -2.05 (-3.13,-1.57) | |
| 12 | -22.92 (-25.03,-21.08) | -19.86 (-24.83,-16.16) | 0.49** | -2.32 (-2.80,-1.77) | -2.00 (-2.67,-1.53) | 0.418 |
| 15 | -24.64 (-28.18,-21.14) | -19.58 (-24.73,-17.66) | | -2.40 (-2.86,-1.90) | -1.93 (-2.70,-1.57) | |
| 18 | -24.56 (-26.50,-15.47) | -21.97 (-27.65,-17.43) | | -2.30 (-2.77,-1.67) | -2.25 (-3.50,-1.50) | |
| GENOTYPE (p) | 0.003** | | | 0.098 | | |

STRAIN RATE ($s^{-1}$)

| | ESR ($s^{-1}$) | | AGE | ASR ($s^{-1}$) | | AGE |
|---|---|---|---|---|---|---|
| Age (m) | DE50-MD | WT | (p) | DE50-MD | WT | (p) |
| 9 | 3.07 (2.50,4.17) | 2.40 (1.73,3.33) | | 1.47 (1.03,2.83) | 1.19 (0.83,1.80) | |
| 12 | 2.90 (2.37,3.70) | 2.35 (2.10,3.13) | 0.688 | 1.32 (1.10,1.90) | 1.25 (0.53,1.70) | 0.770* |
| 15 | 2.93 (2.37,3.37) | 2.20 (1.80,3.50) | | 1.47 (1.23,2.03) | 1.00 (0.67,1.77) | |
| 18 | 2.90 (1.87,3.33) | 2.80 (1.73,3.80) | | 1.42 (0.93,1.63) | 1.20 (0.73,2.25) | |
| GENOTYPE (p) | 0.006 | | | | | |

**Supplementary Table 4: Results of linear mixed model analysis to explore the effect of genotype and age on deformation parameters.** Tabulated values are expressed as median (minimum, maximum). Abbreviations: CS; Circumferential strain. LS; Longitudinal strain SSR; systolic strain rate. ESR; early diastolic strain rate. ASR; active diastolic strain rate; m; months. * Results of circumferential ASR were logarithmically transformed (natural log) prior to analysis in linear mixed models. ** A significant interaction between age and genotype was identified only for average peak circumferential strain (see text).

**Supplementary Table 5: Results of linear mixed model analysis, describing the overall effect of wall region (septum versus anterolateral wall) and age on regional peak longitudinal strain and strain rate within each genotype**. Tabulated values are expressed as median (minimum, maximum). Abbreviations: LS; Longitudinal strain. SSR; Longitudinal strain rate. There was no interaction between wall region and age in LMM analysis for either genotype. Septal wall strain values were more negative than those of the anterolateral wall in both groups of dogs

| | AGE (months) | DE50-MD | | | WT | | |
|---|---|---|---|---|---|---|---|
| LS (%) | | Septum | Anterolateral | Wall Region | Septum | Anterolateral | Wall Region |
| | 9 | -18.84 (-22.36,-15.52) | -14.72 (-18.28,-9.39) | **p<0.001** | -16.42 (-19.49,-11.35) | -18.28 (-22.96,-14.24) | **p<0.001** |
| | 12 | -18.28 (-22.76,-15.74) | -15.37 (-19.31,-10.82) | | -14.93 (-18.20,-11.35) | -18.28 (-23.76,-16.13) | |
| | 15 | -17.33 (-20.82,-16.16) | -14.04 (-15.74,-12.34) | | -15.79 (-18.74,-12.62) | -18.71 (-21.47,-16.82) | |
| | 18 | -17.55 (-21.21,-16.29) | -13.97 (-17.09,-12.31) | | -16.49 (-18.44,-11.30) | -21.30 (-22.67,-18.03) | |
| Age | | p=0.284 | | | p=0.222 | | |
| SSR ($s^{-1}$) | 9 | -2.12 (-2.64,-1.83) | -1.82 (-2.21,-1.29) | **p<0.001** | -2.12 (-2.72,-1.57) | -1.52 (-2.8,-1.13) | **p<0.001** |
| | 12 | -2.35 (-2.88,-1.69) | -1.67 (-2.48,-1.09) | | -2.13 (-2.84,-1.87) | -1.69 (-2.28,-1.13) | |
| | 15 | -2.23 (-2.65,-1.86) | -1.67 (-2.24,-1.22) | | -2.16 (-2.60,-1.78) | -1.45 (-2.28,-1.08) | |
| | 18 | -1.89 (-2.36,-1.71) | -1.49 (-1.85,-0.88) | | -2.21 (-2.72,-1.86) | -1.68 (-2.14,-1.37) | |
| | | p=0.152 | | | p=0.532 | | |

**Supplementary Table 6: Summary of study population for blood-borne cardiac biomarker studies in wild type (WT) and DE50-MD dogs**. Study populations are for A) cardiac troponin I (cTnI) and B) NTproBNP. Highlighted cells represent dogs included in the study.

A.

| Dog ID | Genotype | age at cTnI assessment | | |
|---|---|---|---|---|
| | | 6 | 12 | 18 |
| WT-Q5 | WT | | | |
| WT-T3 | WT | | | |
| WT-T7 | WT | | | |
| WT-Y2 | WT | | | |
| WT-Y3 | WT | | | |
| **n** | **WT** | **4** | **4** | **4** |

| Dog ID | Genotype | age at cTnI assessment | | |
|---|---|---|---|---|
| | | 6 | 12 | 18 |
| DE50-P1 | DE50-MD | | | |
| DE50-R4 | DE50-MD | | | |
| DE50-T4 | DE50-MD | | | |
| DE50-T6 | DE50-MD | | | |
| DE50-U1 | DE50-MD | | | |
| DE50-Y1 | DE50-MD | | | |
| DE50-Y5 | DE50-MD | | | |
| DE50-Z2 | DE50-MD | | | |
| DE50-Z3 | DE50-MD | | | |
| DE50-Z6 | DE50-MD | | | |
| **n** | **DE50-MD** | **8** | **8** | **10** |

B.

| Dog ID | Genotype | age at NT-proBNP assessment | | | | | |
|---|---|---|---|---|---|---|---|
| | | 3 | 6 | 9 | 12 | 15 | 18 |
| WT-E3 | WT | | | | | | |
| WT-E5 | WT | | | | | | |
| WT-G2 | WT | | | | | | |
| WT-H4 | WT | | | | | | |
| WT-J1 | WT | | | | | | |
| WT-K4 | WT | | | | | | |
| WT-K5 | WT | | | | | | |
| WT-M2 | WT | | | | | | |
| WT-P3 | WT | | | | | | |
| WT-Q5 | WT | | | | | | |
| WT-T3 | WT | | | | | | |
| WT-T7 | WT | | | | | | |
| WT-Y2 | WT | | | | | | |
| WT-Y3 | WT | | | | | | |
| **n** | **WT** | **12** | **11** | **14** | 13 | 14 | 13 |

| Dog ID | Genotype | age at NT-proBNP assessment | | | | | |
|---|---|---|---|---|---|---|---|
| | | 3 | 6 | 9 | 12 | 15 | 18 |
| DE50-E4 | DE50-MD | | | | | | |
| DE50-G4 | DE50-MD | | | | | | |
| DE50-H6 | DE50-MD | | | | | | |
| DE50-I1 | DE50-MD | | | | | | |
| DE50-I2 | DE50-MD | | | | | | |
| DE50-K2 | DE50-MD | | | | | | |
| DE50-K3 | DE50-MD | | | | | | |
| DE50-L1 | DE50-MD | | | | | | |
| DE50-L6 | DE50-MD | | | | | | |
| DE50-M1 | DE50-MD | | | | | | |
| DE50-P1 | DE50-MD | | | | | | |
| DE50-R4 | DE50-MD | | | | | | |
| DE50-T4 | DE50-MD | | | | | | |
| DE50-T6 | DE50-MD | | | | | | |
| DE50-U1 | DE50-MD | | | | | | |
| DE50-Y1 | DE50-MD | | | | | | |
| DE50-Y5 | DE50-MD | | | | | | |
| DE50-Z2 | DE50-MD | | | | | | |
| DE50-Z3 | DE50-MD | | | | | | |
| DE50-Z6 | DE50-MD | | | | | | |
| **n** | **WT** | **19** | **16** | **13** | 13 | 12 | 13 |

**Supplementary Information: Detailed descriptions of imaging protocols.**

**Colour tissue Doppler imaging**

Tissue Doppler imaging (TDI) was performed to evaluate LV myocardial velocities using a previously described method.[1, 2] Standard echocardiographic views were first obtained, and the grey-scale receiver gain was adjusted to optimise the clarity of the endocardial and epicardial boundaries of the LV free wall (the inferolateral wall). The sector width was narrowed to include only the segment of the LV myocardium under investigation and to achieve a frame rate of at least 150 frames per second (fps). Colour-coded tissue velocities were superimposed, adjusting the Doppler velocity range to avoid signal aliasing, and the Doppler receiver gain to optimise colouring of the myocardium. Standard right parasternal short axis echocardiographic views were first obtained at the level of the midventricle (papillary muscle level) with optimisation of the grey-scale receiver gain to maximise clarity of the endocardial and epicardial boundaries of the LV free wall (the inferolateral wall). The sector width was narrowed to include only the segment of the LV myocardium under investigation and to achieve a frame rate of at least 150 frames per second. Colour-coded tissue velocities were superimposed, adjusting the Doppler receiver gain and velocity range to optimise colouring and avoid signal aliasing.

**Speckle Tracking Echocardiography**

Strain parameters were derived using speckle tracking echocardiography (STE), every 3 months between 9 and 18 months of age. Studies were performed by the same observer using the same machine and transducer as for the conventional TTE assessment. All views intended for STE analysis were recorded at the highest frame rate possible, with a minimum acceptable rate of 80 frames per second. Echocardiographic loops were stored from the left apical 4 chamber view and right parasternal short axis view at the level of the LV papillary muscles to obtain longitudinal and circumferential parameters respectively.[3]The endocardial borders for each loop were manually traced during peak systole and the region of interest (ROI) was adjusted to include the entire width of the LV myocardium, which was then divided automatically into 6 segments of equal area by the software. Segments were named according to the American Society of Echocardiography segmental model for chamber quantification[4]. The tracking quality for each segment was evaluated both by the software and following visual assessment at low speed by the observer. Cardiac loops were only included if adequate tracking was confirmed using both methods and there was no evidence of any arrhythmia aside from sinus arrhythmia.[5] Where adequate tracking could not be achieved after 3 attempts, that loop was discarded and the next consecutive recorded loop was used.

**Cardiac magnetic resonance imaging**

Cardiac magnetic resonance imaging image acquisition was performed during transient apnoea at end expiration, induced following a brief period of controlled hyperventilation. Skeletal muscle imaging was also performed for the purpose of a separate study under the same general anaesthetic in some dogs[6] For those dogs, sagittal T1w turbo spin echo sequences were acquired of lumbar spine (to include the fifth lumbar vertebrae (L5)), in addition to T2w thin slice gradient echo images of both femurs. Two- and 4-chamber LV vertical long axis (VLA) cine images were prescribed from survey gradient echo transverse, sagittal and dorsal standard localising scans (scouts). Cine image acquisition was performed using a gradient echo pulse sequence with balanced steady-state free precession (bright-blood imaging). A series of short axis images of the LV was obtained using the 2- and 4-chamber VLA sequences. Care was taken to ensure that the image plane was parallel to the plane of the mitral annulus and that acquired images encompassed the entire LV and LA from the apex of the LV to the dorsal aspect of the LA. Contiguous 5mm slices were acquired, with an interslice gap of 0 mm and 30 frames per cardiac cycle.

Left ventricular mass, end-systolic and end-diastolic volumes were obtained using the disc summation method. Manual contours were drawn along the endocardial and epicardial borders from the end systolic and end-diastolic frames of the short axis stack. The end-diastolic image was defined as the frame with maximum dilatation and the end-systolic image as the frame with maximum contraction during the recorded cine loops. Papillary muscles were excluded but LV trabeculae were included in the ventricular lumen volume. The most basal slice included was the slice where the endocardial contours were interrupted by the LV outflow (even at end diastole) but greater than 50% of the ventricular lumen circumference was surrounded by a wall thickness consistent with ventricular myocardium. The LV mass was calculated both in systole and diastole by subtracting the endocardial volume from the epicardial volume and multiplying the result by the myocardial mass density (1.05g/ml).[7, 8] Where possible, contouring of the RV lumen was repeated in both the transverse and the short axis stack imaging planes. The equivalent frames to those selected for LV volumetric measurements were used for end-systolic and end-diastolic volume calculation, with the tricuspid and pulmonary valves used as the limits for right ventricular volume. Only endocardial contouring was performed, excluding the papillary muscles from RV blood volume calculation for consistency.

Left ventricular and RV stroke volumes were calculated by subtracting calculated end-systolic ventricular volumes from end-diastolic ventricular volumes. The EF for each ventricle (RVEF and LVEF) was calculated as follows: Ejection fraction = Stroke volume/End-diastolic volume* 100. Measurements for RV volume and EF were accepted if calculated RV stroke volumes from transverse and short axis imaging planes were within 10% agreement. Likewise, measurements for LV volumes were accepted if values for systolic and diastolic mass were within 10% agreement. Average values were reported for LV mass and for RV volumes.


1. Chetboul V, Carlos C, Blot S, Thibaud JL, Escriou C, Tissier R, Retortillo JL, Pouchelon J-L (2004) Tissue Doppler assessment of diastolic and systolic alterations of radial and longitudinal left ventricular motions in Golden Retrievers during the preclinical phase of cardiomyopathy associated with muscular dystrophy. Am J Vet Res 65:1335–1341

2. Chetboul V, Escriou C, Tessier D, Richard V, Pouchelon J-L, Thibault H, Lallemand F, Thuillez C, Blot S, Derumeaux G (2004) Tissue Doppler imaging detects early asymptomatic myocardial abnormalities in a dog model of Duchenne’s cardiomyopathy. Eur Heart J 25:1934–1939

3. Voigt J-U, Pedrizzetti G, Lysyansky P, et al (2015) Definitions for a Common Standard for 2D Speckle Tracking Echocardiography: Consensus Document of the EACVI/ASE/Industry Task Force to Standardize Deformation Imaging. Journal of the American Society of Echocardiography 28:183–193

4. Voigt J-U, Pedrizzetti G, Lysyansky P, et al (2015) Definitions for a common standard for 2D speckle tracking echocardiography: consensus document of the EACVI/ASE/Industry Task Force to standardize deformation imaging. European Hear J - Cardiovasc Imaging 16:1–11

5. Pedro B, Stephenson H, Linney C, Cripps P, Dukes-McEwan J (2017) Assessment of left ventricular function in healthy Great Danes and in Great Danes with dilated cardiomyopathy using speckle tracking echocardiography. J Vet Cardiol 19:363–375

6. Hornby NL, Drees R, Harron R, Chang R, Wells DJ, Piercy RJ (2021) Musculoskeletal magnetic resonance imaging in the DE50-MD dog model of Duchenne muscular dystrophy. Neuromuscular Disord 31:736–751

7. Schulz-Menger J, Bluemke DA, Bremerich J, et al (2013) Standardized image interpretation and post processing in cardiovascular magnetic resonance: Society for Cardiovascular Magnetic Resonance (SCMR) Board of Trustees Task Force on Standardized Post Processing. J Cardiov Magn Reson 15:35

8. Vinnakota KC, Bassingthwaighte JB (2004) Myocardial density and composition: a basis for calculating intracellular metabolite concentrations. Am J Physiol-heart C 286:H1742–H1749
